# SCOPE: a normalization and copy number estimation method for single-cell DNA sequencing

**DOI:** 10.1101/594267

**Authors:** Rujin Wang, Dan-Yu Lin, Yuchao Jiang

## Abstract

Whole genome single-cell DNA sequencing (scDNA-seq) enables characterization of copy number profiles at the cellular level. This technology circumvents the averaging effects associated with bulk-tissue sequencing and increases resolution while decreasing ambiguity in tracking the evolutionary history of cancer. ScDNA-seq data is, however, highly sparse and noisy due to the biases and artifacts that are introduced during the library preparation and sequencing procedure. Here, we propose SCOPE, a normalization and copy number estimation method for scDNA-seq data of cancer cells. The main features of SCOPE include: (i) a Poisson latent factor model for normalization, which borrows information across cells and regions to estimate bias, using negative control cells identified by cell-specific Gini coefficients; (ii) modeling of GC content bias using an expectation-maximization algorithm embedded in the normalization step, which accounts for the aberrant copy number changes that deviate from the null distributions; and (iii) a cross-sample segmentation procedure to identify breakpoints that are shared across cells from the same subclone. We evaluate SCOPE on a diverse set of scDNA-seq data in cancer genomics, using array-based calls of purified bulk samples as gold standards and whole-exome sequencing and single-cell RNA sequencing as orthogonal validations; we find that, compared to existing methods, SCOPE offers more accurate copy number estimates. Further, we demonstrate SCOPE on three recently released scDNA-seq datasets by 10X Genomics: we show that it can reliably recover 1% cancer cell spike-ins from a background of normal cells and that it successfully reconstructs cancer subclonal structure from ∼10,000 breast cancer cells.

## Introduction

Copy number variations (CNV) refer to duplications and deletions that lead to gains or losses of large segments of the chromosomes. CNVs are an abundant source of variations^1^ and have been associated with diseases^2^ such as HIV acquisition and progression,^3^ autism,^4^ schizophrenia,^5^ and systemic autoimmune diseases.^6,7^ In cancer, somatic CNVs, also referred to as copy number aberrations (CNAs), are prevailing, as shown by the Cancer Genome Atlas^8^ and the International Cancer Genome Consortium;^9^ these CNAs have been associated with cancer progression and metastases.^10,11^ It is, therefore, important to detect CNVs with high sensitivity and specificity.

In order to retain statistical power for association testing, CNV profiles across a large population of samples are required.^12^ Recent advances in next-generation sequencing enable genome-wide CNV detection in a high-throughput manner. However, biases and artifacts are introduced during the library preparation and sequencing step, making data normalization crucial for accurate CNV profiling. Several algorithms have been developed to remove experimental noise from GC content bias,^13^ sequencing depth,^14^ mappability,^15^ amplification efficiency,^16^ and other latent systematic artifacts.^17,18^ Yet profiling of somatic CNVs in cancer remains challenging due to the heterogeneous nature of the tumor samples—the observed copy number signals from bulk DNA sequencing (DNA-seq) are averaged and attenuated across multiple genotypically distinct cancer subpopulations.^19–21^ Tumor purity further dampens the CNV signals, and overall tumor ploidy leads to genome-wide gains or losses of chromosomal copies, all of which make statistical estimation and inference less tractable.^19,20^

Whole genome single-cell DNA-seq (scDNA-seq) enables the characterization of copy number profiles at the cellular level without the cell subpopulation confounding. This circumvents the averaging effects associated with bulk-tissue DNA-seq, increasing resolution while decreasing ambiguity in deconvolving cancer subclones and thus elucidating cancer evolutionary history (Supplementary Table S1).^22–24^ Conventional whole-genome amplification methods for scDNA-seq include degenerate oligonucleotide primed PCR (DOP-PCR), multiple displacement amplification (MDA), and multiple annealing and looping-based amplification cycles (MALBAC).^25^ MDA^26^ lacks uniformity, but it has a low sequencing error rate and is thus better suited for calling point mutations; DOP-PCR^27^ and MALBAC,^28^ on the other hand, have higher uniformity and have been successfully used for calling CNVs;^19–21^ see Supplementary Table S2 on details of the perspective amplification methods. Recently, 10X Genomics released the Single-Cell CNV Solution (https://www.10xgenomics.com/solutions/single-cell-cnv), which automatically uses microfluidic droplets to barcode cells and performs library construction. In the first step, the single cells are encapsulated into hydrogel cell beads and then lysed. This is followed by a second step of amplification and cell bead partitioning with 10X barcoded gel beads. The amplified and barcoded fragments are then pooled and converted to sequencing libraries. This significantly increases throughput and offers great potential for profiling CNVs across a large population of cells.

However, scDNA-seq data across all existing platforms and technologies is sparse, noisy, and highly variable, even within a homogeneous cell population. The extremely shallow and highly non-uniform depth of coverage, which is caused by non-linear amplification and significant dropout events during the library preparation and sequencing step,^25,29^ makes detecting CNVs by scDNA-seq challenging. Furthermore, the cancer genomes undergo large chromosome or chromosome-arm level deletions or duplications, as well as changes in cellular ploidy, both of which lead to recurrent and frequent CNAs that disrupt a large proportion of the genome across multiple samples.^30^ Existing methods for scDNA-seq either build an optimized normal/reference set for normalization^31^ or adopt a cell-specific normalization procedure for removing systematic biases (for example, those due to GC content), followed by a *post hoc* procedure for ploidy estimation and adjustment.^32^ These methods, which are adapted from those that are developed for bulk DNA-seq data, do not address the challenges and complexities caused by aberrant copy number changes or the complicating factor of tumor ploidy; they also cannot adequately and correctly remove the artifacts, for they make the invalid assumption that all read counts are sampled from diploid regions under the null when estimating the noise terms. The recurrent CNV signals can either be accidentally removed during the normalization step or bias the correction step.

To meet the widespread demand for CNV detection with single-cell resolution, we propose a new statistical and computational framework, SCOPE, for *<u>S</u>*ingle-cell *<u>COP</u>*y number *<u>E</u>*stimation in cancer. The distinguishing features of SCOPE include: (i) utilization of cell-specific Gini coefficients for quality control and for identification of normal/diploid cells, which are then used as negative control samples in a Poisson latent factor model for read depth normalization; (ii) modeling of GC content bias using an expectation-maximization (EM) algorithm embedded in the Poisson regression models to account for the discretized copy number states along the genome; and (iii) a cross-sample iterative segmentation procedure to identify breakpoints that are shared across cells with the same genetic background. We evaluate the performance of SCOPE on real scDNA-seq datasets from several cancer genomic studies, as summarized in Supplementary Table S1. Compared to existing methods, SCOPE is shown to more accurately estimate subclonal copy number aberrations and to have higher correlation with array-based copy number profiles of purified bulk samples from the same breast cancer patient. We show that the copy number profiles by scDNA-seq are also well recapitulated, although at low resolution, by whole-exome sequencing (WES) and single-cell RNA-sequencing (scRNA-seq). We finally demonstrate SCOPE on the recently released scDNA-seq data that was produced using the 10X Genomics single-cell CNV pipeline, showing that it can reliably recover 1% of the cancer cell spike-ins from a background of normal cells and successfully reconstruct cancer subclonal structure across 10,000 breast cancer cells. SCOPE is compiled as an open-source R package available at https://github.com/rujinwang/SCOPE.

## Material and Methods

### Methods overview

An overview of the SCOPE workflow is shown in Figure 1. SCOPE takes as input the mapped reads from assembled BAM files,^33^ which are pre-processed using the same bioinformatic pipeline (see Supplementary Note S1 for details). SCOPE then generates consecutive bins along the genome and computes the cell-by-bin read depth matrix, as well as the mappability and GC content for each bin. For data normalization, SCOPE adopts a Poisson latent factor model with an embedded EM algorithm to capture both cell- and bin-specific systemic biases and artifacts, as well as GC content bias. SCOPE also incorporates a cross-sample segmentation procedure, enabling shared breakpoints across cells from the same cancer subclone. It then outputs integer-valued copy numbers, allowing direct ploidy estimation without the need for *post hoc* adjustment.

**Figure 1.**
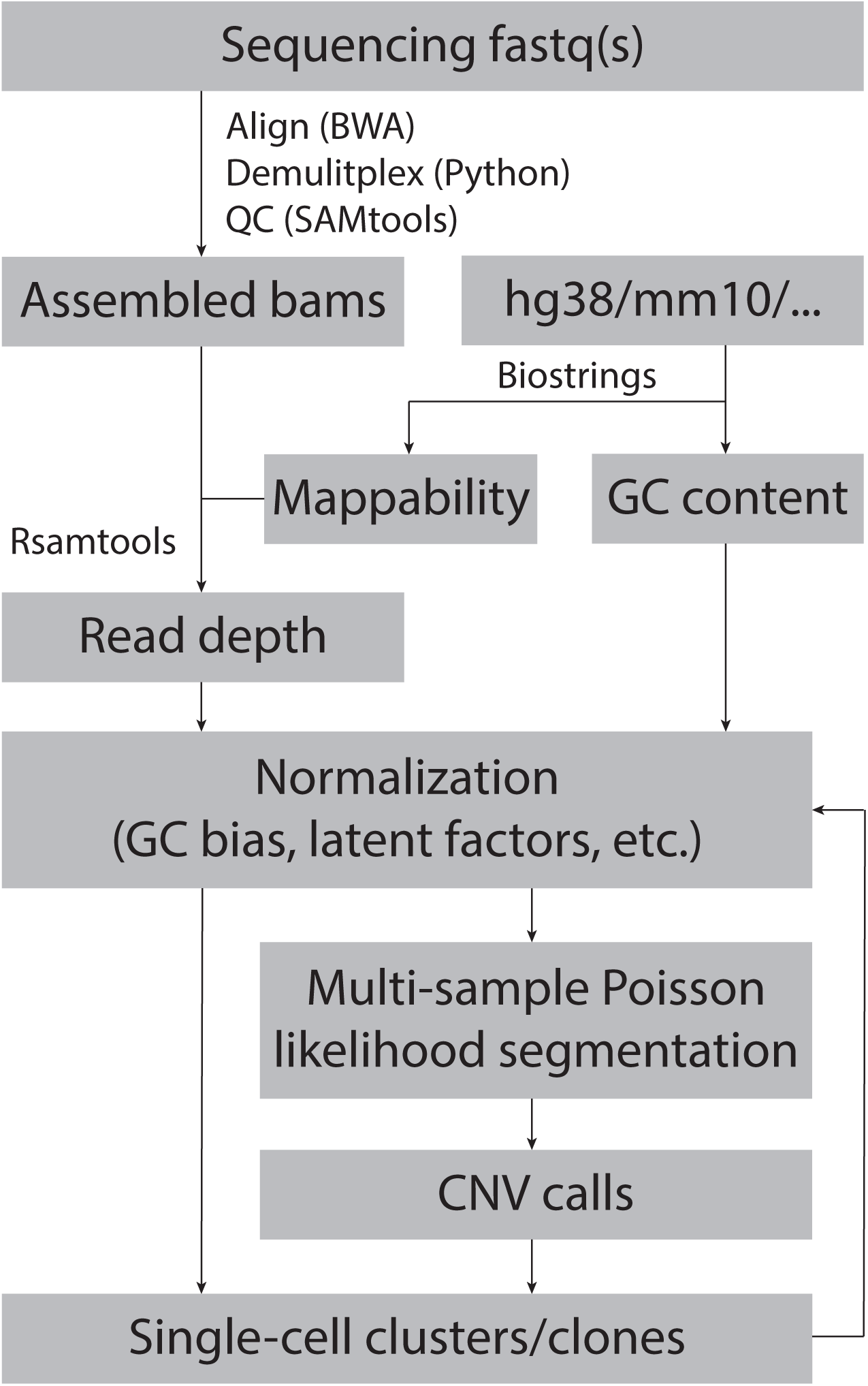
A flowchart outlining the procedures for profiling single-cell CNV by SCOPE. Bioinformatic pre-processing returns assembled BAM files as input for computing depth of coverage. Given the reference genome, mappability and GC content are calculated for each bin. SCOPE then adopts a Poisson latent factor model with an embedded EM algorithm for data normalization to remove biases due to sample-specific sequencing depth, bin-specific amplification/mapping efficiency, GC content, as well as other systemic latent artifacts. A cross-sample Poisson likelihood segmentation is performed to return copy number profiles, which are further used to infer cancer subclones.

In the following sections, we start with an overview of the *sample-specific* normalization procedure by Ginkgo,^32^ SCNV,^31^ and HMMcopy.^34^ We then discuss the limitations of the existing models and the reason that existing methods cannot adequately capture the biases and artifacts in scDNA-seq data, explaining why it is important to employ careful statistical modeling for *cross-sample* normalization with further adaptations specific to the single-cell setting to detect CNVs by scDNA-seq data. We then discuss the importance of data segmentation, offering an overview of existing methods and discussing why cross-sample segmentation is necessary. Finally, we describe the normalization model and the segmentation procedure for SCOPE, leaving algorithmic details to the supplements.

### Review of existing methods

Several methods have been developed for CNV detection by scDNA-seq, including Ginkgo^32^ and SCNV.^31^ To compute depth of coverage, both methods aggregate read-depth information into bins of variable lengths. To generate boundaries for the bins, *in silico* generated reads at every position of the genome are mapped back to the genome, and the bin sizes are chosen such that each bin has the same number of uniquely mapped reads. This step ensures that the bins have approximately the same number of mapped reads on average. To correct for GC content bias, both Ginkgo^32^ and SCNV^31^ perform normalization within each individual cell separately and employ a locally weighted linear regression (i.e., LOESS regression) to fit the relationship between GC content and log-normalized bin counts. Both methods assume that all reads are from the diploid regions, and both control for the GC biases by modeling the unimodal relationship between GC content and log-scaled read counts. This ignores the multiplicative effects contributed by the different sequential and discretized copy-number states in each single cell.

Furthermore, because depth of coverage within each cell is normalized against its own coverage baseline, each cell has a normalized value with mean zero and thus completely masks the cellular ploidy. To solve this issue, both Ginkgo and SCNV adopt a *post hoc* ploidy estimation procedure and further scale the normalized results by the estimated ploidy to reflect the true copy numbers. Specifically, they estimate the ploidy by minimizing a cost function that measures the difference between the scaled copy number (the normalized copy number multiplied by the ploidy) and its rounded copy number to the nearest integer. However, the copy numbers and the ploidy, the latter of which is the genome-wide average of the copy numbers, are interrelated—the normalized copy numbers depend on the true underlying ploidy, and the two-step approach estimates ploidy based on the normalization results. This leads to a cyclic chicken-and-egg problem. In the existing methods mentioned above, the recurrent CNV signals, as well as the complicating factors such as ploidy, can either be accidentally removed during the normalization step, requiring that they be recovered in a second step, or can bias the correction for known biases, such as those due to different GC content.

During the revision of this manuscript, we discovered another method, HMMcopy, which was initially developed for bulk whole-genome sequencing data^35^ but was recently applied to scDNA-seq data with adaptations.^34^ Instead of applying the default LOESS regression method to reduce GC content bias, the authors implemented a modal regression algorithm that normalizes bin counts to integer values, as expected of single-cell profiles. However, as we show later through benchmark analysis, the performance of the enhanced version of HMMcopy is data dependent and the method suffers from low stability—a finding that concords with another recent benchmark report^36^ on this method.

Thus far, we have focused on sample-specific normalization methods for scDNA-seq data. However, extremely low depth of sequencing coverage, nonlinear cell-specific biases due to amplification and sequencing, and other low-rank systemic artifacts that cannot be directly measured or quantified but are shared across cells and regions^37^ make the *sample-specific* bias correction procedure inadequate to successfully and unbiasedly capture all noise terms. To address these challenges, we instead adopt *cross-sample* normalization, which relies on multiple samples processed in the same experiment run and borrows information across both regions and samples to estimate the bias terms. This normalization strategy, based on matrix factorization, has been applied to different types of bulk omics data^16,18,38,39^ to adjust for GC content bias and other latent artifacts. Another refinement of this normalization strategy is found in RUV^38^ and CODEX2,^18^ which extensively utilize negative control samples and/or negative control regions/genes to estimate latent factors. For CNV detection, it has been shown that this increases CNV signal-to-noise ratio and achieves high sensitivity for both rare and common variants.^18^ The experiences we have gained by more than a decade of next-generation sequencing data analysis motivates many of the ingredients of our new proposed approach for single-cell DNA sequencing.

After proper data normalization, segmentation is performed to return regions with homogeneous copy number profiles. Existing methods adopt segmentation procedures based on either circular binary segmentation (CBS) or hidden Markov model (HMM),^11^ and they segment each cell separately.^31,32^ Specifically, Ginkgo^32^, which adopts the CBS^40^ algorithm, provides either independent segmentation, where cell-specific normalized read counts are segmented, or segmentation with reference samples, which calibrate segmentation against the cells with the most uniform coverage and the highest quality. SCNV^31^ relies on a set of normal cells to serve as the composite control and adapts the model proposed in Shen and Zhang^41^ designed for bulk DNA-seq for bin-free segmentation. However, all of these methods lack the ability to construct CNV profiles by integrating shared cellular breakpoints from the same genetic background. This is extremely important in single-cell settings, where multiple cells from the same subclone share the same genetic background and thus the same breakpoints for CNVs. Copynumber^42^ pools all cells together for joint segmentation to improve boundary detection accuracy. However, pooling across all cells forfeits the benefits of scDNA-seq and more importantly attenuates the signals of the breakpoints.

### SCOPE model for data normalization

SCOPE is based on a Poisson latent factor model^16,18^ for count-based read depth normalization. However, it is completely standalone and is specifically adapted for the single-cell setting. To estimate CNVs by bulk DNA-seq across subjects and to estimate CNVs by scDNA-seq across cells from the same subject are two different problems – in cancer genomics, the former profiles inter-tumor heterogeneity across patients, while the latter profiles intra-tumor heterogeneity looking at single cells within a patient. SCOPE’s key innovation is its integration of both null and non-null regions in a genome wide fashion in order to produce an unbiased estimation of both GC content bias and latent factors for cell- and position-specific background correction. The Poisson mixture model for normalization allows identification of discretized and integer-valued copy numbers, as expected of single-cell profiles. Unlike the two-step approach with post hoc ploidy adjustment, SCOPE offers direct ploidy estimates based on the estimated copy numbers along the genome, completely off the shelf.

Specifically, let *Y* = {*Y_ij_; i* = 1, … *m; j* = 1, … *n*} be the raw read count matrix, where *Y_ij_* is the read depth for cell *j* ∈ {1, …, *n*} and bin *i* ∈ {1, …, *m*}. SCOPE assumes that

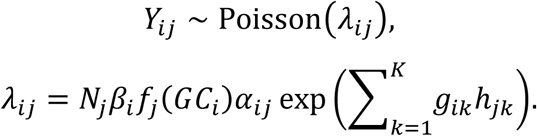

*N_j_* is a cell-specific library size factor, which can be globally estimated as the median ratio;^43^ *β_i_* reflects bias due to bin-specific length, capture and amplification efficiency; *f_j_*(*GC_i_*) is a sample-specific non-parametric function to capture the GC content bias, *g_ik_* and *h_jk_* (1 ≤ *k* ≤ *K*) are the *k*th bin- and cell-specific latent factors with orthogonality restraints, which force identifiability; and *α_ij_* specifies the sequential multiplicative increment due to different copy number states. *α_ij_* is discretized to fit the single-cell setting with integer-valued copy numbers and to ensure identifiability with *T_j_* different copy number states within a specific cell *j* (1 ≤ *j* ≤ *n*):

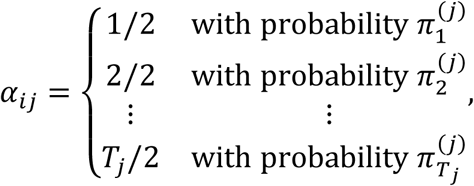

Where 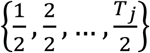 corresponds to the linear and discretized increments in read depths across all copy number states in cell *j*, with corresponding incidence rates 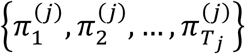 and 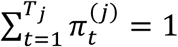. We denote 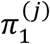as the incident rate for heterozygous deletion, 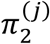 as the probability of a bin residing in a null region, and 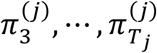 as the incident rates for amplifications in cell *j*. The optimal number of copy number groups *T_j_* is determined by the Bayesian information criterion (BIC) for within each cell separately (Supplementary Figure S1). By introducing the *α_ij_* term, we aim to unmask biases based on a mixture of Poisson distributions with observed-data likelihood:

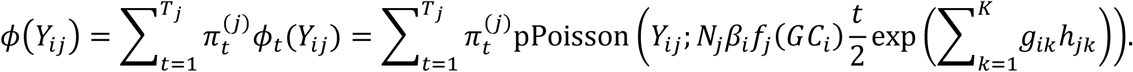

For each individual cell *j*, we adopt an EM algorithm within the iterative estimation procedure, where the missing data indicates carrier status:

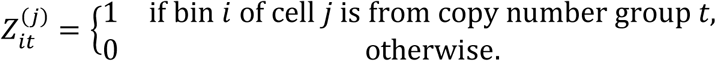

Given 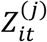, we have 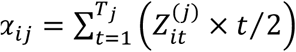.

For parameter estimation, SCOPE adopts an iterative procedure, where EM is embedded as one step to estimate the GC content bias. For EM initializations, SCOPE utilizes ploidy estimates from a first-pass normalization run to ensure fast convergence and to avoid local optima (Supplementary Figure S2). See algorithmic details in the Supplementary Note S2. For estimation of *k* (the number of latent factors), SCOPE by default adopts as a model selection metric another BIC based on normalization results across all cells and regions. Notably, we demonstrate that with the EM algorithm to account for the local genomic contexts and the use of normal cells to estimate bin-specific noise terms, the procedure by SCOPE (outlined above) is robust to the different choice of *k* (see under “Performance assessment via spike-in studies and with varying parameters” for more details).

For illustration, Figure 2 shows fitting of GC content bias in two ways: (i) assuming all bins are from the null region and fitting a non-parametric function using read depths across all bins—the method that, to our best knowledge, is used by all existing methods, and (ii) adopting an EM algorithm with the missing data being the carrier status for each bin, which is a simplified implementation of SCOPE. Three cells are chosen as examples, one diploid, one hypodiploid, and one hyperdiploid. For the diploid cell, the fitting by hyperdiploid cells, however, sequential multiplicative increments in *f*(*GC*)—the GC SCOPE is the same as that by the all-null fitting, as expected. For hypodiploid and — content biases—are observed due to prominent copy number changes along the genome. The “all-null” fit is biased by the global genomics structures due to deletions and duplications, but with the embedded EM algorithm, SCOPE is able to successfully identify the true null regions and correctly estimate the GC bias term (Figure 2, Supplementary Figure S1).

**Figure 2.**
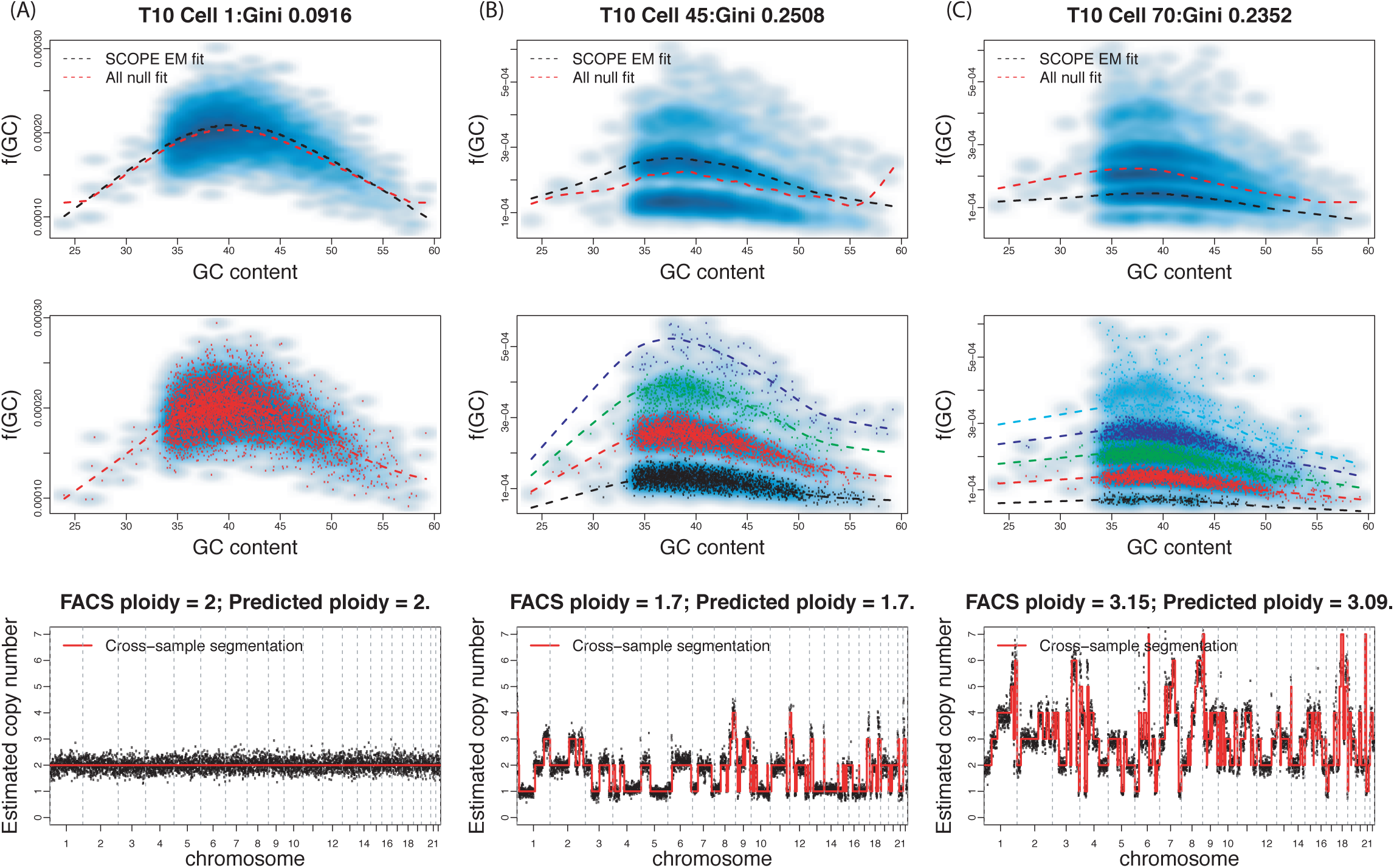
EM algorithm for correction of GC content bias. GC content bias *f*(*GC*), EM fitting, and cross-sample segmentation results of scDNA-seq data from breast cancer patient T10 are shown across three cells: (A) a diploid cell; (B) a hypodiploid cell with four copy number states; (C) a hyperdiploid cell with five copy number states. Predicted ploidy, taken as genome-wide average, agrees with previous FACS sorting results.

The histograms in Figure 3 show the normalization results for a breast cancer dataset across four methods: CODEX2, Ginkgo, Ginkgo with *post hoc* adjustment by the estimated ploidy, and SCOPE. We observe that neither CODEX2 nor Ginkgo attains unbiased separation of different copy number states. To solve this issue and to obtain integer-valued copy number estimates, Ginkgo^32^ adopts a *post hoc* ploidy estimation procedure, followed by a cell-specific scaling step; these ensure that the average copy number across all bins matches the estimated ploidy, an adjustment that allows Ginkgo to achieve better separation of CNV signals. SCOPE, on the other hand, retains precise segregation of CNV signals from the biological and technical noise, and distributions of the normalized score are centered at the expected integer copy-number values.

**Figure 3.**
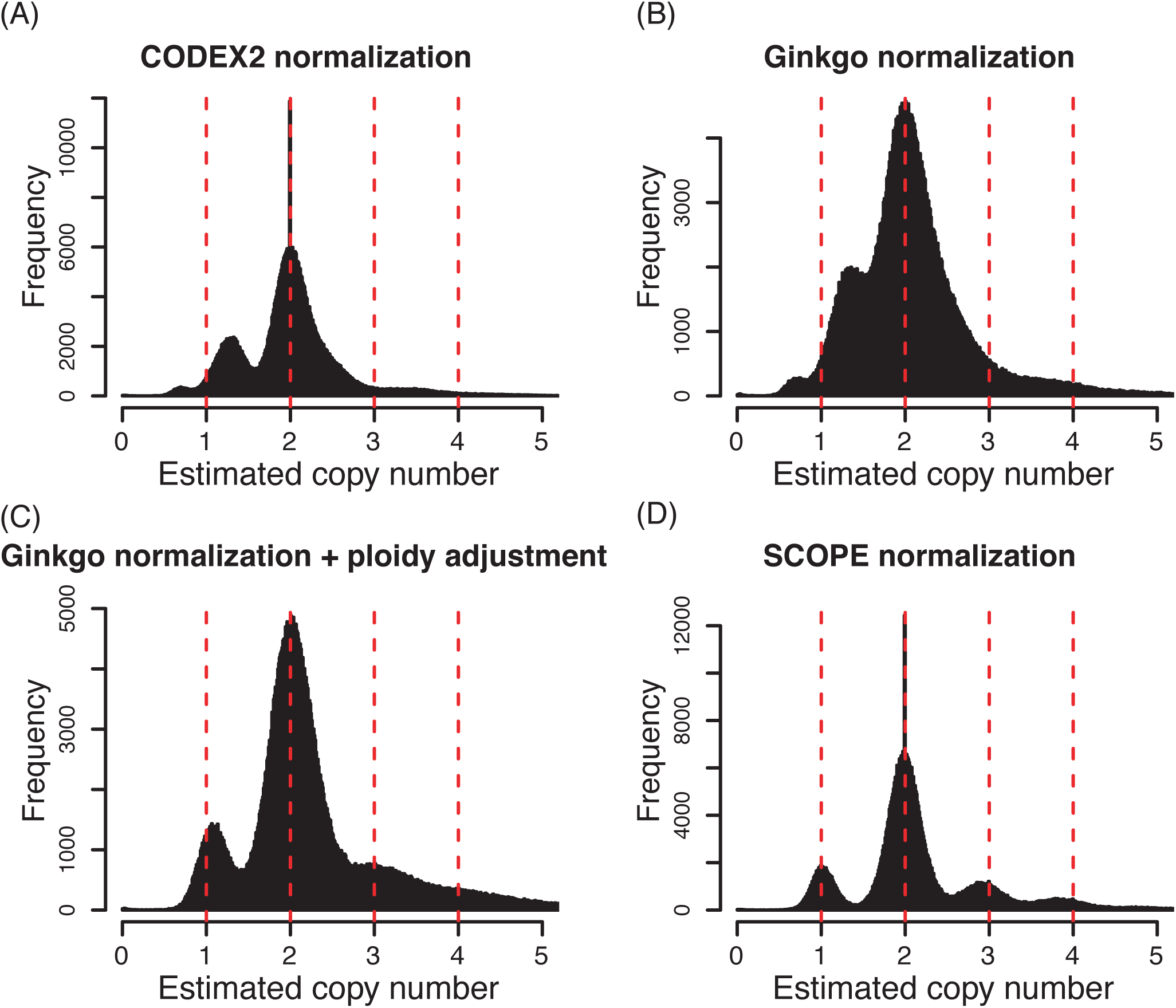
Normalization results for scDNA-seq data of ploygenomic tumor T10.^22^ Normalized *z*-scores are shown for (A) CODEX2 as 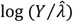, (B) Ginkgo, which centers bin counts of each cell at one and further fits a locally weighted linear regression (LOESS curve) to adjust for GC content bias, (C) Ginkgo with *post hoc* ploidy adjustment, which scales relative copy numbers by the optimal ploidy multiplier, and (D) SCOPE as 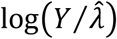, which automatically estimates discretized copy numbers and returns ploidy.

### Identification of negative control cells

In most, if not all, single-cell cancer genomics studies, diploid cells are inevitably picked up for sequencing from adjacent normal tissues, and they can thus serve as normal controls for read depth normalization. However, not all platforms/experiments allow or adopt flow-sorting based techniques before scDNA-seq, and cell ploidy and case-control labeling are therefore not always readily available. To solve this issue, SCOPE opts to use the scDNA-seq data to *in silico* identify normal cells as controls. Specifically, for each cell, we calculate its Gini coefficient as two times the area between the diagonal line and of cell j n be calculated as

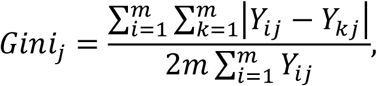

which serves as a robust and scale-independent measurement of coverage uniformity. As such, cell-specific Gini coefficients can be used as good proxies for identifying cell outliers, which have extreme coverage distribution due to failed library preparation, and for indexing normal cells out of the entire cell population.

The utility of Gini coefficients is supported by empirical evidence showing that diploid, hyperdiploid, and hypodiploid cells, categorized by single-cell flow sorting, can be perfectly classified using the estimated Gini indices (Supplementary Figure S4). This suggests that the Gini coefficient is an effective metric, with which to identify negative control samples. In practice, SCOPE does not require identification of all diploid cells from the cell population, only requiring 10 to 20 cells to serve as normal controls (refer to the section of “Performance assessment via spike-in studies and with varying parameters” on choosing the Gini coefficient threshold). The normal cells so identified are then used in the in the cross-sample normalization step as normal controls to estimate bin-specific noise terms {*β_i_*, *g*_*i*1_, …, *g_iK_* | 1 ≤ i ≤ *m*} that are not biased by the CNV signals. Refer to Supplementary Note S2 for details.

### Detecting simultaneous changepoints across samples

For segmentation, SCOPE adopts a Poisson likelihood-based recursive segmentation same subclone. Specifically, for cell *j* let *Y_sj_*, …, *Y_tj_* and 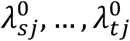 be the observed and normalized read depth from a region spanning bin *s* to bin *t*, where 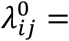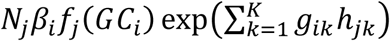 is the “control” depth of coverage under the null, i.e., the coverage we expect to see if there is no CNV. We further denote 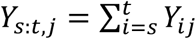 and 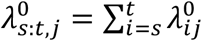. The scan statistic for one cell is 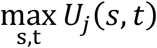, where *U_j_*(*s,t*) is calculated from three sub-splits:

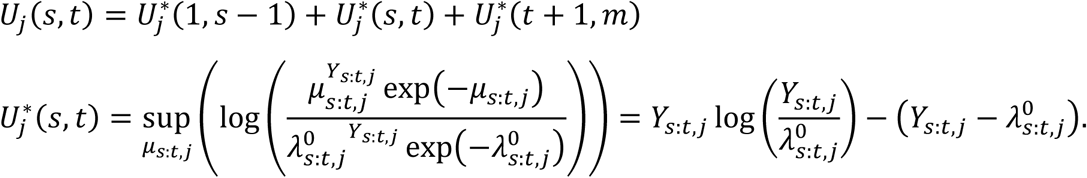

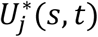 is a generalized likelihood ratio test statistic, derived from the null model *Y_s:t,j_* ~ Poisson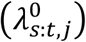 against the alternative *Y_s:t,j_* ~ Poisson(*μ_s:t,j_*). ^^44^^ For simultaneous changepoint detection across all cells, the scan statistic is 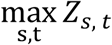 where

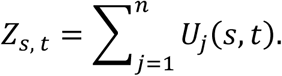

SCOPE performs CBS^40^ given *Z_s,t_* as the scan statistic, and uses a cross-sample modified BIC (mBIC)^45^ as the stopping rule:

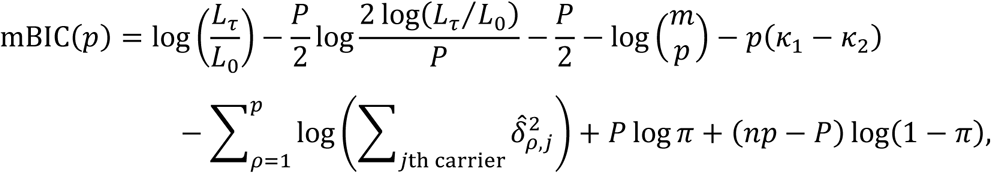

where 1 = *τ*_0_ < *τ*_1_ < *τ*_2_ < ⋯ < *τ_p_* < *τ*_*p*+1_ = *m* denote the change points that are shared log(*L_τ_*/*L*_0_) is the generalized log-likelihood ratio for the alternative model, which has *p* changepoints, against the null model, which has no changepoints; *P* denotes the total number of shift parameters (*δ*), where 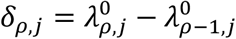 for cell *j* with changepoint *ρ* (1 ≤ *ρ* ≤ *p*); and *k*_1_ and *k*_2_ are pre-defined numerical constants. When the carrier probability *π* is not known a priori, it is estimated empirically by

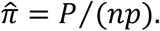

The optimal number of changepoints is determined via max*_p_* mBIC(*p*). The mBIC for regularizing the segmentation considers the number of carriers for each breakpoint, thereby accommodating subclonal events. For more details on the interpretation of the terms in mBIC, see Zhang and Siegmund.^45^

After segmentation, SCOPE reports integer-valued copy numbers and allows direct ploidy estimation, which is calculated as the weighted average of the estimated copy number across the genome. The lower panels in Figure 2 show SCOPE’s cross-sample segmentation results across three cells; the ploidy predicted by SCOPE is in concordance with that from single-cell flow sorting. For further downstream analysis, SCOPE also includes the option to cluster cells based on the matrix of normalized - scores, estimated copy numbers, or estimated changepoints, a process that returns clusters of cells from the same subclones with the same mutational profiles. As a final step, cells from the same subclone can be combined together to generate pseudo-bulk samples that are presumably homogeneous. Another normalization on the *in silico* generated pseudo-bulk samples returns copy number profiles with higher resolution. More details are described in the Methods section. The pseudo-bulk samples also have the potential for somatic point mutation profiling, with the sequencing sparsity averaged out. Refer to the Discussion section for more details.

### Availability of data and material

SCOPE is an open-source R package available at https://github.com/rujinwang/SCOPE, which will be released in Bioconductor (currently under review). Ginkgo was downloaded from https://github.com/robertaboukhalil/ginkgo. HMMcopy was downloaded from https://github.com/shahcompbio/single_cell_pipeline/tree/master/single_cell/workflows/hmmcopy. Breast cancer scDNA-seq data were downloaded from the NCBI Sequence Read Archive with accession number SRA018951 and SRP114962; breast cancer aCGH benchmark data were downloaded from the NCBI Gene Expression Omnibus with accession number GSE16607. ScDNA-seq data from 10X Genomics were downloaded from https://support.10xgenomics.com/single-cell-dna/datasets. A detailed data summary is provided in Supplementary Table S1.

## Results

We first set out to compare the biases and differences in depth of coverage and coverage uniformity between the conventional whole-genome amplification methods and the 10X Genomics pipeline. Compared to protocols that use conventional methods for cell isolation and whole-genome amplification (WGA), 10X Genomics produces data with both higher throughput (by the number of cells captured and sequenced) and significantly lower sequencing depth (Supplementary Table S1). Conventional WGA data achieves a mean total read count of 5.3 million per cell, while 10X Genomics data achieves a mean total read count of 0.4 million per cell (Supplementary Figure S5A). To assess coverage uniformity, we compared the cell-specific median absolute deviation (MAD) of all pairwise differences in read counts between neighboring bins, with a default bin size of 500kb for all platforms and datasets (refer to the section of “Performance assessment via spike-in studies and with varying parameters” on choice of bin size). We pre-normalized the read count data by a cell-specific factor for library size difference and a bin-specific factor for baseline coverage difference, then calculated the MADs across all cells; the MADs are resilient to outliers, as the transitions between copy number states are relatively infrequent along the genome. Strikingly, our results show that even though cells from the 10X Genomics pipeline are sequenced at a much shallower depth, the coverage uniformity metrics show trivial differences between data from the 10X Genomics and the conventional methods, except that the former has a higher number of cell outliers (Supplementary Figure S5B).

### Analysis of scDNA-seq data of breast cancer patients with aCGH for validation

We first demonstrated SCOPE on the scDNA-seq data of two breast cancer patients, T10 and T16, from Navin et al.,^22^ who sequenced 100 flow-sorted single cells from each patient. Fluorescence activated cell sorting (FACS) analysis of the single cells showed different distributions of ploidy: hyperdiploid, hypodiploid, and diploid.^22^ Array comparative genomic hybridization (aCGH) was applied to the same tumor sectors from both patients, and this returned array-based copy number profiles of flow-sorted cell populations.^46^ Cell ploidy by FACS, copy number profiles of purified bulk samples by aCGH, and initially reported copy number profiles of single cells estimated by scDNA-seq all suggested that patient T10 had three distinct clonal subpopulations, indicating a polygenomic tumor. In patient T16, a relatively homogeneous cancer cell population was identified in both primary tumor and metastasis, indicating a monogenomic tumor.^22^ We sought to apply SCOPE to this dataset to replicate previous findings, and further benchmarked the method by using copy number calls by aCGH as orthogonal validations.

Figure 4 gives heatmaps of genome-wide estimated copy numbers across all cells from T10 and T16. For T10, SCOPE identified two subpopulations of hyperdiploid cancer cells, one subpopulation of hypodiploid cancer cells, and a normal cell subpopulation, which is consistent with the previous report. For T16, SCOPE returned two cancer cell subclones, one for the hyperdiploid subpopulation from the primary tumor and the other for the hyperdiploid subpopulation from the metastasis. Upon careful inspection of the consensus copy number profiles of these two subpopulations, we find them highly similar, indicating that the same subclone of origin from the primary tumor survived the selection bottleneck and led to relapse. Notably, SCOPE also identified three pseudodiploid cells (red annotation bars in Figure 4B), in concordance with the previous report. Overall, we showed that SCOPE was able to reproduce subclonal structures that are consistent with previous results. The normalization results by SCOPE better separate CNV signals from noise than other existing methods, as can be seen from the histograms of the normalized z-scores shown in Figure 3 and Supplementary Figure S6 (for T10 and T16, respectively).

**Figure 4.**
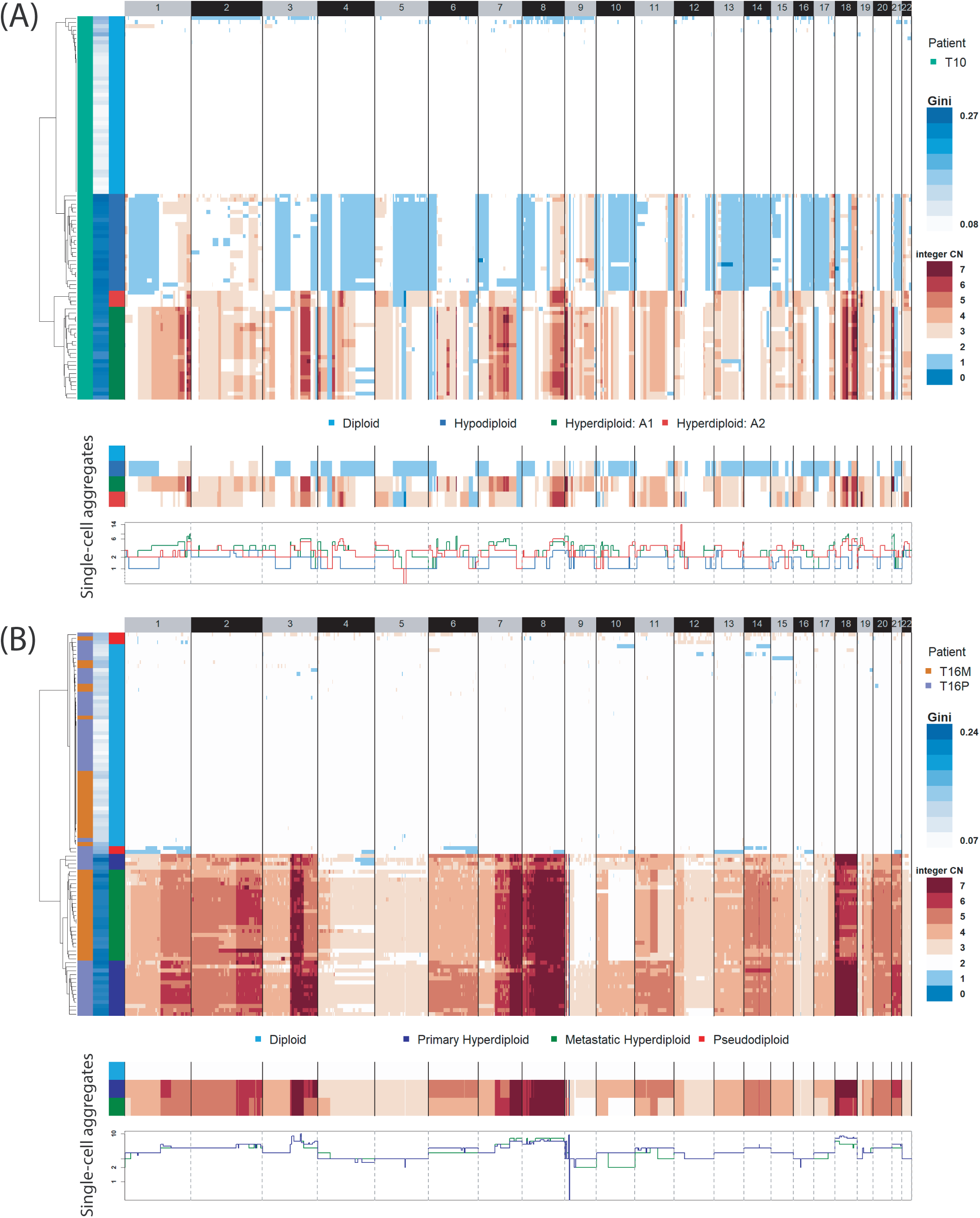
SCOPE successfully detects subclonal structures for triple negative breast cancer samples. (A) Inferred copy number profiles of single cells from polygenomic tumor T10. (B) Inferred copy number profiles of single cells from monogenomic tumor T16. Upper heatmaps show post-segmentation copy number profiles of integer values. Hierarchical clustering separates normal cells from tumor cells and reveals three cancer subclones in T10 and two highly similar subclones in T16. Lower heatmaps and stairstep plots show consensus copy-number profiles of all single cells from the same cancer subclones.

If we focus only on normal cells, there is an even clearer separation of normalized copy numbers, as shown by the disappearance of the spike at two from the normal cells (Supplementary Figure S7). In addition, the ploidy estimates by SCOPE are highly concordant with those from previous reports based on single-cell flow sorting (Supplementary Figure S8).

To further assess the performance of SCOPE and to benchmark against existing methods, we adopted CNV calls from aCGH of purified bulk samples^46^ from the same patient as gold standards (Supplementary Table S3). FACS revealed three cancer subpopulations (aneuploid 1, aneuploid 2, and hypodiploid), which is, again, in concordance with SCOPE’s results. This was followed by aCGH for copy number profiling of the purified bulk samples. The relative copy numbers (i.e., absolute copy numbers divided by overall ploidy) from both aCGH (taken from Navin et al.^46^) and scDNA-seq (produced by SCOPE, Ginkgo, and HMMcopy) are shown in Supplementary Figure S9 across the three cancer subpopulations: for aCGH, the subpopulations were inferred by the initial step of flow sorting; for SCOPE, Ginkgo and HMMcopy, the subpopulations were inferred and aggregated based on the estimated single-cell copy number profiles. Notably, array-based intensity measurements were normalized against a sample-specific baseline, producing copy number signals that have a mean of one for all three subpopulations. Therefore, relative copy numbers (shown as horizontal solid lines in Supplementary Figure S9 for diploid regions) were used by SCOPE, Ginkgo, and HMMcopy to mask ploidy for comparison against aCGH calls. Using spearman correlation and root mean squared error (RMSE) as performance metrics, SCOPE outperforms the other two methods by the aCGH gold standards (Figure 5).

**Figure 5.**
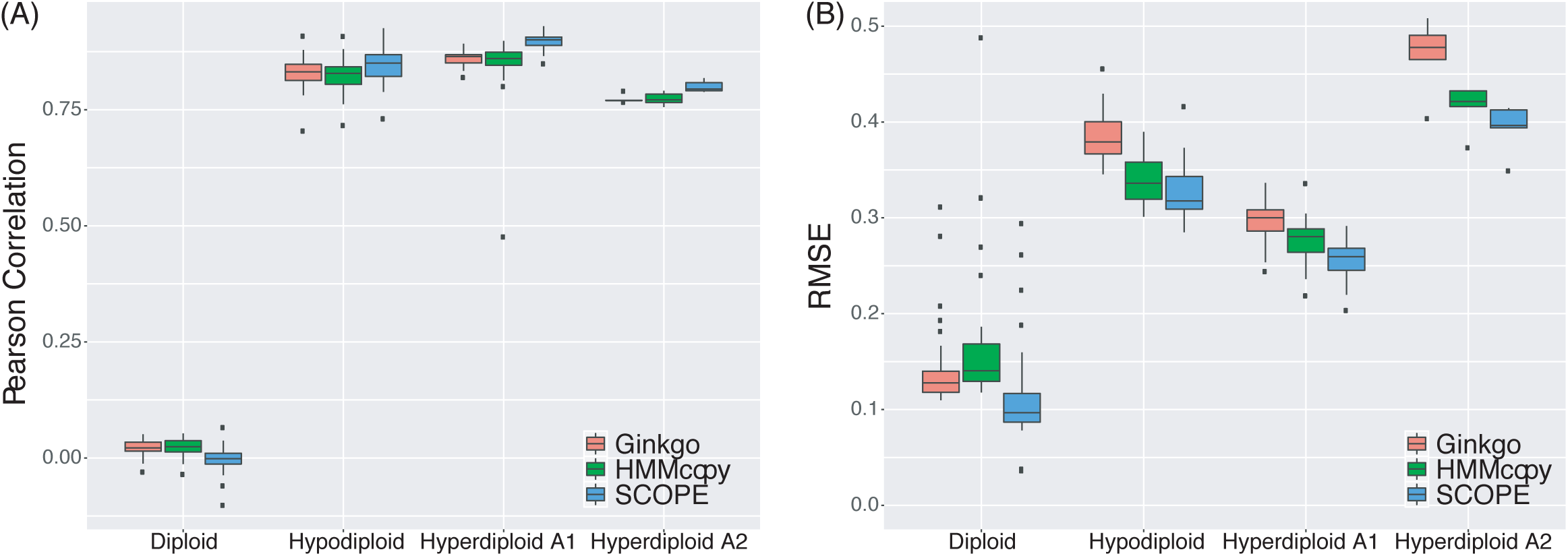
Orthogonal validation of single-cell copy number profiles by aCGH of purified bulk samples by FACS. Spearman correlation and RMSE as performance metrics comparing results by Ginkgo, and HMMcopy, and SCOPE against the gold standard. SCOPE achieves higher correlation and lower estimation errors. Extreme outliers outside the vertical plotted range for HMMcopy were omitted.

### Analysis of scDNA-seq data of triple-negative breast cancer patients with paired WES and scRNA-seq

We further applied SCOPE to scDNA-seq data of temporally separated tumor resections from triple negative breast cancer patients with chemotherapy treatment.^24^ Specifically, scDNA-seq was performed on matched pre-treatment, mid-treatment, and post-treatment tumor samples from the same patient. We applied SCOPE to three “clonal extinction” patients, where tumor cells were previously reported to exist only in the pre-treatment samples (Supplementary Table S1). For example, 92 cells from patient KTN302 were sequenced at two timepoints, pre- and mid-treatment. In this case, SCOPE successfully detected two subclones of aneuploid cells in the pre-treatment tumors and found that all the cells from the mid-treatment group were normal, as shown in Figure 6A. The consensus copy number profiles of the two subpopulations indicate a large number of shared CNAs between the two inferred subclones; this is likely due to a punctuated copy number evolution,^47^ which produces the majority of CNAs in the early stages of tumor evolution. Results for patients KTN126 and KTN129 are included in Supplementary Figure S10.

**Figure 6.**
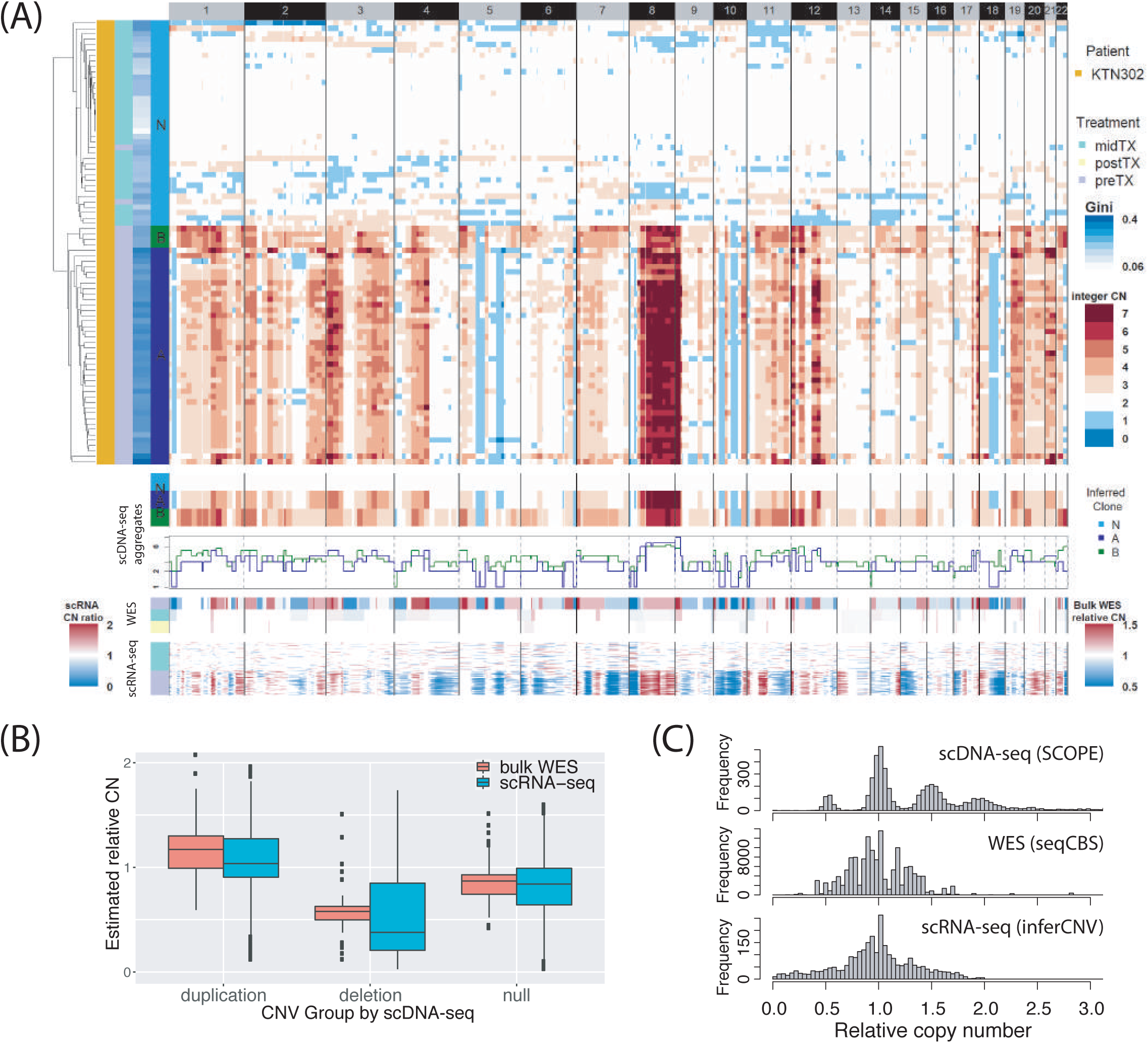
Copy-number profiles of triple-negative breast cancer patients KTN302, inferred by scDNA-seq, bulk WES, and scRNA-seq. (A) Upper three panels show heatmap of inferred copy-number profiles by SCOPE with cells clustered by hierarchical clustering, heatmap and stairstep plot of consensus copy number profiles for each cell subpopulation (N for normal cell, A, B, and C for cancer subclones). Lower two panels show copy number profiles inferred by WES and scRNA-seq, which are further used to validate CNV calls by SCOPE. (B) Orthogonal validations using copy number profiles by WES and scRNA-seq. Each genomic segment is categorized as amplification (amp), deletion (del), or copy-number neural (null) based on the copy number estimates returned by SCOPE. The boxplot shows the estimated relative copy numbers by bulk WES and scRNA-seq, which are higher in amplified regions and lower in deleted regions, compared to the null regions. For scRNA-seq and bulk WES, the relative copy numbers are estimated comparing to a sample-specific baseline – for cases where the tumor cells are hyperdiploid/hypodiploid, the baseline is shifted up/down making the true null regions have relative copy numbers less/greater than one. (C) Relative copy numbers estimated by scDNA-seq, WES, and scRNA-seq. For WES and scRNA-seq, the copy number signals do not show clear separation between different states and further have mean one by taking the sample mean as reference. This over/under normalizes ploidy in hyper/hypodiploid cells resulting in both false negatives and false positives.

We used bulk-tissue WES and single-cell RNA-seq data to validate the copy number profiles returned by SCOPE. For the data from patient KTN302, WES and scRNA-seq were performed on the matched normal (blood), pre-treatment, mid-treatment, and post-treatment bulk samples (Supplementary Table S1) and were used to profile CNAs separately for validation. For bulk WES, a modified CBS algorithm based on seqCBS^41^ was utilized for the paired tumor-normal experimental design, which returns relative copy numbers (i.e., the ratios of cancer cell copy number to normal cell copy number). Despite signal attenuation due to intra-tumor heterogeneity and normal cell contamination, we observe large deletions on chromosome 5, 10, and 18, and a distinct copy-number amplification on chromosome 8, in accordance with the profiles returned by scDNA-seq (Figure 6A, Supplementary Figure S10). For scRNA-seq, we used the InferCNV toolkit^48^ and employed a sliding-window approach of 50 genes^49^ to infer CNVs, using as input the transcript per million (TPM) matrix returned by SALMON (Supplementary Figure S11).^50^ In spite of the low resolution, we observed deletions on chromosome 5, 10, and 18, as well as duplications on chromosome 8, in accordance with WES and scDNA-seq calls (Figure 6A, Supplementary Figure S10).

To quantitatively assay the quality of the call set produced by SCOPE, we plotted the relative copy number estimates by WES and scRNA-seq for the inferred amplified, deleted, and copy-number-neutral regions by SCOPE. Although the copy number signals from WES and scRNA-seq are of low resolution and are attenuated due to normal cell contaminations, the relative copy numbers (i.e., absolute copy numbers divided by ploidy) from both WES and scRNA-seq are less than one for deletion regions, but greater than one for putative amplifications, a finding which agrees with results produced by SCOPE (Figure 6B, Supplementary Figure S12). While we demonstrated that the deletion and duplication signals by SCOPE using scDNA-seq can be recapitulated from two orthogonal platforms, we also found that, in the WES and scRNA-seq data, the relative copy numbers in the null regions are less than one and that the signals for duplications are dampened towards the null (Figure 6B, Supplementary Figure S12). This unexpected result indicates a potential pitfall associated with profiling CNVs by bulk DNA-seq and scRNA-seq: all copy number events are defined in reference to the population average, resulting in a genome-wide mean of one for the relative copy number estimates (Figure 6C). This masks the ploidy and further results in over-normalization of ploidy in hyperdiploid samples and under-normalization of ploidy in hypodiploid samples. SCOPE solves this problem by directly estimating discretized copy numbers of integer values, which, unlike the traditionally continuous copy numbers, are free of whole-genome bias due to ploidy (Figure 6C).

### Analysis of scDNA-seq data of gastric cancer spike-ins and breast cancer dissections from the 10X Genomics

We finally demonstrated SCOPE on three recently released scDNA-seq datasets from the 10X Genomics Single-Cell CNV Solution pipeline. We first adopted two publicly available spike-in datasets from the 10X Genomics website, where 1% and 10% MKN-45 gastric cancer cell lines are mixed with ∼500 and ∼1000 normal BJ fibroblast cells, respectively (Supplementary Table S1). The second dataset consists of ∼10,000 nuclei extracted and sequenced from five different dissections of frozen breast tumor tissue from a triple negative ductal carcinoma (Supplementary Table S1).

As a proof of concept, we started by applying SCOPE to the two 10X Genomics spike-in datasets, sequencing 1,055 single cells with a 1% spike-in of cancer cells and 462 single cells with a 10% spike-in of cancer cells. We adopted stringent quality control procedures to remove cells with extreme Gini coefficients and to remove bins that (i) have low mappability; (ii) reside in “blacklist” regions; and (iii) have extremely low read depth across cells. For the two spike-in datasets, this resulted in 5,055 and 5,064 bins for the copy number estimation, respectively. Despite the higher sparsity and lower sequencing depth of this data, SCOPE successfully identified the cancer cell cluster from the cluster of normal diploid cells (heatmaps shown in Supplementary Figure S13). Specifically, 11 and 36 cancer cells were identified from the 1% and 10% spike-in datasets respectively, with estimated proportions as 1% and 8%. For data visualization, we performed t-SNE projections^51^ on the normalized *z*-scores, revealing that SCOPE successfully recovers 1% and 10% non-diploid cancer cells from a mixture of diploid cells in the background (Figure 7A-B). The colors in Figure 7A-B indicate the classification by a threshold of 0.12 for the Gini coefficients. We see that while not all diploid cells are used as negative control cells, there is a perfect separation between normal cells and cancer cells (see “Performance assessment via spike-in studies and with varying parameters” for details on setting the threshold of Gini coefficients).

**Figure 7.**
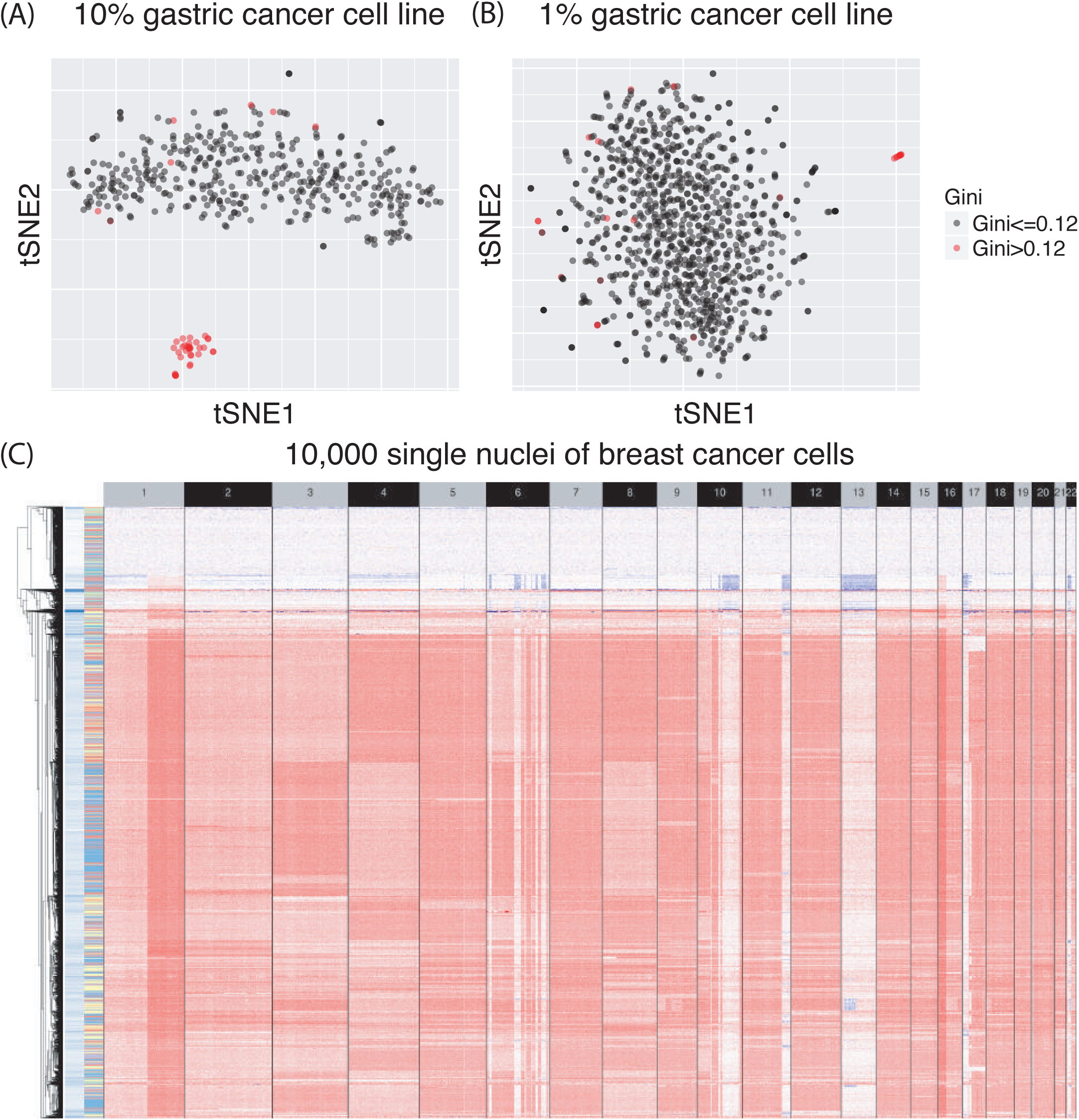
Visualization by t-SNE projections of normalized scDNA-seq data from the 10X Genomics cancer spike-in datasets. (A) 1% and (B) 10% of gastric cancer cell lines were mixed with normal fibroblast cell lines. After normalization, SCOPE successfully identified the cancer cell population from the background of normal cells. (C) Heatmap of normalization results by SCOPE on the scDNA-seq dataset of 10,000 breast cancer nuclei. Cell clustering reveals cancer subclonal structure, not previously detectable by bulk DNA-seq.

We further applied SCOPE to a 10X Genomics scDNA-seq dataset of ∼10,000 cells from five adjacent tumor dissections of a breast cancer patient. We followed the same quality control procedure as previously described, which resulted in 4864, 5016, 5032, 5023, and 5026 bins from each dissection. SCOPE was then separately applied to each dissection for identification of normal cells, read depth normalization, and copy number estimation. The proportions of normal cells vary across the five dissections, with mean cellular ploidy of sections A to E being 2.12, 3.13, 3.29, 3.36, and 3.53, respectively. This indicates a gradient of normal cells contaminating the tumor (Supplementary Figure S14). We further integrated the inferred copy number profiles by SCOPE across all tumor cells from the five dissections and demonstrated that SCOPE was able to identify subclonal structures. For example, the distinct duplication events on chromosome 3 and chromosome 4 are mutually exclusive and mark a split in the tumor evolutionary history. Hierarchical clustering based on the normalization result suggests that the different cancer subclones consist of cancer cells from all sections. This indicates an early branching evolutionary model, where copy number aberrations happen at a potentially early stage in the disease advancement (Figure 7C).

### Performance assessment via spike-in studies and with varying parameters

To further assess performance of SCOPE and to benchmark against existing methods, we conducted *in silico* spike-in studies. We started with the read depth data of diploid cells from breast cancer patient T10^22^ and applied stringent filtering steps to remove any regions harboring potential sporadic CNVs, resulting in 39 normal cells and 1,545 genomic regions that are CNV-free. We then added multiple CNV signals to the background count matrix under the null. These signals had different copy number states but shared changepoints across cells. generate these CNV signals, we scaled the raw depth of coverage spanned by the CNV from *y* to *y* × *c*/2, where *c* is sampled from a normal distribution, with mean equal to the underlying copy number and standard deviation 0.1. We ran simulations under three designs: (i) spike-ins with copy number states 1, 2, and 3 (Supplementary Figure S15A); (ii) spike-ins with copy number states 2, 3, and 4 (Supplementary Figure S16A); and (iii) spike-ins with copy number states 1, 2, and 4 (Supplementary Figure S17A). The different copy number states had varying genome-wide incidence rates (Supplementary Table S4, S5, S6) and each simulation was repeated 20 times. Overall, when compared to Ginkgo and HMMcopy, SCOPE returned the highest precision and recall rates under all simulation settings (Supplementary Table S4, S5, S6). Further inspection of the normalized-scores by the three methods showed that the normalization results by SCOPE have lower variability for deletion, null, and duplication regions across all runs compared to the other two methods (Supplementary Figure S15B, S16B, S17B). In several cases, HMMcopy tended to return copy number estimates that were inflated across the entire genome (Supplementary Figure S15B). This led to false negatives for true deletions, as well as false positives for true null regions. And in concordance with a recent benchmark report,^36^ HMMcopy could not correctly predict the absolute copy numbers in the absence of intermediate copy numbers (Supplementary Table S6, Supplementary Figure S17).

To systematically investigate how performance is influenced by varying parameters (e.g., bin size, threshold of cell-specific Gini coefficients to identify normal controls, and the number of Poisson latent factors), we carry out additional evaluation and benchmark analysis on the scDNA-seq data from the breast cancer patient T10, with the aCGH calls as gold standard. Our results indicate that a bin size of 3Mb, compared to a bin size of 200Kb, leads to smoother normalization results, but it misses smaller CNVs due to low resolution (Supplementary Figure S18). Bin sizes between 200Kb and 1Mb do not affect the performance of SCOPE, especially for chromosome-arm level CNVs (Supplementary Figure S19). To make the optimal choice of the Gini coefficient threshold, SCOPE does not need to include all normal cells as negative controls – 10 to 20 normal cells suffice to achieve accurate estimation (Supplementary Figure S20). We provide empirical evidence for this claim in the section “Performance assessment via spike-in studies and with varying parameters”. In addition, we show that SCOPE’s, the number of Poisson latent factors (Supplementary Figure S21), since SCOPE adopts the EM algorithm to account for genomic contexts when estimating GC content biases and uses only the normal cells to estimate the bin-specific noise terms. As such, we show that SCOPE is robust to the choice of Gini coefficient threshold and the number of Poisson latent factors. In summary, compared to existing methods, SCOPE directly provides copy number and ploidy estimates, and it does so completely off the shelf without much need for manual tuning by the users.

## Discussion

Here we propose SCOPE, a new method to remove technical noise and improve CNV signal-to-noise ratio for scDNA-seq data. SCOPE includes a Poisson latent factor model for data normalization and a non-parametric functional term for adjusting GC content bias in a context-specific manner. The EM algorithm embedded in the Poisson log linear model accounts for the different copy number states as it adjusts for GC content bias—a step we show to be crucial for cancer genomic studies, where recurrent copy number changes across the genome lead to significant deviations from the null. Capacity is increasing in single-cell isolation and single-cell whole-genome sequencing, and there is an increasing need to profile CNVs, which are a non-negligible source of genetic variation, at the single-cell level. We believe that SCOPE can be a useful tool for the genetics and genomics community, for it lays the statistical foundation that will make robust and accurate single-cell CNV profiling possible.

We demonstrated SCOPE on two breast cancer scDNA-seq datasets, where SCOPE successfully parsed normal cells, exposed a mixture of tumor cells based on their inferred copy number profiles, and unmasked the ploidy. We showed that this enabled characterization of genetically distinct populations of tumor cells that were not evident in bulk-tissue analysis, shedding light upon tumor heterogeneity at a much finer resolution. We benchmarked SCOPE against other existing methods, using array-based CNV calls from purified bulk samples as the gold standard, and showed that SCOPE outperformed those methods. We further demonstrated SCOPE on scDNA-seq data from the recently released Single Cell CNV Solution by 10X Genomics and showed that SCOPE could accurately detect 1% of the gastric cancer cells from the background of normal fibroblast cells, and that it successfully reconstructed the subclonal structure across 10,000 breast cancer cells.

While copy number profiling is important beyond cancer, scDNA-seq holds great promise for deciphering tumor heterogeneity. Due to the low depth of coverage, scDNA-seq has thus far been primarily used to profile somatic CNVs in cancer cells, and the resolutions of the detected breakpoints range from hundreds of kilobases to megabases. Therefore, SCOPE by default is designed for large CNV detection in cancer cells; for relatively short CNVs and for common germline CNVs, the sensitivity is low due to technological limitations. The running time across all adopted datasets for both normalization and segmentation are shown in Supplementary Table S7. It is recommended that normalization with different numbers of Poisson latent factors and segmentation across different chromosomes be run in parallel on a high-performance cluster.

Correct inference of tumor evolution history depends on accurate copy number estimates, which in turn depend on effective normalization. While we show that data normalization is a first-order concern, SCOPE’s cross-sample segmentation procedure also returns changepoints that are shared across cells from the same genetic backgrounds, a unique feature that is necessary for the single-cell setting. SCOPE also enables hierarchical clustering and phylogeny reconstruction, based on the normalized log-ratios, the estimated copy numbers, or the estimated changepoints—data that is useful for further downstream analysis. To further reduce the ambiguity in unmasking intra-tumor heterogeneity, future research may include force-calling somatic point mutations from scDNA-seq and integrating both copy number and point mutation profiles for reconstructing tumor evolutionary history; a recent report demonstrated that merging cell subsets with shared copy numbers enabled inference of clone-specific single-nucleotide resolution events and clonal phylogenies.^34^

A few recently developed methods for CNV detection use scRNA-seq.^48,52,53^ While these gene-expression based approaches have been successfully applied to detect chromosome or chromosome-arm level CNAs, multimodal data alignment between scDNA-seq and scRNA-seq has the potential to increase the resolution in detecting CNAs in a larger cell population. This alignment could also shed light upon the interplay between genomic and transcriptomic variations in cancer, for it is still unclear how transcriptomic variation is modulated by genetic variation and phylogenetic evolution in cancer. The joint analysis framework would allow quantification of clonal differences in gene expression while accounting for DNA confounding.

## Acknowledgements

This work was supported by the National Institutes of Health (NIH) grant P01 CA142538 (to DYL and YJ), R35 GM118102 (to YJ), a developmental award from the UNC Lineberger Comprehensive Cancer Center 2017T109 (to YJ), and a pilot award from the UNC Computational Medicine Program (to YJ).

## Authors’ Contributions

YJ initiated and envisioned the study. RW and YJ formulated the model, developed and implemented the algorithm. RW, DYL, and YJ performed data analysis. RW and YJ wrote the manuscript, which was edited by DYL. All authors read and approved the final manuscript.

## Supplementary Notes

### Note S1: Bioinformatic pre-processing and binning method

In this paper, we adopted publicly available scDNA-seq data from two different sources: data from the NCBI Sequence Read Archive (SRA) and data from the 10X Genomics website. For data from the NCBI, we adopt the following bioinformatic pre-processing procedures: (i) obtain fastq files from SRA files using fastq-dump from the NCBI SRA toolkit; (ii) align reads using BWA;^1^ (iii) add read group, dedup, sort, and index bam files using SAMtools.^2^ For data from the 10X Genomics, reads that contain cellular barcodes from the barcode list of interest are demultiplexed using a Python script. Demultiplexed reads with error-corrected and confirmed cellular barcodes are used for further analysis. The second and third steps follow the same as above.

To computer read depth, SCOPE enables reconstruction of user-defined genome-wide consecutive bins prior to downstream analysis. Existing analyses adopt a variable binning strategy, where in silico generated reads of the reference assembly are mapped back to the genome and bins are selected such that each bin has the same number of uniquely mappable reads.^3,4^ However, this method still does not directly offer an optimal length of the bins while and the bin size directly affects the resolution at which CNVs can be called. As such, Ginkgo enables binning with either variable or fixed size.^5^

Here, SCOPE by its default adopts a fixed binning method to compute the depth of coverage while removing reads that are mapped to multiple genomic locations and to “blacklist” regions. This is followed by an additional step of quality control to remove bins with extreme mappability to avoid erroneous detections. Specifically, “blacklist” bins, including segmental duplication regions (http://humanparalogy.gs.washington.edu/build37/data/GRCh37GenomicSuperDup.tab) and gaps in reference assembly from telomere, centromere, and/or heterochromatin regions (https://gist.github.com/leipzig/6123703), are masked by default prior to getting coverage. Furthermore, to compute mappability for hg19, we employed the 100-mers mappability track from the ENCODE Project6 and computed weighted average of the mappability scores if multiple ENCODE regions overlap with the same bin. SCOPE further filters out bins with low mappability with a default threshold of 0.9 to reduce artifacts.

To calculate mappability for hg38, we adopted the UCSC liftOver utility (http://hgdownload.cse.ucsc.edu/goldenpath/hg19/liftOver/). For other reference genomes, we first construct consecutive reads that are one base pair apart along the bin. The length of the reads is set to be the same as that from the sequencing platform and the read sequences are taken from the reference genome of interest. We then find possible positions across the genome that the reads can map to allowing for a default number of mismatches. Finally, we compute the mean of the probabilities that the overlapped reads map to the target places where they are generated and use this as the mappability of the bin.

### Note S2: Iterative parameter estimation procedure for SCOPE

#### Initialization

Identify negative control cells using cell-specific Gini coefficients. Apply Poisson latent factor model with negative controls to obtain 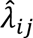 and initialize 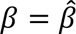. Let 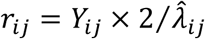 be the estimated relative copy number, which has mean 2 across all cells. Denote *P_j_* as the ploidy for cell *j* and 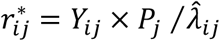 as the absolute copy number. We pre-estimate *P_j_* for EM initialization:

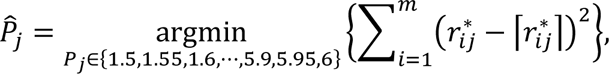

where 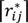 rounds 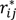 to the nearest integer and the square error between 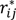 and 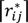 is minimized.

#### Iteration

1. Within each iteration (given *β*, *g*, and *h* from previous iteration)
  a. a)M-step: 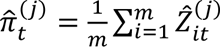 for all *t* ∈ {1, ⋯, *T_j_*} For each cell *j*, fit the LOESS curve of 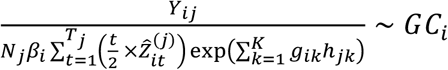 and use the fitted value as *f_j_*(*GC_i_*).
  b. b)E-step:

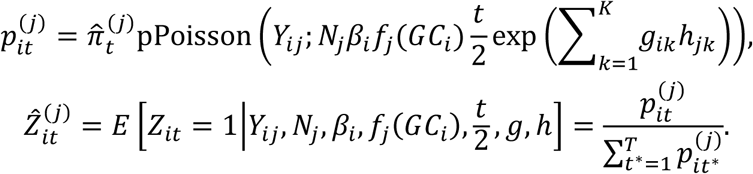
  c. Repeat a) – b) till convergence.
2. Within each iteration (given *g*, *h*, *f*(*GC*), *Z*^(1)^, ⋯, *Z*^(n)^) Let *J_c_* be the indices of negative control samples/cells.

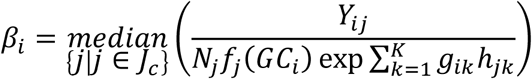
3. Within each iteration (given *β*, *f*(*GC*), *Z*^(1)^, ⋯, *Z*^(n)^) Let *h^old^* be the estimated *h* from the previous step.
  a. For each bin *i*, fit Poisson log-linear regression with *Y_iJc_* as response, 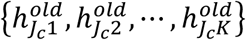 as covariates, 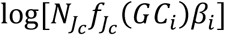 as fixed offset to obtain updated estimates as {*g*_*i*1_, ⋯, *g_ik_*}.
  b. For each cell *j*, fit Poisson log-linear regression with *Y_:j_* as response, {*g*_*i*1_, ⋯, *g_ik_*} as covariates, 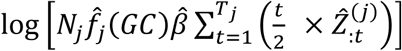 as fixed offset to obtain updated estimates 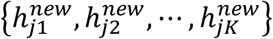
  c. Center each row of *g* × (*h*^*new*^)^*T*^ and apply SVD to the row-centered matrix to obtain the *K* right singular vectors to update *h^new^*.
  d. Repeat a) – c) with *h^old^* = *h^new^* till convergence.
4. Repeat steps 1 – 3 till convergence.

## Supplementary Tables

**Table S1:** Summary on bulk and single-cell datasets adopted in this study. (A) scDNA-seq data from two breast cancer patients T10 and T16.^3^ (B) scDNA-seq data of four breast cancer patients.^7^ CNV profiles by bulk whole-exome sequencing (WES) and scRNA-seq data were used for benchmarking. (C) scDNA-seq data from the 10X Genomics platform (https://www.10xgenomics.com/solutions/single-cell-cnv).

(A) Navin et al., Nature, 2011
| Protocol | Sample | # of cells | Total read counts per cell | Remarks |
| --- | --- | --- | --- | --- |
| DOP-PCR | T10 | 100 | 4,287,124 | Cell SRR054599 and SRR053670 from T10 were removed due to low proportion of mapped reads; cell SRR089717 from T16M was excluded due to extreme Gini index as an outlier. |
|  | T16P | 52 | 8,847,387 |  |
|  | T16M | 48 | 11,651,933 |  |

(B) Kim et al., Cell, 2018
| Protocol | Tumor type | Patient ID | # of cells by scDNA-seq | Median total read counts per cell | # of cells by scRNA-seq | # of bulk samples by WES | Bulk sample time points |
| --- | --- | --- | --- | --- | --- | --- | --- |
| DOP-PCR | Extinction | P1/KTN126 | 93 | 3,869,140 | 735 | 3 | blood+pre+post |
|  | Extinction | P2/KTN129 | 90 | 2,856,810 | 885 | 4 | blood+pre+mid+post |
|  | Extinction | P6/KTN206 | N.A. | N.A. | 881 | 3 | blood+pre+post |
|  | Extinction | P9/KTN302 | 92 | 2,998,194 | 880 | 4 | blood+pre+mid+post |

(C) 10X Genomics
| Type | # of cells | Median total read counts per cell | # of cells after stringent QC |
| --- | --- | --- | --- |
| BJ with 1% MKN45 Spike-In | 1056 | 528,751 | 1055 |
| BJ with 10% MKN45 Spike-In | 462 | 731,877 | 462 |
| Breast Tissue nuclei section A | 2173 | 291,457 | 2164 |
| Breast Tissue nuclei section B | 2224 | 347,381 | 2185 |
| Breast Tissue nuclei section C | 1722 | 422,030 | 1705 |
| Breast Tissue nuclei section D | 1916 | 337,901 | 1898 |
| Breast Tissue nuclei section E | 2053 | 414,925 | 2032 |

**Table S2:** Three conventional whole-genome amplification methods for scDNA-seq. Degenerate oligonucleotide primed PCR (DOP-PCR); multiple displacement amplification (MDA); multiple annealing and looping-based amplification cycles (MALBAC) have been proposed for whole-genome DNA amplification at the single-cell level, each with pros and cons. Table was adapted from Liu et al..^8^

| Protocols | Description | Pros | Cons |
| --- | --- | --- | --- |
| DOP-PCR<sup>9</sup> | PCR-based amplification using degenerate oligonucleotides and thermostable polymerase | High uniformity (better for calling CNVs) | High error rate; low coverage |
| MDA<sup>10</sup> | Isothermal amplification of randomly primed regions of genome using random hexamers and Phi29 polymerase | Low error rate; great coverage (better for calling SNVs) | Lack of uniformity |
| MALBAC<sup>11</sup> | Limited isothermal amplification using degenerate primers followed by PCR amplification | High uniformity (better for calling CNVs) | Intermediate error rate |

**Table S3: Gold-standard CNV calls from array comparative genomic hybridization (aCGH).** The aCGH experiments were performed on six tumor sectors (S1-S6) from T10 after FACS sorting. Distinct ploidy distributions were reported --hypodiploid, aneuploid 1, aneuploid 2, and diploid,^12^ each with array intensities corresponding to relative copy numbers. We utilized the UCSC liftOver utility (http://hgdownload.cse.ucsc.edu/goldenpath/hg17/liftOver/) to convert the hg17 coordinates to hg19 and adopted stringent quality control procedures to remove bins with low mappability and bin within “blacklist” regions. Aneuploid 1 sector from S4, and diploid sectors from S5 and S6 were excluded due to bad quality. Table is separately attached.

**Table S4.** Performance assessment via spike-in analysis with copy number states 1, 2, and 3. Copy number signals were manually added to the depth of coverage data from the null genomic regions of patient T10. SCOPE is benchmarked against Ginkgo and HMMcopy and show its outperformance with higher precision and recall rates.

| frequency |  |  | Ginkgo |  | HMMcopy |  | SCOPE |  |
| --- | --- | --- | --- | --- | --- | --- | --- | --- |
| cn=1 | cn=2 | cn=3 | precision | recall | precision | recall | precision | recall |
| 0.10 | 0.30 | 0.60 | 0.99 (0.01) | 0.91 (0.01) | 0.31 (0.05) | 0.16 (0.04) | 1.00 (0.00) | 0.97 (0.01) |
| 0.20 | 0.30 | 0.50 | 0.98 (0.01) | 0.94 (0.01) | 0.47 (0.12) | 0.31 (0.20) | 1.00 (0.00) | 0.98 (0.01) |
| 0.30 | 0.30 | 0.40 | 0.94 (0.01) | 0.97 (0.01) | 0.56 (0.01) | 0.58 (0.01) | 1.00 (0.00) | 0.98 (0.01) |
| 0.40 | 0.30 | 0.30 | 0.93 (0.01) | 0.97 (0.01) | 0.50 (0.02) | 0.45 (0.01) | 1.00 (0.00) | 0.99 (0.01) |
| 0.50 | 0.30 | 0.20 | 0.93 (0.01) | 0.97 (0.01) | 0.40 (0.03) | 0.31 (0.01) | 1.00 (0.00) | 0.99 (0.00) |
| 0.60 | 0.30 | 0.10 | 0.94 (0.01) | 0.97 (0.01) | 0.27 (0.03) | 0.18 (0.01) | 1.00 (0.00) | 1.00 (0.00) |

**Table S5.**
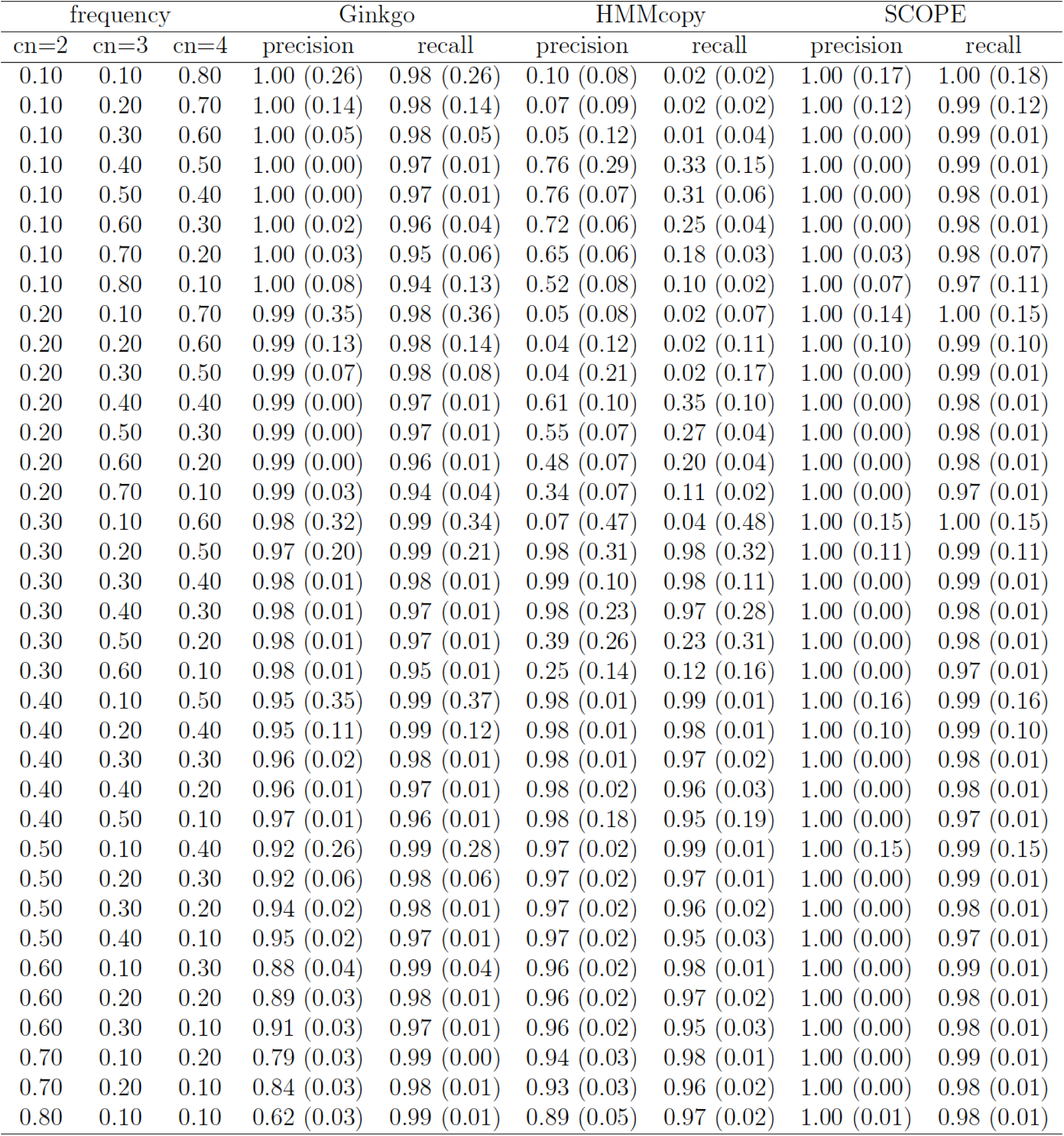
Performance assessment via spike-in analysis with copy number states 2, 3, and 4. Copy number signals were manually added to the depth of coverage data from the null genomic regions of patient T10. SCOPE is benchmarked against Ginkgo and HMMcopy and show its outperformance with higher precision and recall rates.

**Table S6.** Performance assessment via spike-in analysis with copy number states 1, 2, and 4. Copy number signals were manually added to the depth of coverage data from the null genomic regions of patient T10. SCOPE is benchmarked against Ginkgo and HMMcopy and show its outperformance with higher precision and recall rates.

| frequency |  |  | Ginkgo |  | HMMcopy |  | SCOPE |  |
| --- | --- | --- | --- | --- | --- | --- | --- | --- |
| cn=1 | cn=2 | cn=4 | precision | recall | precision | recall | precision | recall |
| 0.10 | 0.80 | 0.10 | 0.85 (0.04) | 1.00 (0.01) | 0.90 (0.05) | 1.00 (0.00) | 1.00 (0.01) | 1.00 (0.00) |
| 0.10 | 0.10 | 0.80 | 0.99 (0.08) | 1.00 (0.12) | 0.54 (0.04) | 0.13 (0.02) | 1.00 (0.03) | 1.00 (0.09) |
| 0.10 | 0.30 | 0.60 | 0.97 (0.11) | 1.00 (0.13) | 0.28 (0.24) | 0.17 (0.28) | 1.00 (0.08) | 1.00 (0.10) |
| 0.10 | 0.40 | 0.50 | 0.93 (0.12) | 1.00 (0.14) | 0.98 (0.01) | 1.00 (0.00) | 1.00 (0.08) | 1.00 (0.08) |
| 0.10 | 0.50 | 0.40 | 0.89 (0.07) | 0.99 (0.06) | 0.97 (0.01) | 1.00 (0.00) | 1.00 (0.00) | 1.00 (0.00) |
| 0.10 | 0.60 | 0.30 | 0.86 (0.02) | 0.99 (0.00) | 0.96 (0.02) | 1.00 (0.00) | 1.00 (0.00) | 1.00 (0.00) |
| 0.10 | 0.70 | 0.20 | 0.70 (0.02) | 0.99 (0.01) | 0.94 (0.03) | 1.00 (0.00) | 1.00 (0.00) | 1.00 (0.00) |
| 0.10 | 0.20 | 0.70 | 0.99 (0.14) | 1.00 (0.18) | 0.37 (0.03) | 0.15 (0.02) | 1.00 (0.09) | 1.00 (0.13) |
| 0.20 | 0.10 | 0.70 | 1.00 (0.00) | 1.00 (0.00) | 0.87 (0.08) | 0.78 (0.24) | 1.00 (0.00) | 1.00 (0.00) |
| 0.20 | 0.30 | 0.50 | 0.93 (0.01) | 0.99 (0.01) | 0.63 (0.12) | 0.72 (0.11) | 1.00 (0.00) | 1.00 (0.00) |
| 0.20 | 0.40 | 0.40 | 0.88 (0.01) | 0.98 (0.01) | 0.97 (0.23) | 0.99 (0.16) | 1.00 (0.00) | 1.00 (0.00) |
| 0.20 | 0.50 | 0.30 | 0.84 (0.02) | 0.98 (0.01) | 0.97 (0.15) | 1.00 (0.10) | 1.00 (0.00) | 1.00 (0.00) |
| 0.20 | 0.60 | 0.20 | 0.84 (0.03) | 0.98 (0.01) | 0.96 (0.12) | 1.00 (0.08) | 1.00 (0.00) | 1.00 (0.00) |
| 0.20 | 0.70 | 0.10 | 0.94 (0.03) | 1.00 (0.00) | 0.94 (0.06) | 1.00 (0.04) | 1.00 (0.00) | 1.00 (0.00) |
| 0.20 | 0.20 | 0.60 | 0.97 (0.02) | 0.99 (0.02) | 0.75 (0.10) | 0.76 (0.20) | 1.00 (0.00) | 1.00 (0.00) |
| 0.30 | 0.30 | 0.40 | 0.89 (0.02) | 0.97 (0.02) | 0.56 (0.01) | 0.58 (0.01) | 1.00 (0.00) | 1.00 (0.00) |
| 0.30 | 0.40 | 0.30 | 0.84 (0.02) | 0.97 (0.01) | 0.43 (0.19) | 0.51 (0.16) | 1.00 (0.00) | 1.00 (0.00) |
| 0.30 | 0.50 | 0.20 | 0.82 (0.02) | 0.97 (0.01) | 0.97 (0.31) | 1.00 (0.26) | 1.00 (0.00) | 1.00 (0.00) |
| 0.30 | 0.10 | 0.60 | 0.97 (0.02) | 0.97 (0.02) | 0.83 (0.02) | 0.67 (0.01) | 1.00 (0.00) | 1.00 (0.00) |
| 0.30 | 0.20 | 0.50 | 0.93 (0.01) | 0.97 (0.01) | 0.70 (0.02) | 0.63 (0.01) | 1.00 (0.00) | 1.00 (0.00) |
| 0.30 | 0.60 | 0.10 | 0.83 (0.03) | 0.98 (0.01) | 0.96 (0.20) | 1.00 (0.18) | 1.00 (0.00) | 1.00 (0.00) |
| 0.40 | 0.30 | 0.30 | 0.87 (0.01) | 0.95 (0.01) | 0.50 (0.02) | 0.45 (0.01) | 1.00 (0.00) | 1.00 (0.00) |
| 0.40 | 0.40 | 0.20 | 0.83 (0.02) | 0.95 (0.01) | 0.34 (0.02) | 0.36 (0.01) | 1.00 (0.00) | 1.00 (0.00) |
| 0.40 | 0.10 | 0.50 | 0.96 (0.02) | 0.94 (0.02) | 0.81 (0.03) | 0.57 (0.01) | 1.00 (0.00) | 1.00 (0.00) |
| 0.40 | 0.20 | 0.40 | 0.91 (0.01) | 0.94 (0.01) | 0.66 (0.03) | 0.52 (0.01) | 1.00 (0.00) | 1.00 (0.00) |
| 0.40 | 0.50 | 0.10 | 0.84 (0.02) | 0.97 (0.01) | 0.19 (0.20) | 0.23 (0.19) | 1.00 (0.00) | 1.00 (0.00) |
| 0.50 | 0.30 | 0.20 | 0.85 (0.01) | 0.92 (0.02) | 0.40 (0.03) | 0.31 (0.01) | 1.00 (0.00) | 1.00 (0.00) |
| 0.50 | 0.10 | 0.40 | 0.95 (0.01) | 0.91 (0.02) | 0.77 (0.04) | 0.46 (0.01) | 1.00 (0.00) | 1.00 (0.00) |
| 0.50 | 0.20 | 0.30 | 0.90 (0.01) | 0.91 (0.01) | 0.59 (0.03) | 0.40 (0.01) | 1.00 (0.00) | 1.00 (0.00) |
| 0.50 | 0.40 | 0.10 | 0.85 (0.02) | 0.95 (0.01) | 0.22 (0.02) | 0.20 (0.02) | 1.00 (0.00) | 1.00 (0.00) |
| 0.60 | 0.10 | 0.30 | 0.94 (0.03) | 0.86 (0.03) | 0.71 (0.04) | 0.36 (0.01) | 1.00 (0.00) | 1.00 (0.00) |
| 0.60 | 0.30 | 0.10 | 0.87 (0.01) | 0.93 (0.02) | 0.27 (0.03) | 0.17 (0.02) | 1.00 (0.00) | 1.00 (0.00) |
| 0.60 | 0.20 | 0.20 | 0.88 (0.01) | 0.88 (0.02) | 0.49 (0.04) | 0.28 (0.01) | 1.00 (0.00) | 1.00 (0.00) |
| 0.70 | 0.10 | 0.20 | 0.91 (0.01) | 0.74 (0.03) | 0.63 (0.08) | 0.25 (0.11) | 1.00 (0.00) | 1.00 (0.00) |
| 0.70 | 0.20 | 0.10 | 0.86 (0.01) | 0.85 (0.02) | 0.34 (0.05) | 0.16 (0.04) | 1.00 (0.00) | 1.00 (0.00) |
| 0.80 | 0.10 | 0.10 | 0.13 (0.31) | 0.12 (0.20) | 0.48 (0.07) | 0.14 (0.03) | 0.54 (0.21) | 0.14 (0.38) |

**Table S7:** Processing time of SCOPE for normalization and cross-sample segmentation. Normalization was ran with one Poisson latent factor and varying *T_j_* ∈ {1, 2, 3, 4, 5} for each cell. Parallel computing on a high-performance cluster (HPC) is recommended for varying number of Poisson latent factors. Cross-sample segmentation processes the entire genome chromosome by chromosome in parallel; the running time is evaluated by taking the mean. Time unit is hour (h). Jobs were run with 80GB RAM on an HPC.

| Dataset | % of negative control cells | Normalization | Segmentation |
| --- | --- | --- | --- |
| T10 | 0.40 (39/98) | 0.56 | 0.06 |
| T16 | 0.49 (49/99) | 0.24 | 0.06 |
| BJ with 1% MKN45 Spike-In | 0.98 (1029/1055) | 0.15 | 0.64 |
| BJ with 10% MKN45 Spike-In | 0.91 (421/462) | 0.20 | 0.27 |
| Breast Tissue nuclei section A | 0.85 (1829/2164) | 1.57 | 1.39 |
| Breast Tissue nuclei section B | 0.33 (718/2185) | 3.54 | 1.36 |
| Breast Tissue nuclei section C | 0.21 (363/1705) | 4.46 | 1.01 |
| Breast Tissue nuclei section D | 0.21 (401/1898) | 4.56 | 1.17 |
| Breast Tissue nuclei section E | 0.16 (323/2032) | 4.67 | 1.39 |

## Supplementary Figures

**Figure S1.**
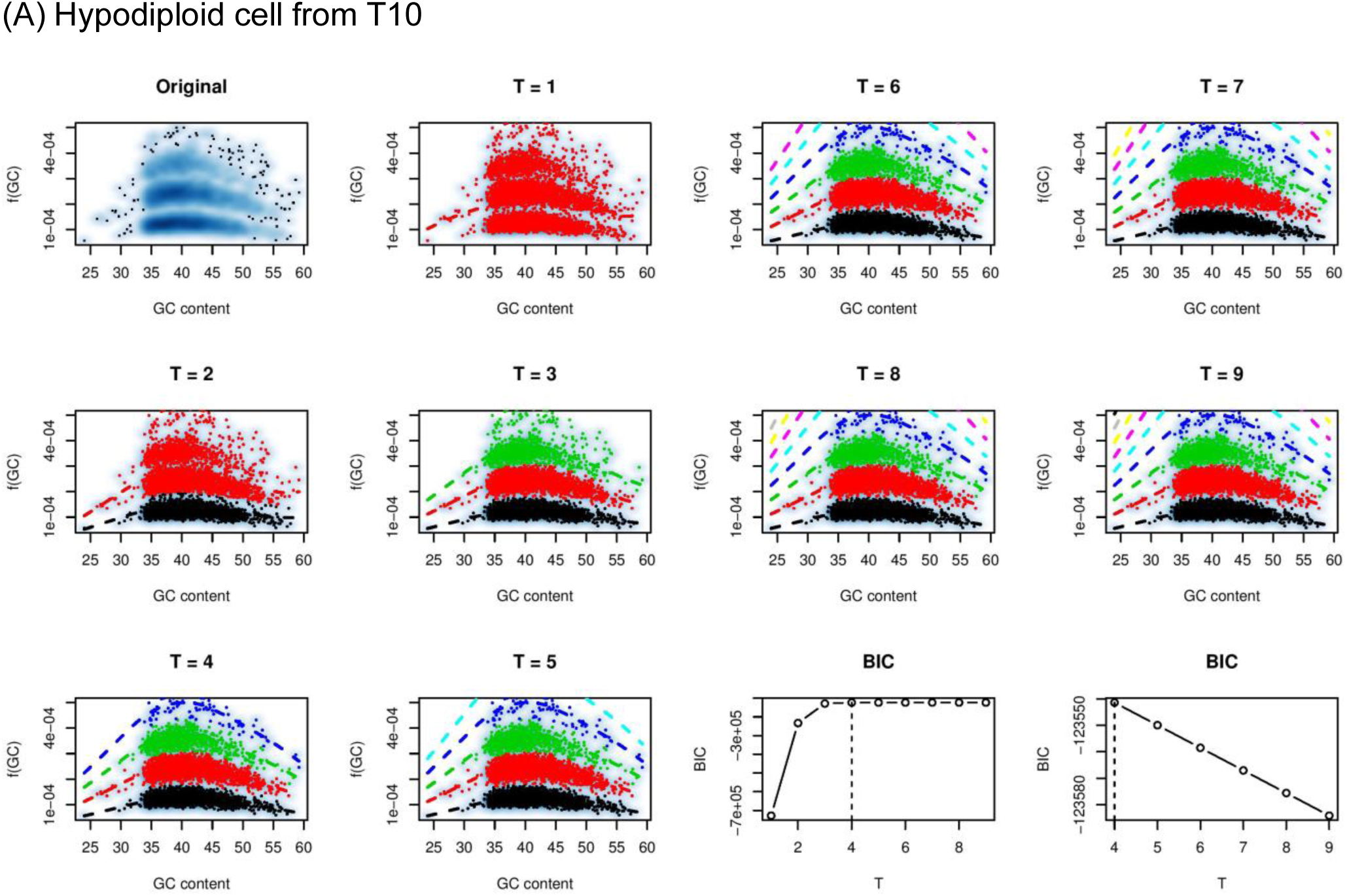
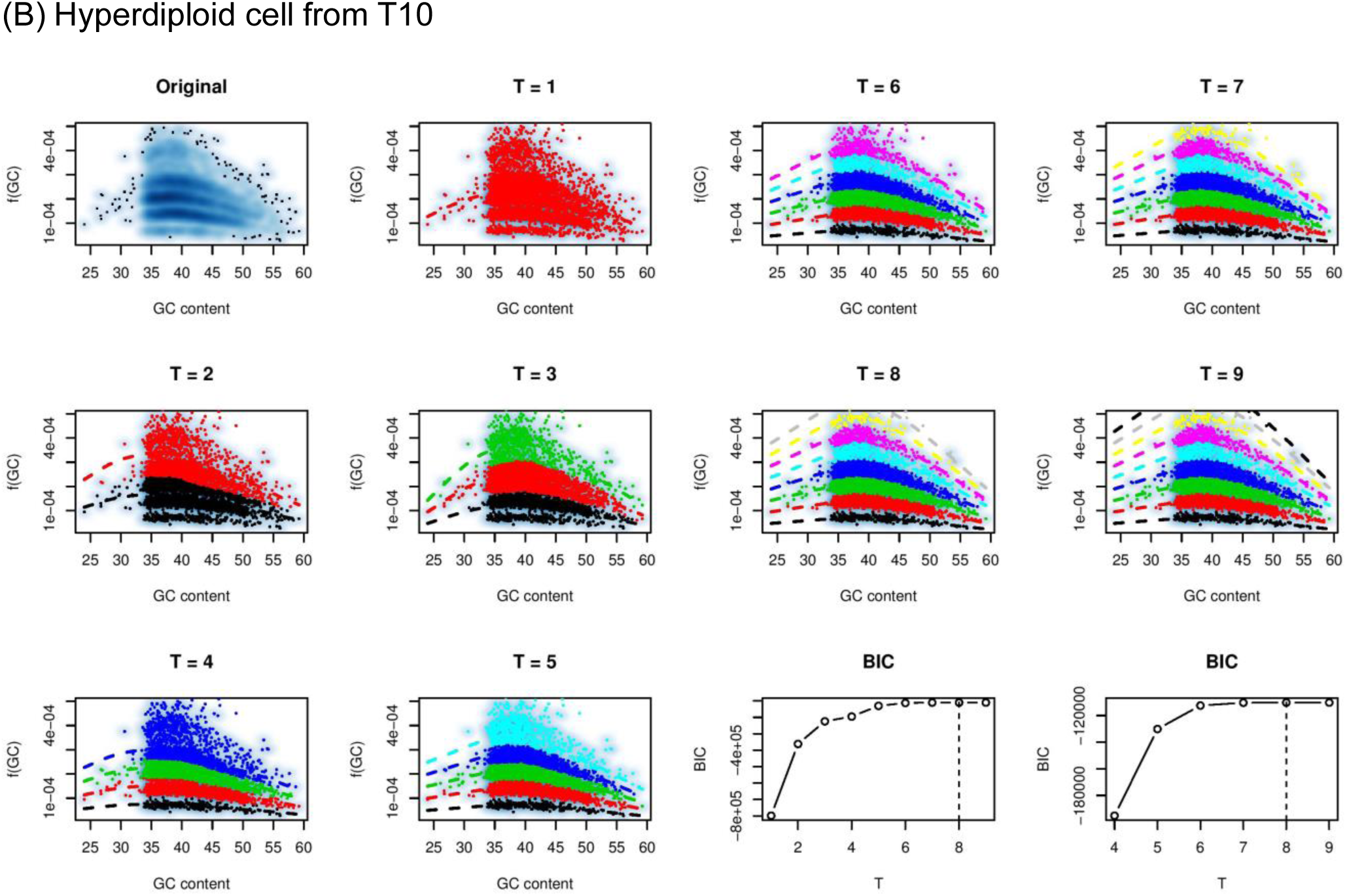
EM algorithm for estimating GC content bias with BIC for model selection. (A) Fitting of (A) hypodiploid and (B) hyperdiploid cell from T10 are shown. The “original” panel shows the f(GC) patterns through the iterative estimation procedure, in which the EM algorithm is embedded.

**Figure S2.**
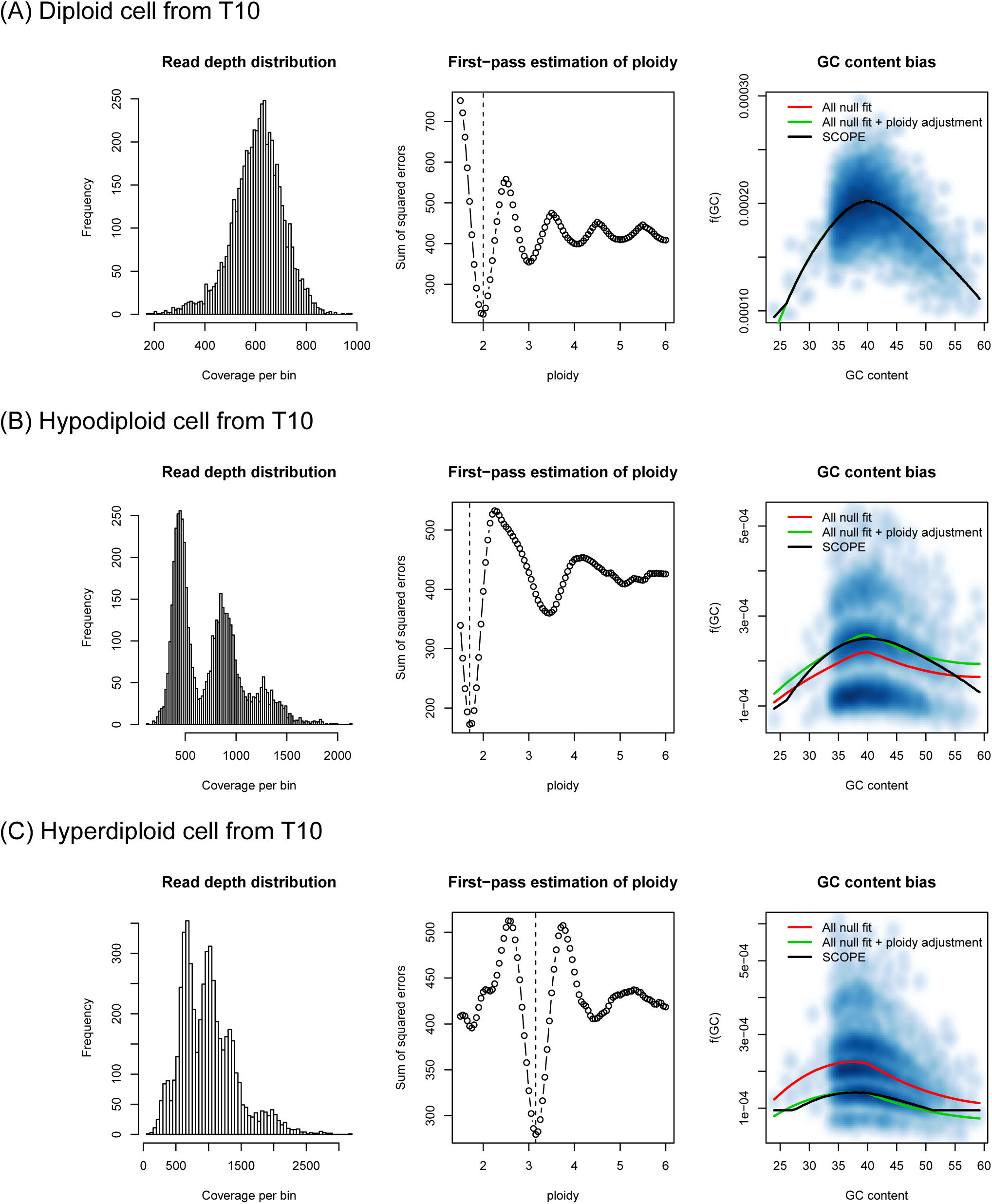
Pre-estimation of ploidy for EM initialization. Cell ploidy is pre-estimated through a first-pass normalization run, where the cost function measuring the difference between the scaled copy number and its integer rounded copy number is minimized.^5^ The empirical cost function shows a typical damped sine wave.^13^ The pre-estimated ploidy is only used for the EM initialization to ensure fast convergence and to avoid local optima.

**Figure S3.**
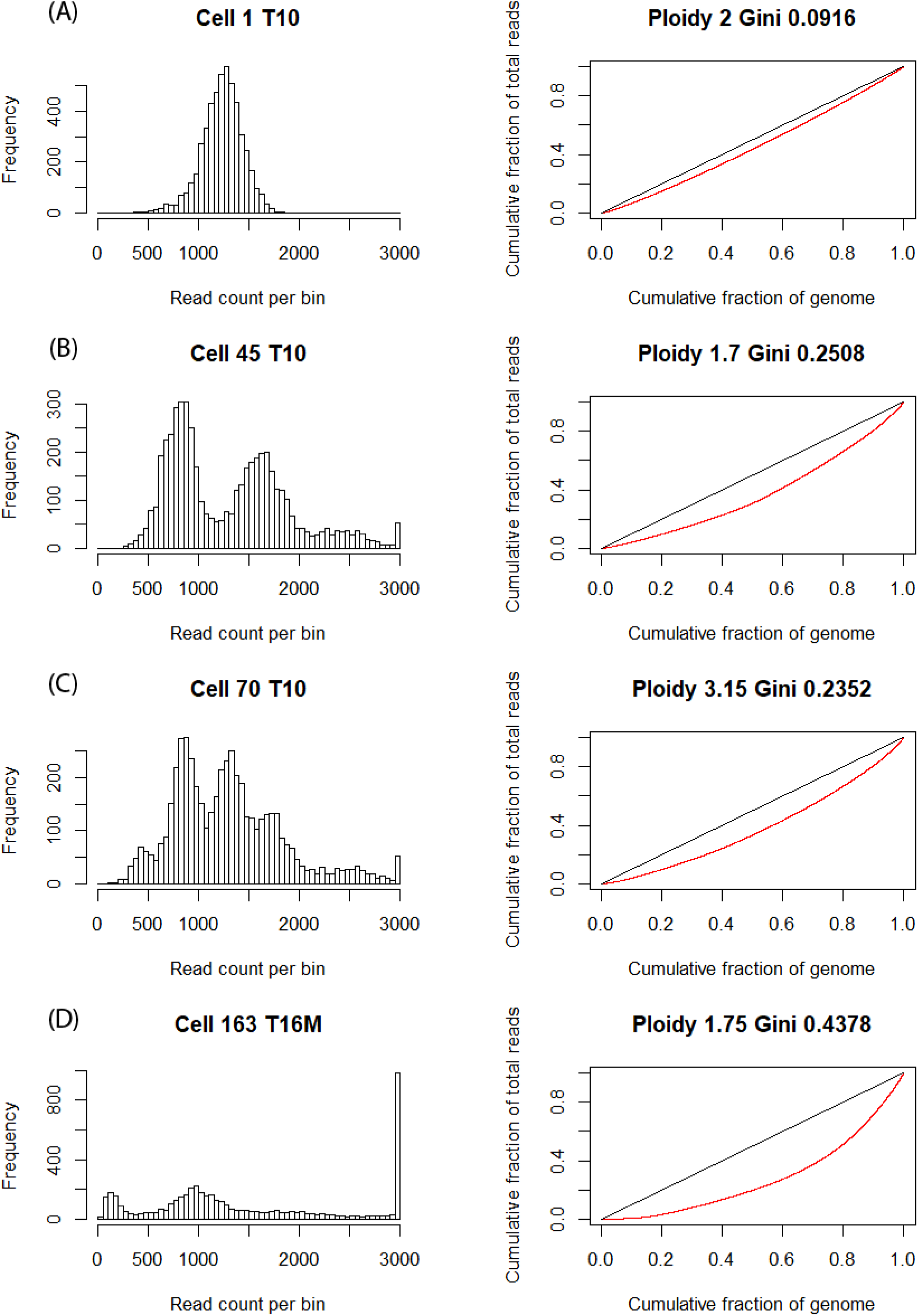
Gini coefficients for assessment of coverage uniformity and identification of normal cells. Depth of coverage distribution and Lorenz curve for (A) a diploid cell, (B) a hyperdiploid cell, and (C) a hypodiploid cell are shown. Aneuploid cells have uneven read depth distribution due to copy number aberrations and thus high Gini indices. (D) Cell with extremely high Gini coefficient, due to, e.g., failed library preparation, is excluded from downstream analysis.

**Figure S4.**
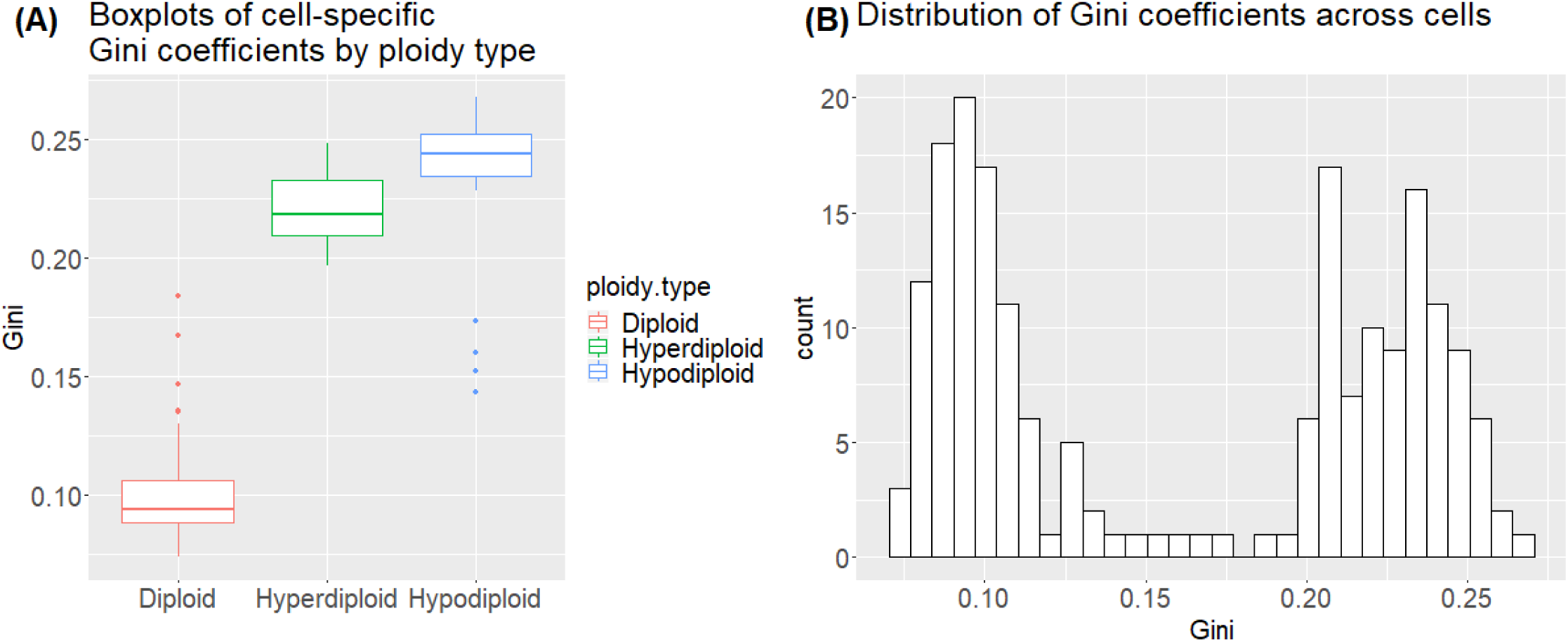
Gini coefficients serve as good metrics for identification of normal cells. (A) Ploidy information for each cell has been previously reported. Hyperdiploid and hypodiploid cells have higher Gini indices compared to diploids. (B) Histogram of Gini coefficients across all cells. A cutoff of 0.12 was used to identify normal controls for SCOPE. We show empirically that SCOPE does not need to include every true normal cell as negative control – as few as 10 to 20 cells are needed for accurate estimation of the noise terms.

**Figure S5.**
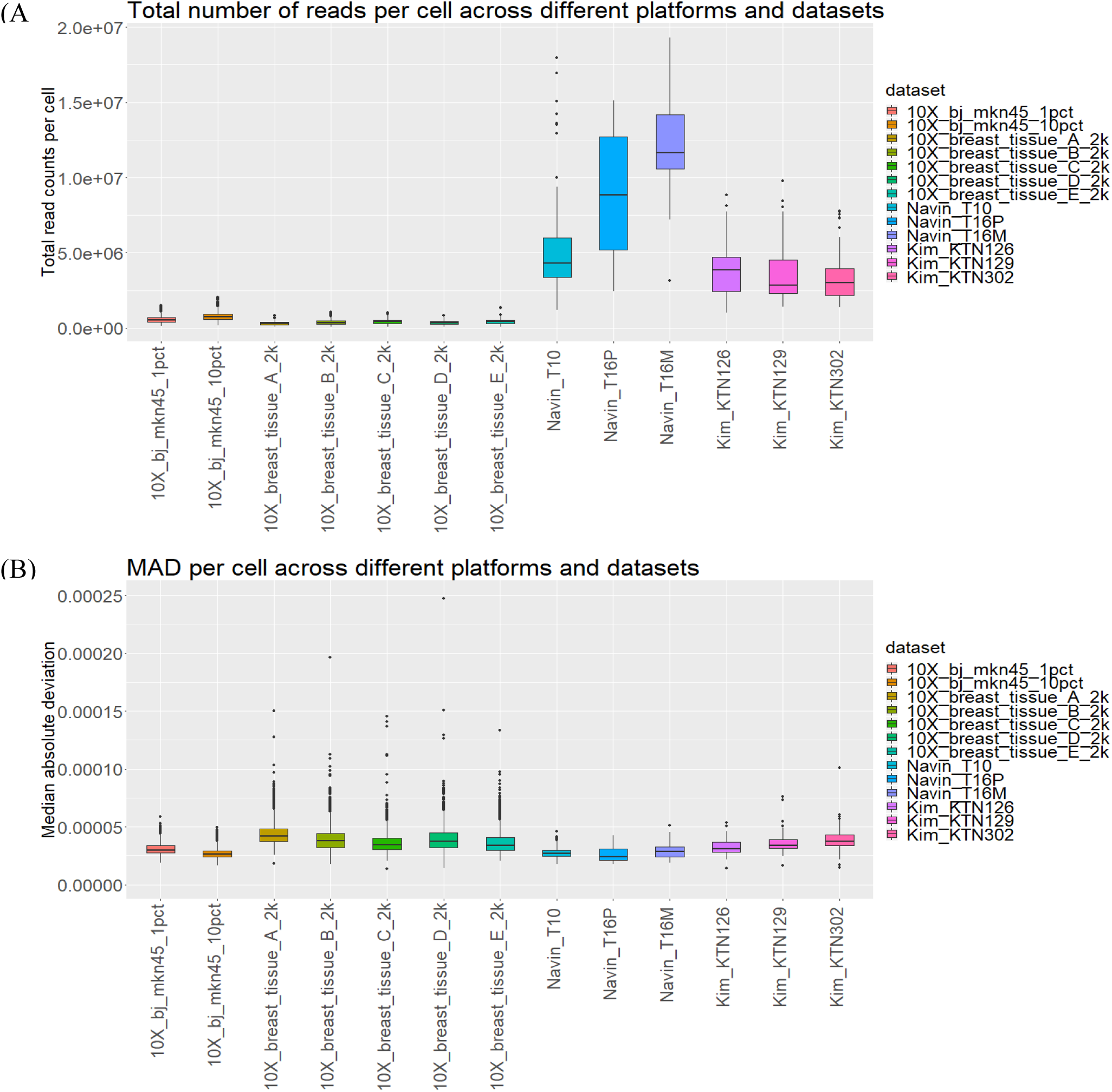
Assessment of read depth and coverage uniformity across different platforms and datasets. (A) Total number of reads per cell across different platforms and datasets. There is a tradeoff between the number of cells and the number of reads per cell. Total read count per cell of the 10X genome amplification (WGA) data (mean 5,350,389 3,835,852 s.d.). (B) Median absolute deviation (MAD) of all pairwise differences in read counts between neighboring bins for each cell across different platfo_Y_rm_/(_s_N_a_*β*_nd_)_datasets. Read counts are pre-normalized for library size factor and bin-specific factor, conventional WGA data (mean 3.25e-5 ± 9.03e-6 s.d.) show trivial differences in MAD between studies. However, more cell outliers with extreme coverage uniformity are observed from the 10X Genomics pipeline and are removed from downstream analysis.

**Figure S6.**
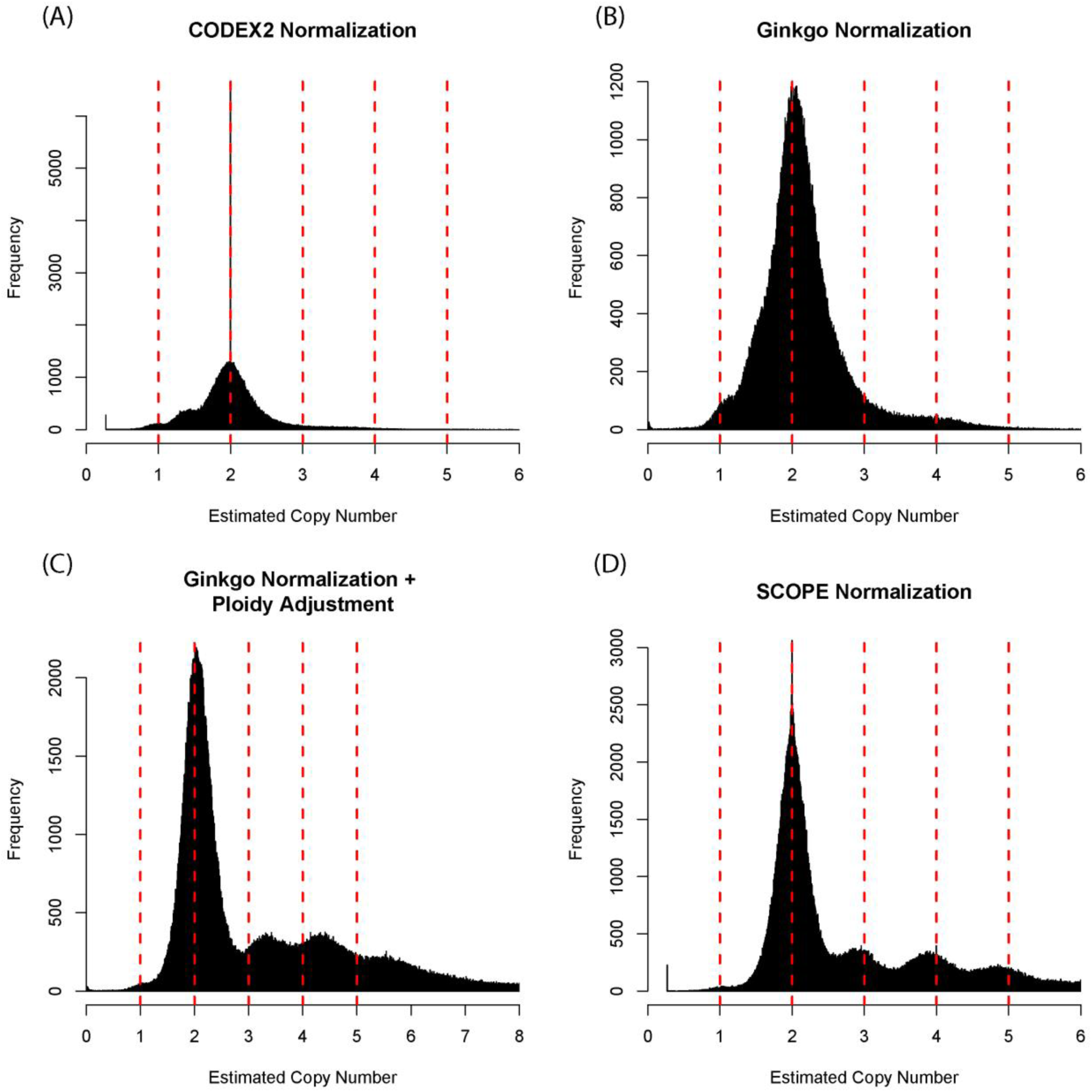
Normalization results for scDNA-seq data of monogenomic tumor T16. Normalized *z*-zscores are shown for (A) CODEX2 as 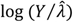, (B) Ginkgo, which centers bin counts of each cell at 1 and further fits a locally weighted linear regression (LOESS curve) to adjust for GC content, (C) Ginkgo with post hoc ploidy adjustment, which follows to be scaled by the optimal copy-number multiplier, and
(D) SCOPE as 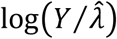.

**Figure S7.**
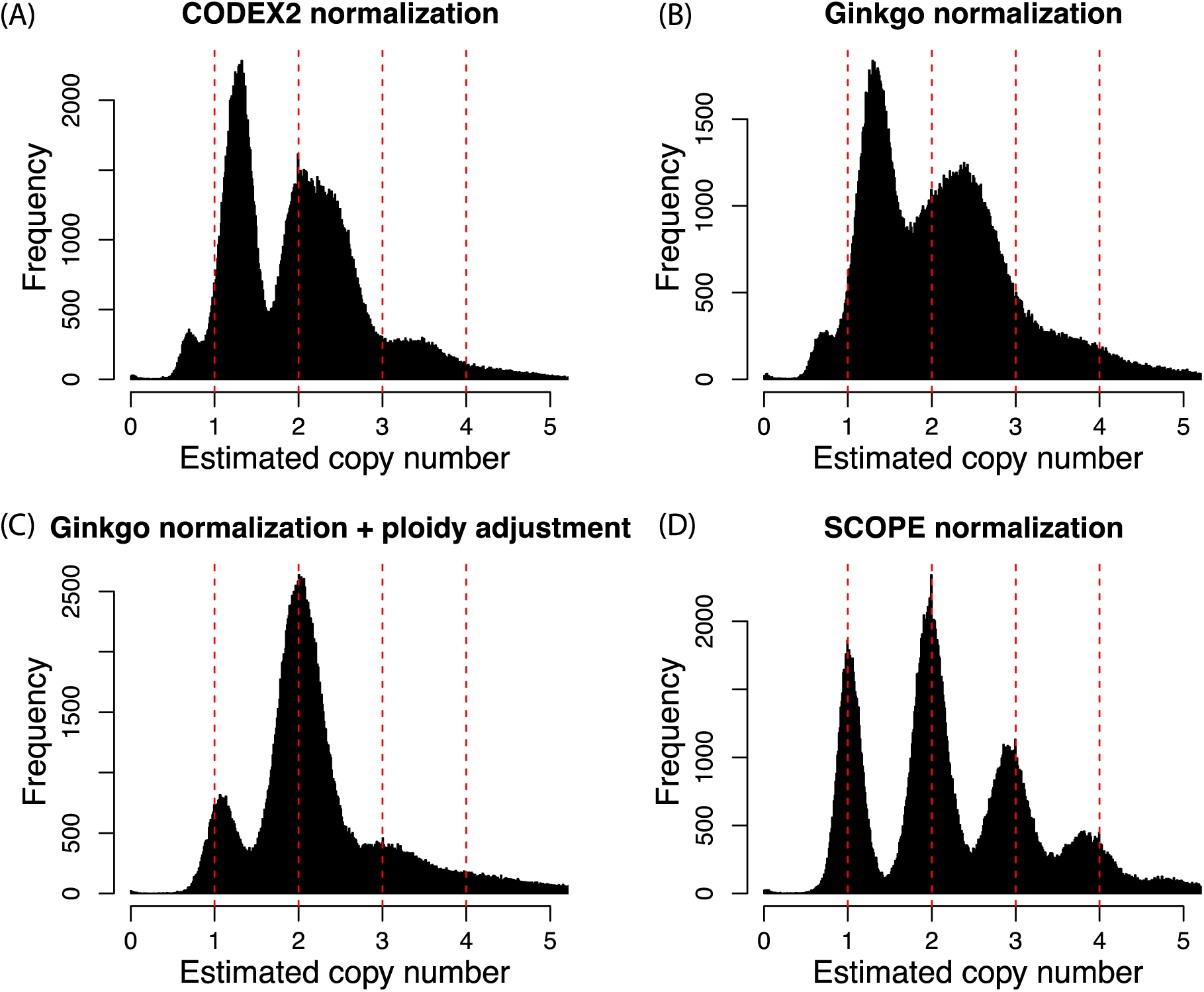
Normalization results for scDNA-seq data of polygenomic tumor T10. Normalized *z*-scores are shown for (A) CODEX2 as 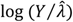, (B) Ginkgo, which centers bin counts of each cell at one and further fits a locally weighted linear regression (LOESS curve) to adjust for GC content, (C) Ginkgo with post hoc ploidy adjustment, which follows to be scaled by the optimal copy-number multiplier, and (D) SCOPE as 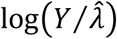. Only tumor cells are shown in the histogram. Distribution across all cells are shown in Figure 2.

**Figure S8.**
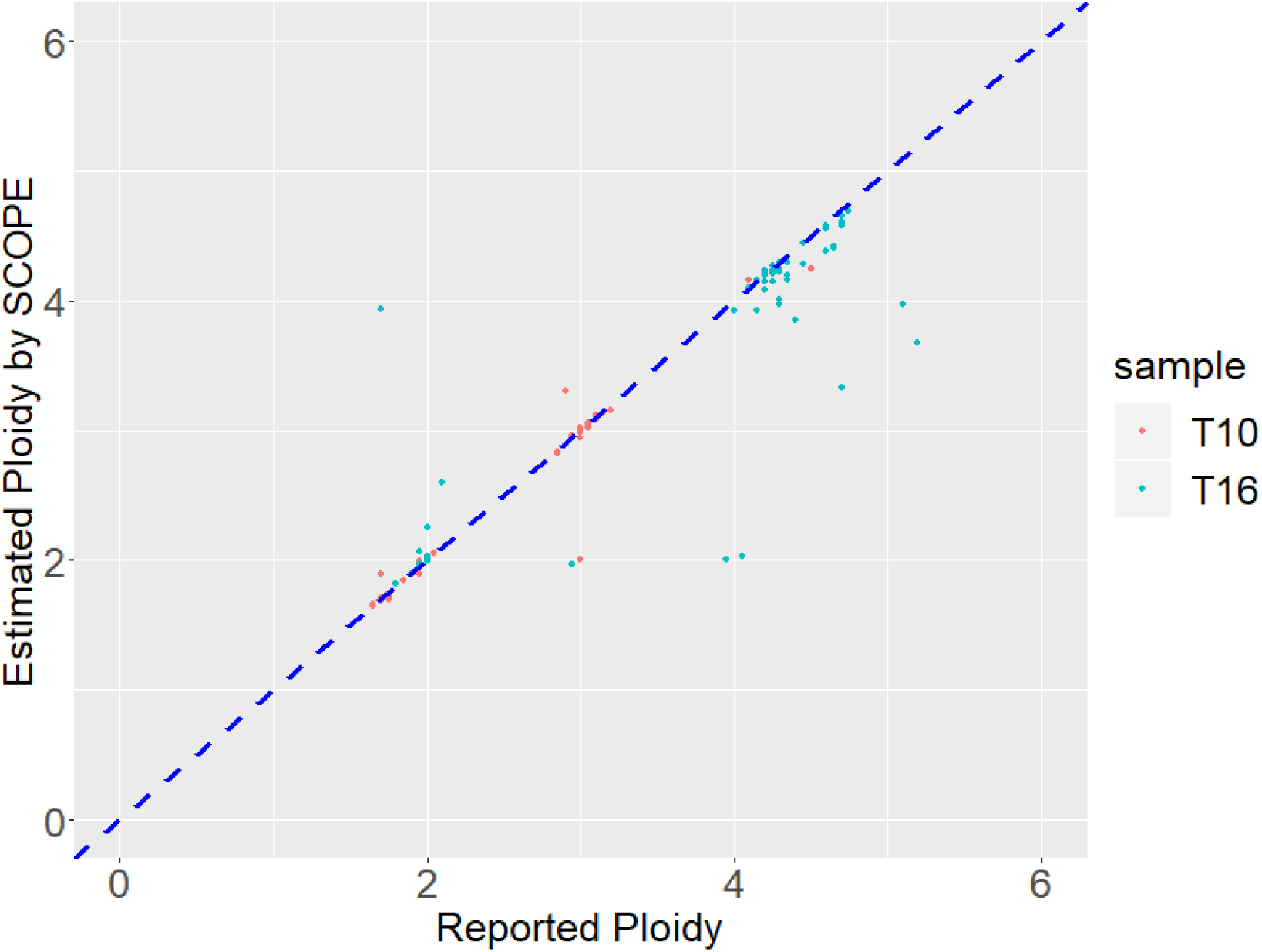
Correlation between ploidy estimated by SCOPE and ploidy previously reported. SCOPE automatically returns ploidy estimates, which are highly concordant with previous reports based on single-cell sorting, with Pearson correlation of 0.982 and 0.929 in T10 and T16, respectively.

**Figure S9.**
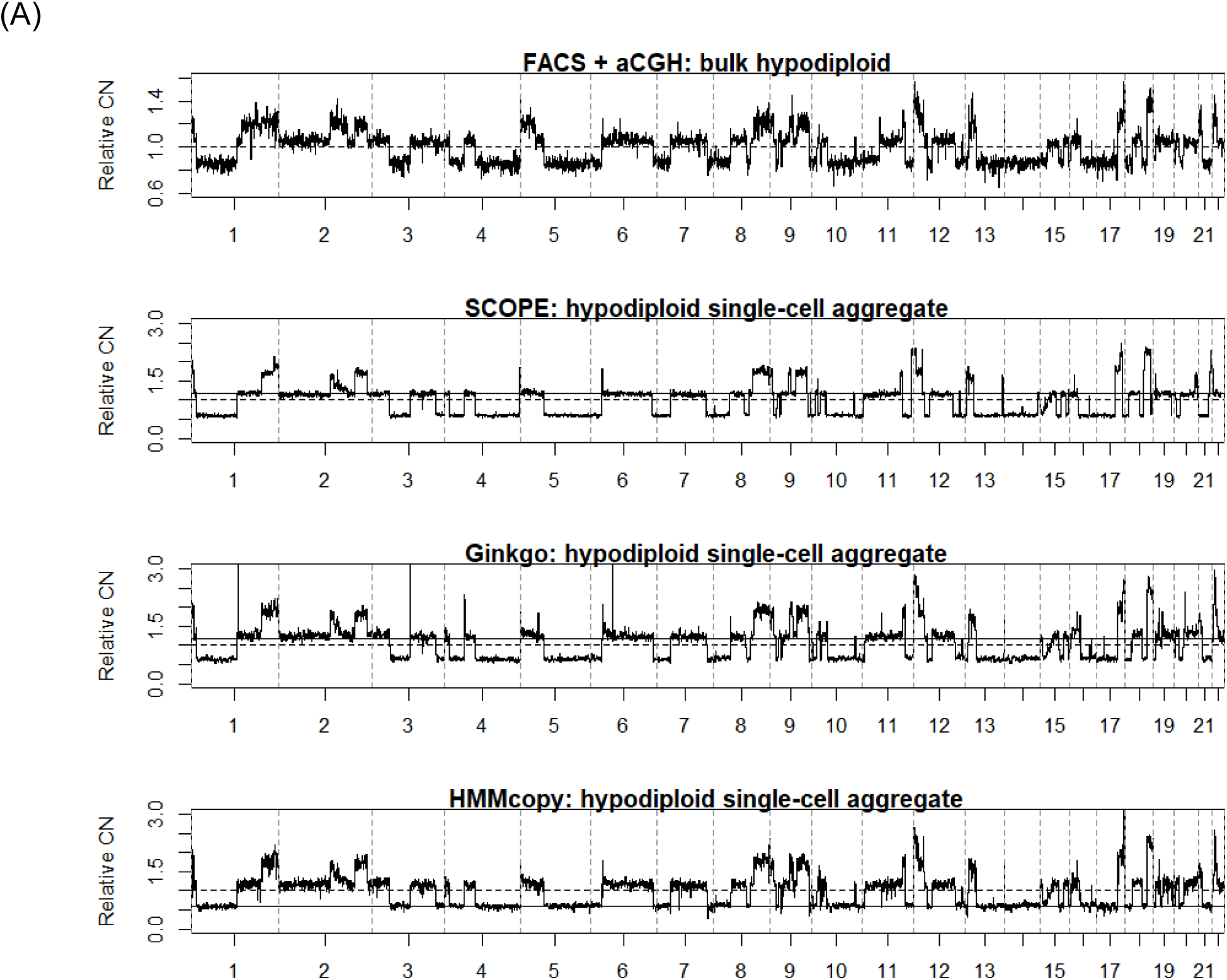
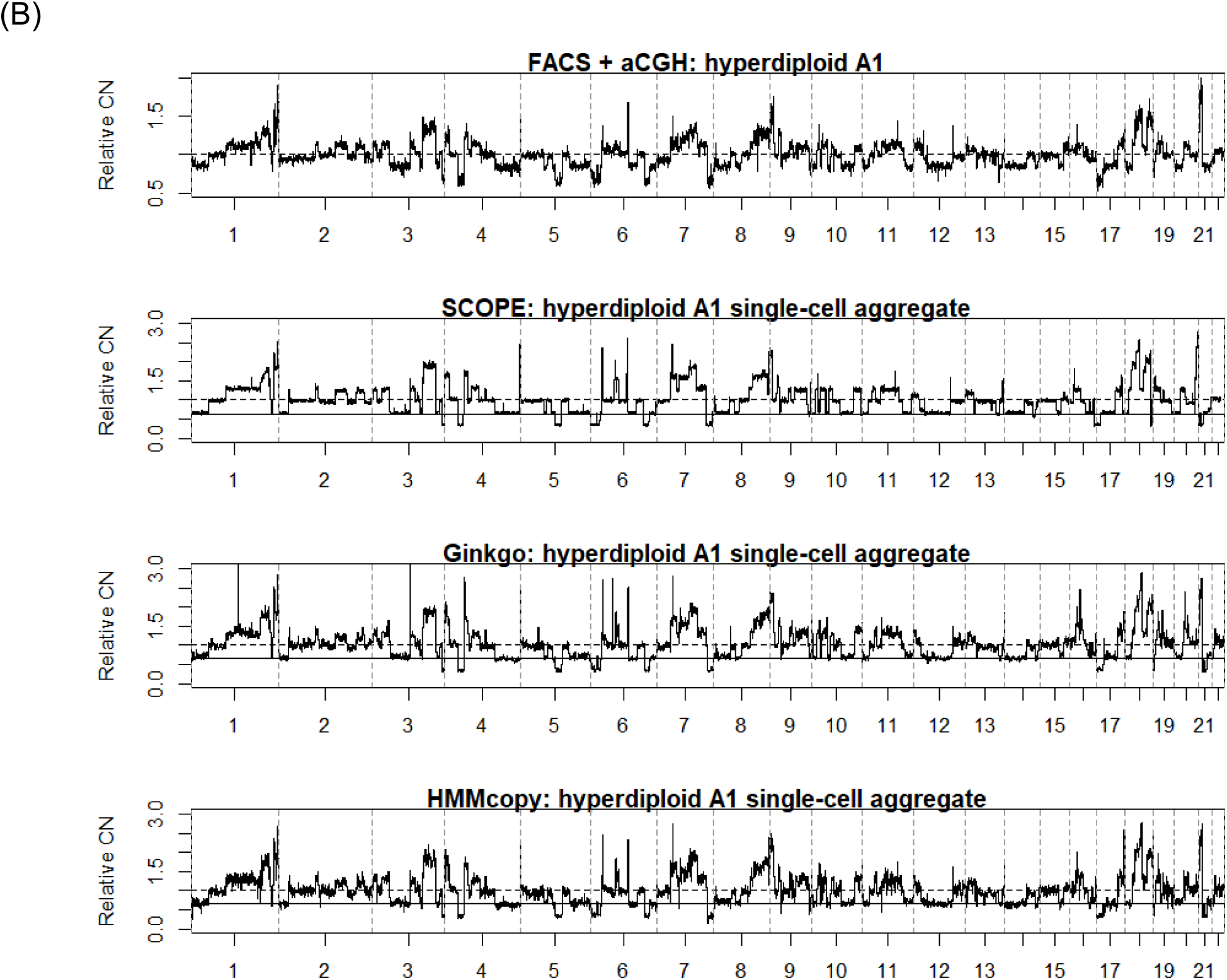
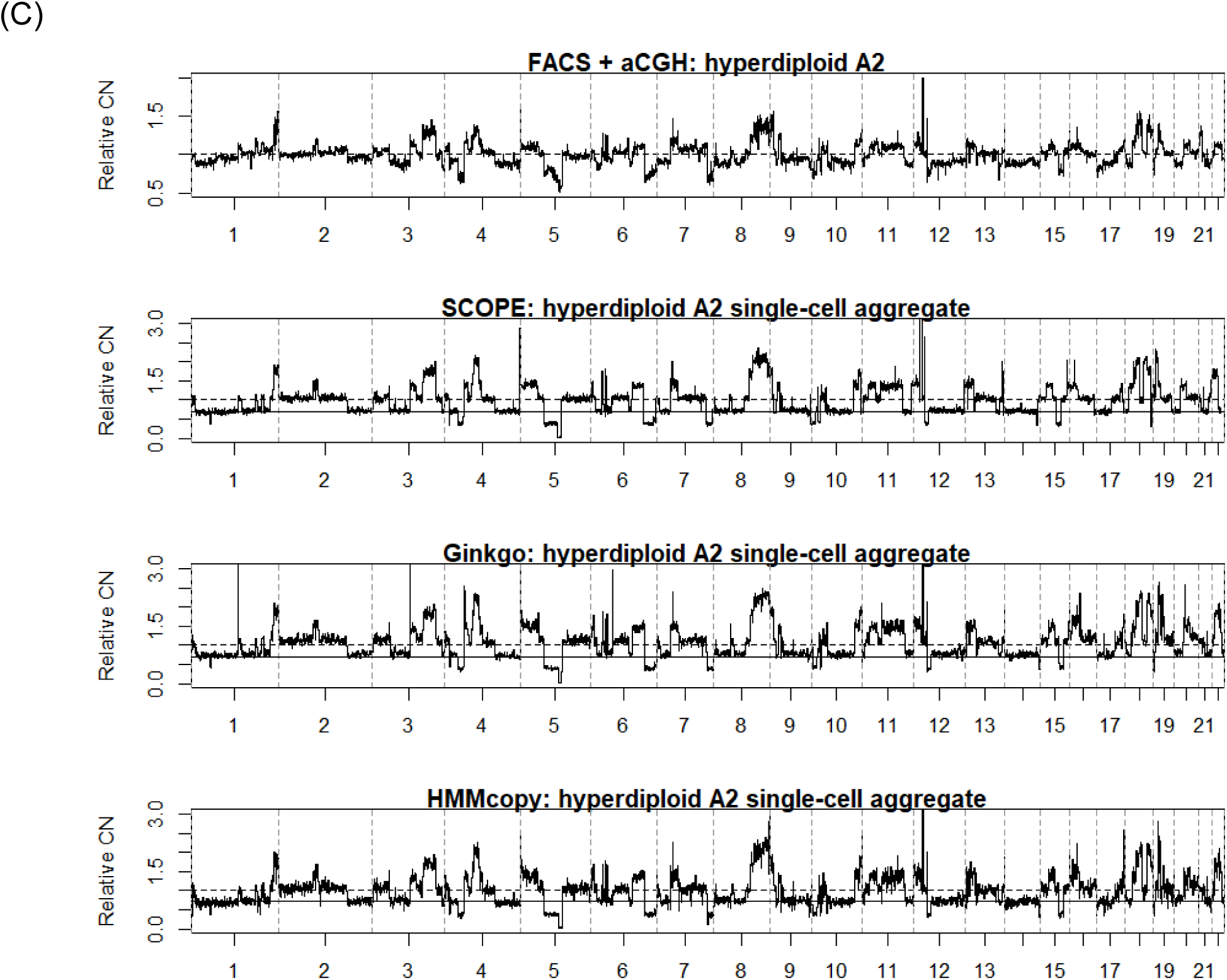
Orthogonal validation of single-cell copy number profiles by aCGH of purified bulk samples by FACS. FACS reveals (A) hypodiploid, (B) hyperdiploid A1, and (C) hyperdiploid A2 cancer subclones within breast cancer patient T10. This is followed by aCGH on the sorted bulk samples to profile copy number aberrations, which are used as ground truth. Relative copy number (CN) estimates from the three subpopulations are shown in (A)-(C) for aCGH, SCOPE, Ginkgo, and HMMcopy.

**Figure S10.**
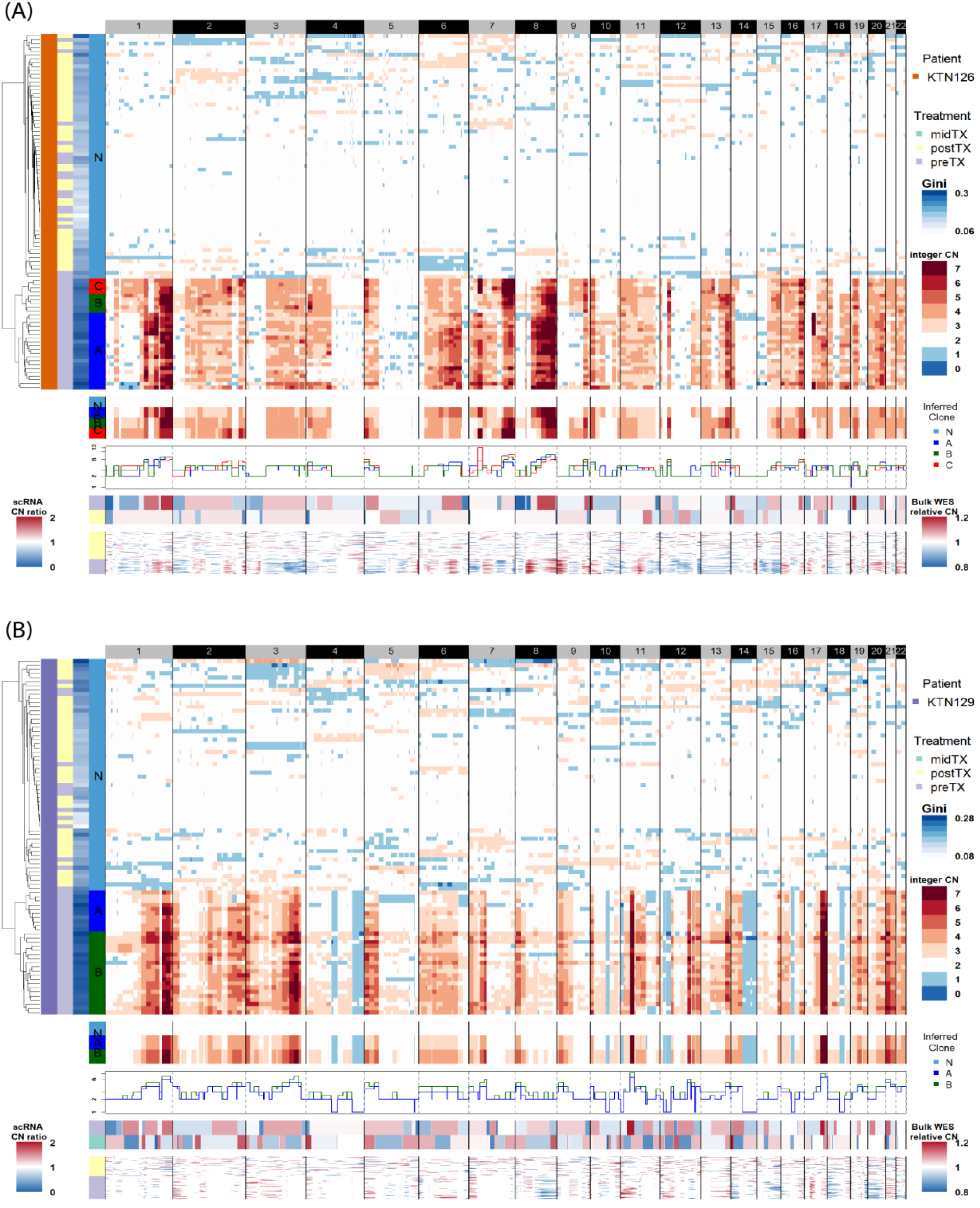
Copy-number profiles of triple-negative breast cancer patients, inferred by scDNA-seq, bulk WES, and scRNA-seq. (A) patient KTN126; (B) patient KTN 129. Upper three panels show heatmap of inferred copy-number profiles by SCOPE with cells clustered by hierarchical clustering, heatmap and stairstep plot of consensus copy number profiles for each cell subpopulation (N for normal cell, A, B, and C for cancer subclones). Lower two panels show copy number profiles inferred by WES and scRNA-seq, which are further used to validate CNV calls by SCOPE.

**Figure S11.**
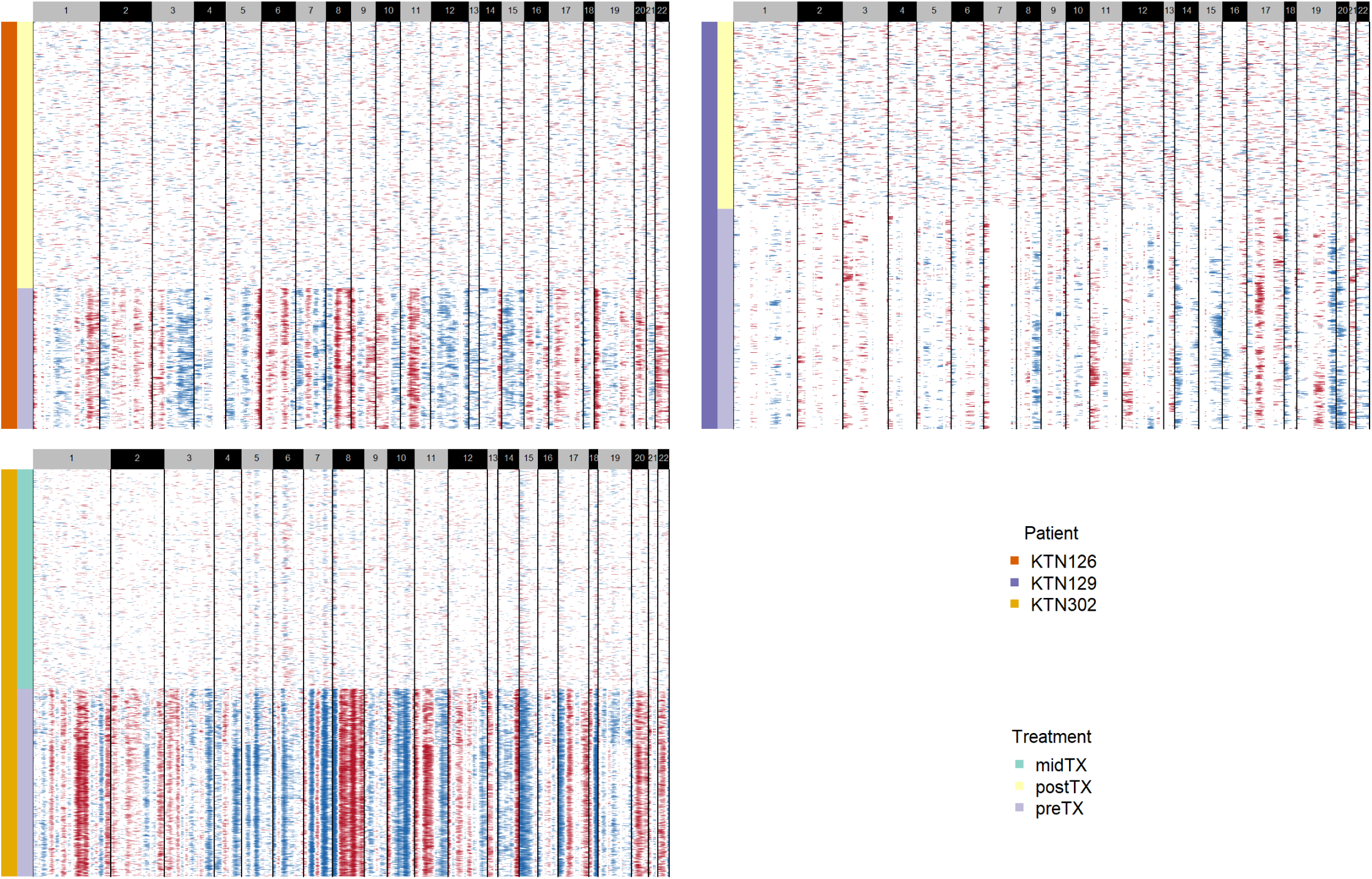
CNV profiling by scRNA-seq using a sliding-window approach. InferCNV^14^ is applied to scRNA-seq data of three clonal extinction patients (KTN126, KTN129, and KTN302) for CNV profiling, which is further used to validate the CNV calls by scDNA-seq. SALMON^15^ was used to get the matrix of transcript per million (TPM) and we further excluded genes that were detected in less than 30% of cells, resulting in 3,781 genes per cell that were used for copy number estimation.

**Figure S12.**
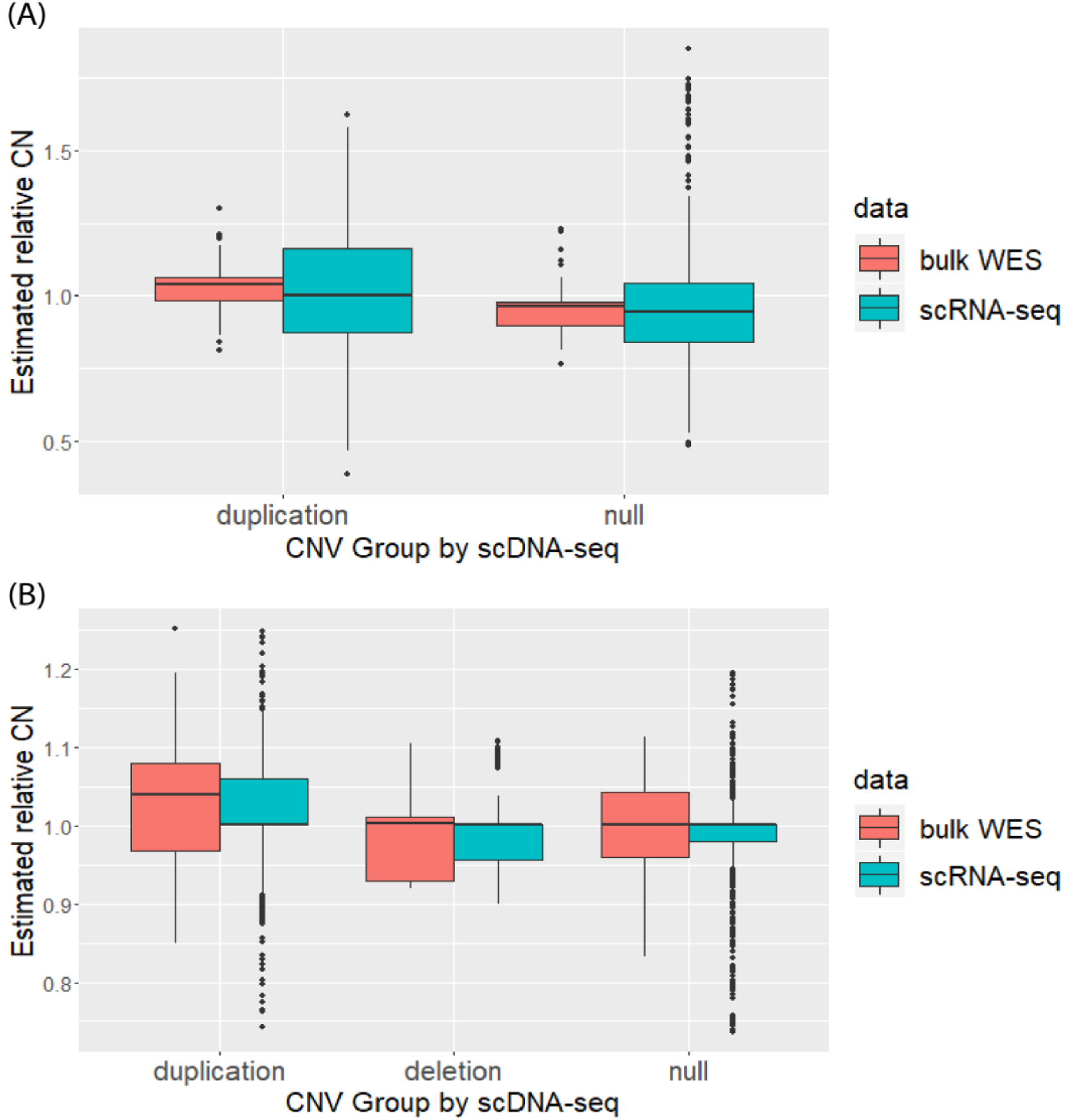
Orthogonal validations using copy number profiles by WES and scRNA-seq. (A) patient KTN126; (B) patient KTN 129. Each genomic segment is categorized as amplification (amp), deletion (del), or copy-number neural (null) based on the copy number profiles returned by SCOPE. The boxplot shows the estimated relative copy numbers by bulk WES and scRNA-seq, which are higher in amplified regions and lower in deleted regions, compared to the null regions. Due to low resolution and high signal-to-noise ratio by WES and scRNA-seq, there is no perfect separation between the different copy number states. Furthermore, for scRNA-seq and bulk WES, the relative copy numbers are estimated comparing to a sample-specific baseline. For cases where the tumor cells are hyperdiploid/hypodiploid, the baseline is shifted up/down making the true null regions have relative copy numbers less/greater than one.

**Figure S13.**
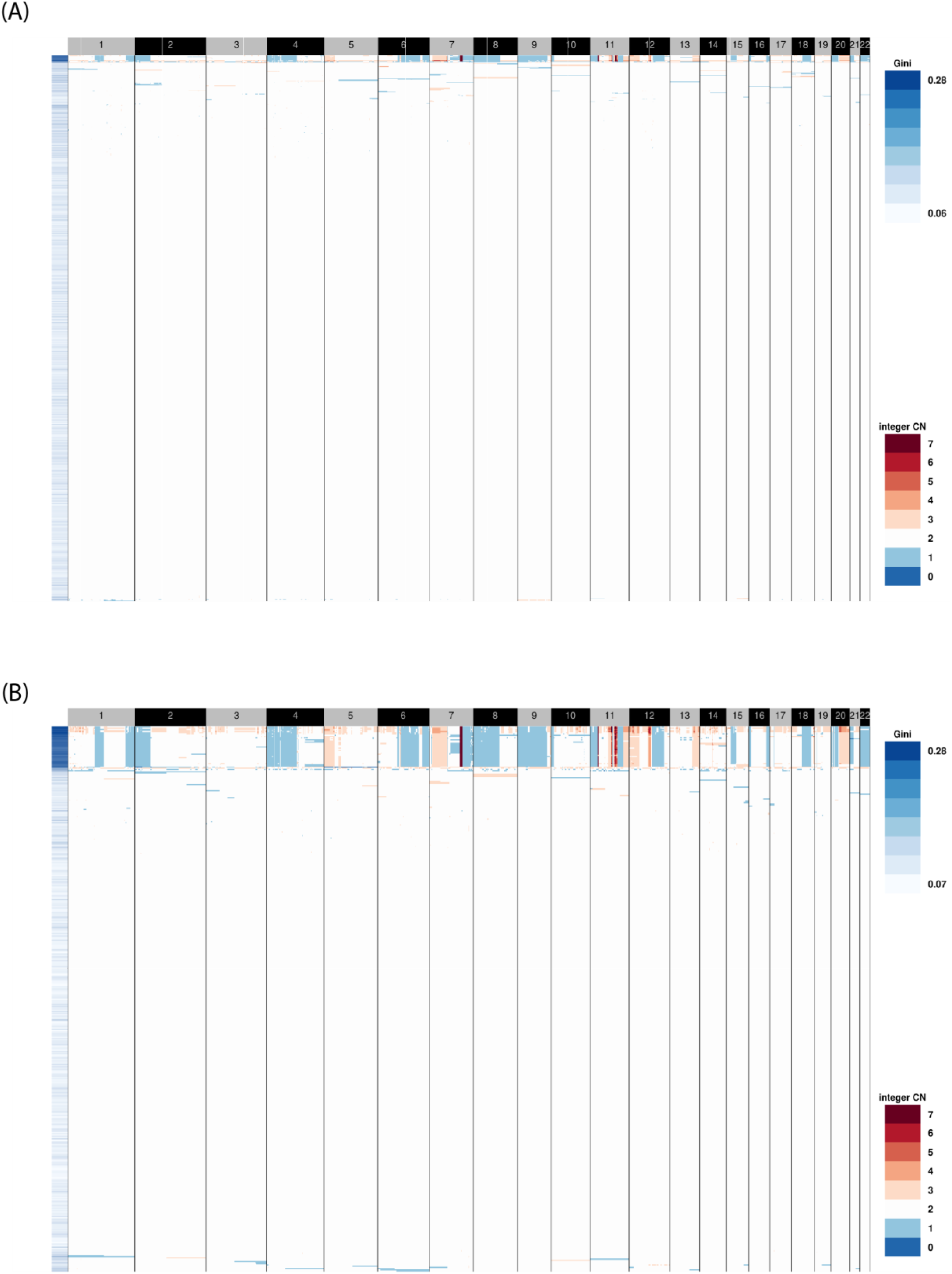
Copy-number profiles by SCOPE of the 10X Genomics cancer spike-in datasets. (A) 1% and (B) 10% gastric cancer cell lines were mixed with normal fibroblast cell lines. Heatmaps after cross-sample segmentation are shown. SCOPE successfully identifies the tumor cells from the background of normal.

**Figure S14.**
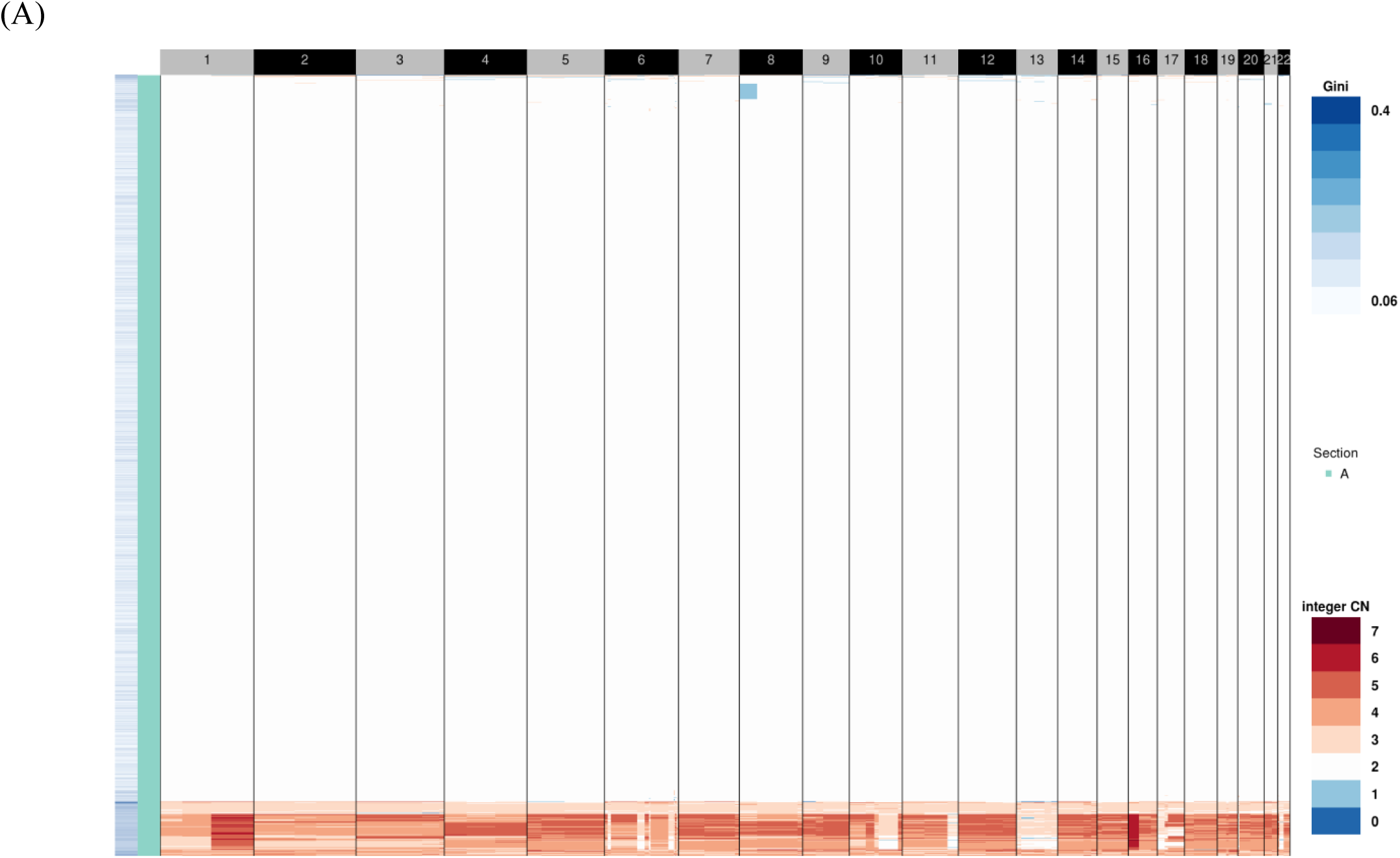
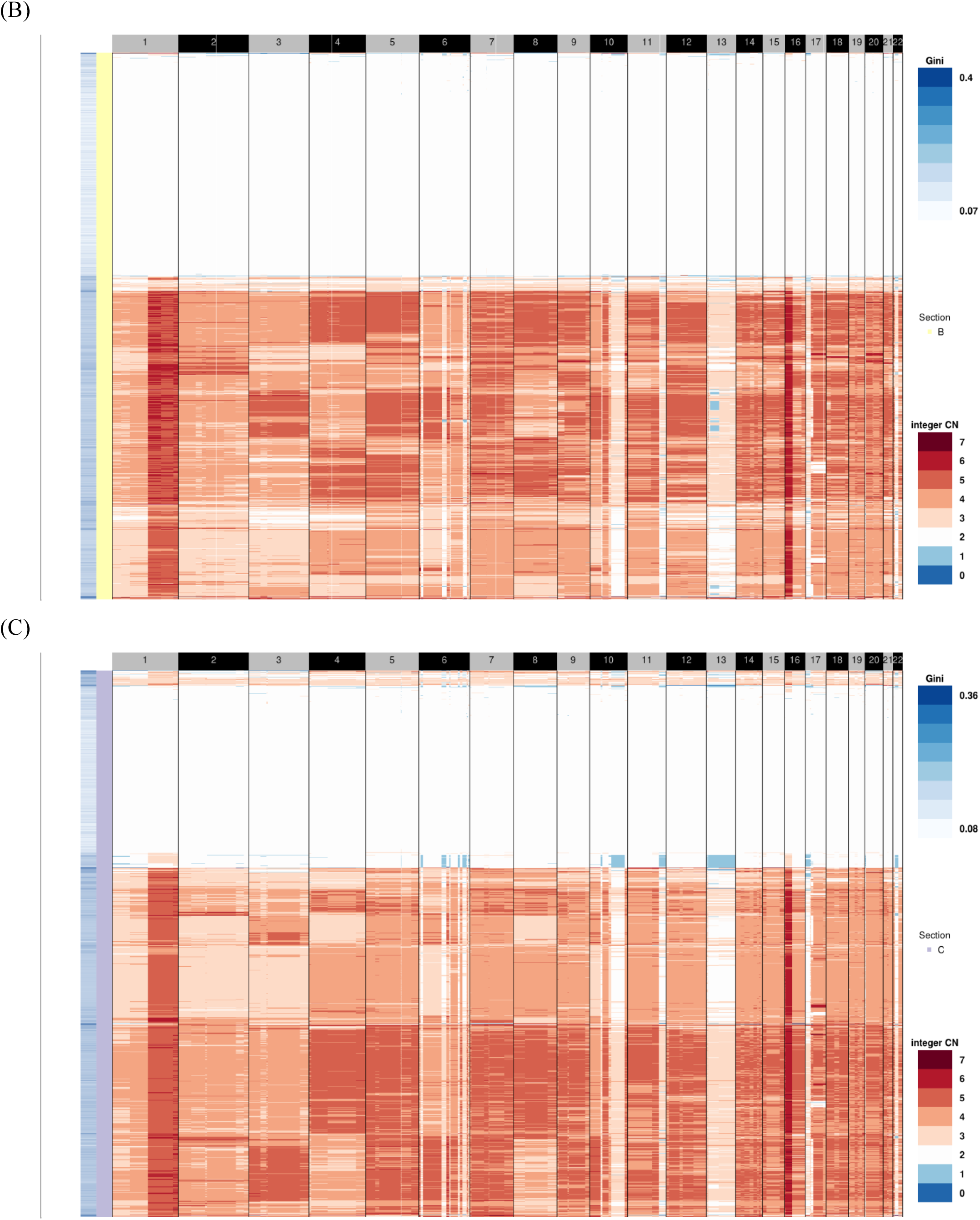
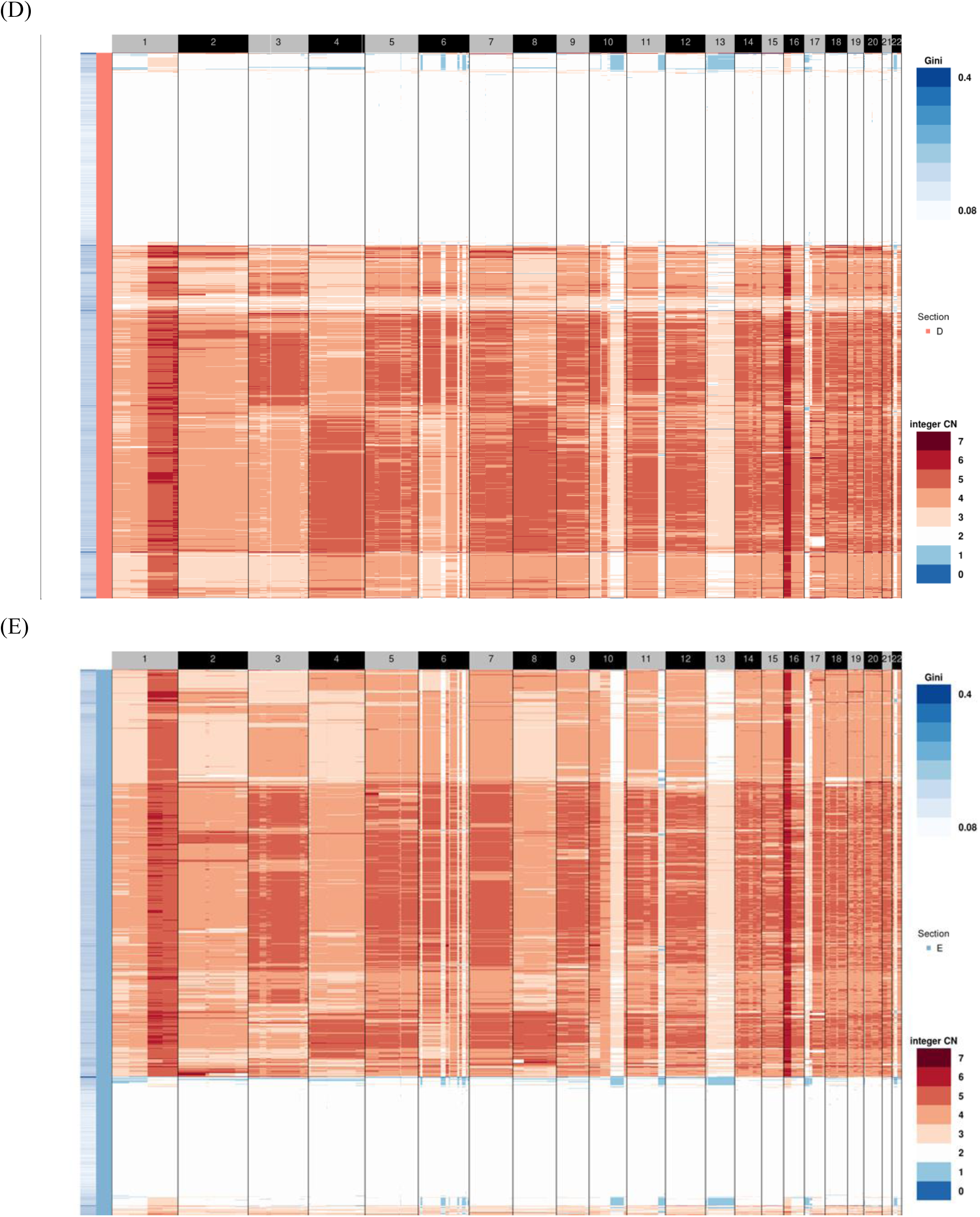
Single-cell copy number profiles across five breast cancer dissections. Heatmaps of inferred copy-number profiles by SCOPE across five tumor dissections after cross-sample segmentation are shown, where each row depicts the whole genome of a single cell. A gradient of diploid cells contaminating the tumor is observed from dissection A (shown in panel A) to dissection E (shown in panel E).

**Figure S15.**
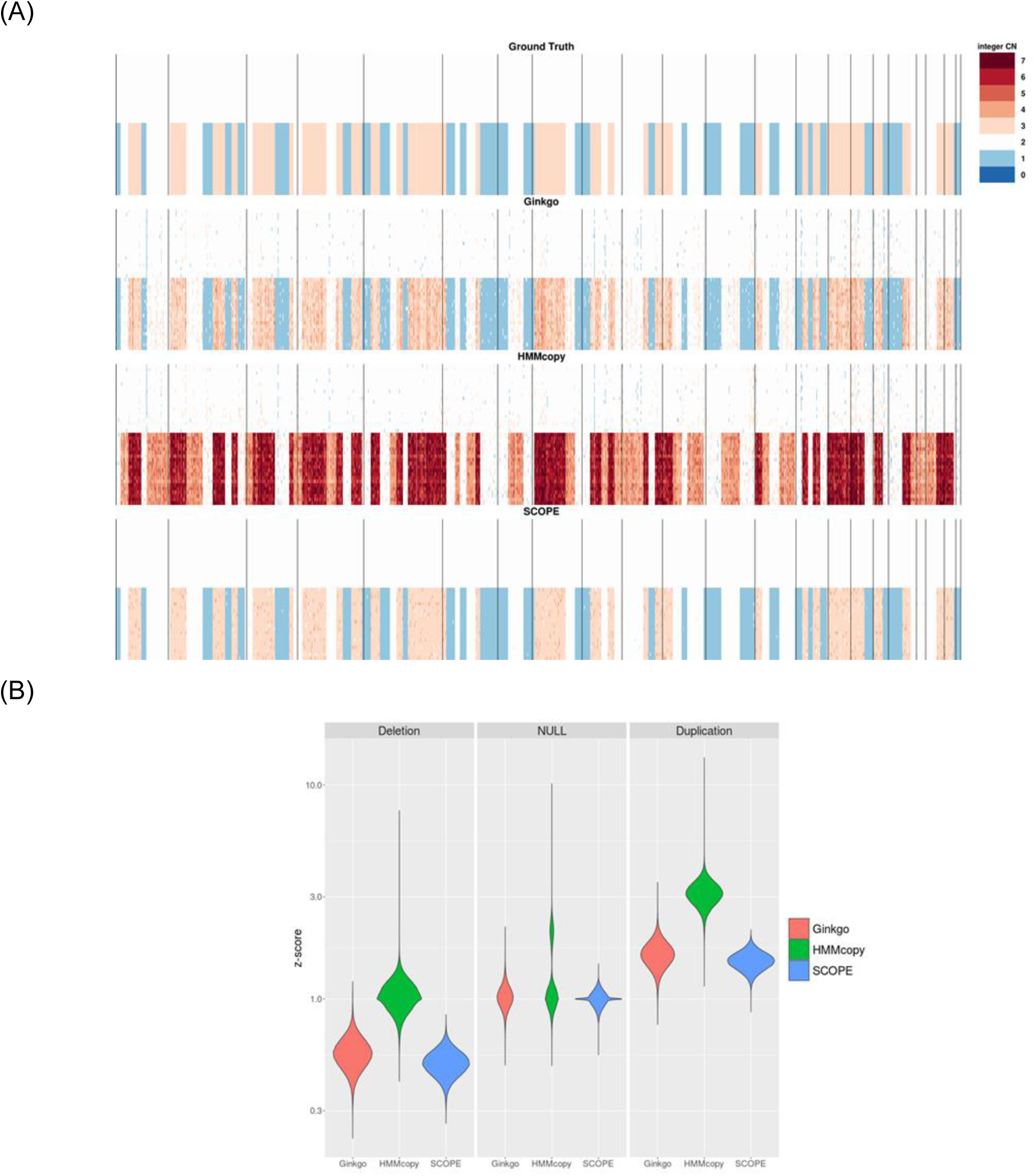
Normalization results from spike-in studies with copy number states 1, 2, and 3. Copy number signals were manually added to the depth of coverage data from the null genomic regions of patient T10. (A) Heatmaps of ground truth, Ginkgo’s normalization result, HMMcopy’s normalization result, and SCOPE’s normalization result. (B) Normalized z-scores for the true deletions, null regions, and duplications. HMMcopy failed to recover the true copy number states with a genome-wide copy number inflation.

**Figure S16.**
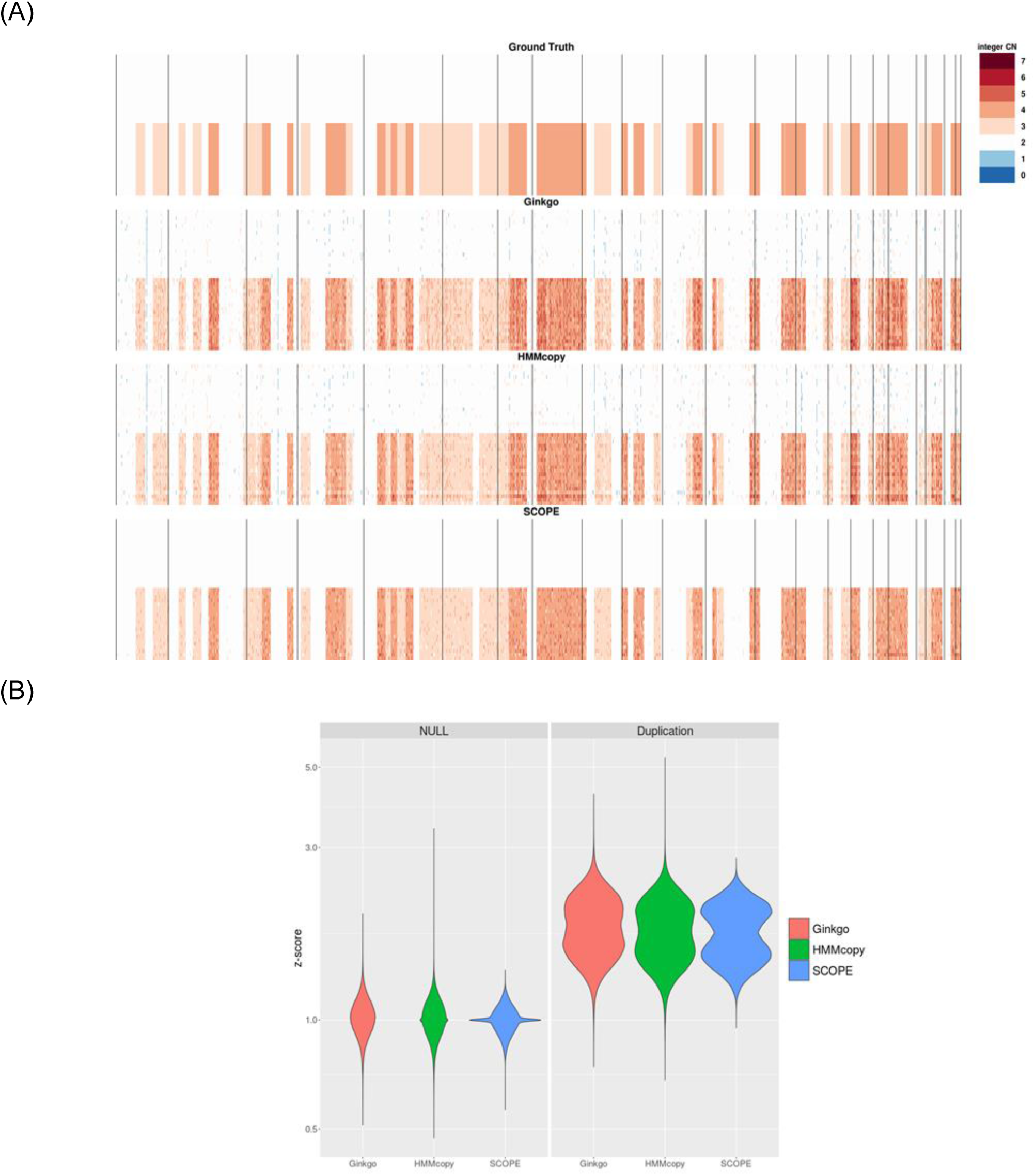
Normalization results from spike-in studies with copy number states 2, 3, and 4. Copy number signals were manually added to the depth of coverage data from the null genomic regions of patient T10. (A) Heatmaps of ground truth, Ginkgo’s normalization result, HMMcopy’s normalization result, and SCOPE’s normalization result. (B) Normalized z-scores for the true deletions, null regions, and duplications.

**Figure S17.**
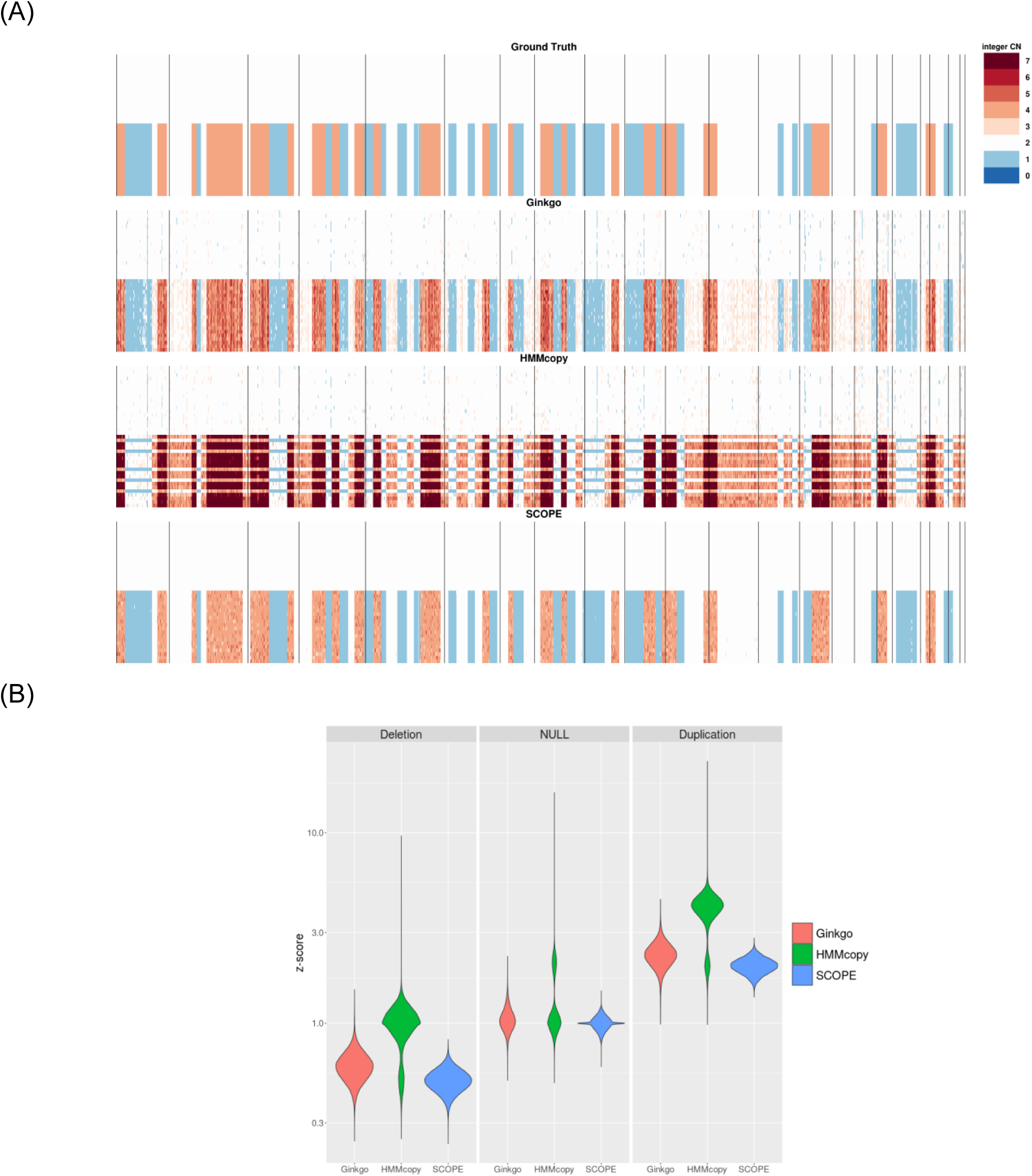
Normalization results from spike-in studies with copy number states 1, 2, and 4. Copy number signals were manually added to the depth of coverage data from the null genomic regions of patient T10. (A) Heatmaps of ground truth, Ginkgo’s normalization result, HMMcopy’s normalization result, and SCOPE’s normalization result. (B) Normalized z-scores for the true deletions, null regions, and duplications. HMMcopy failed to recover the true copy number states with a genome-wide copy number inflation.

**Figure S18.**
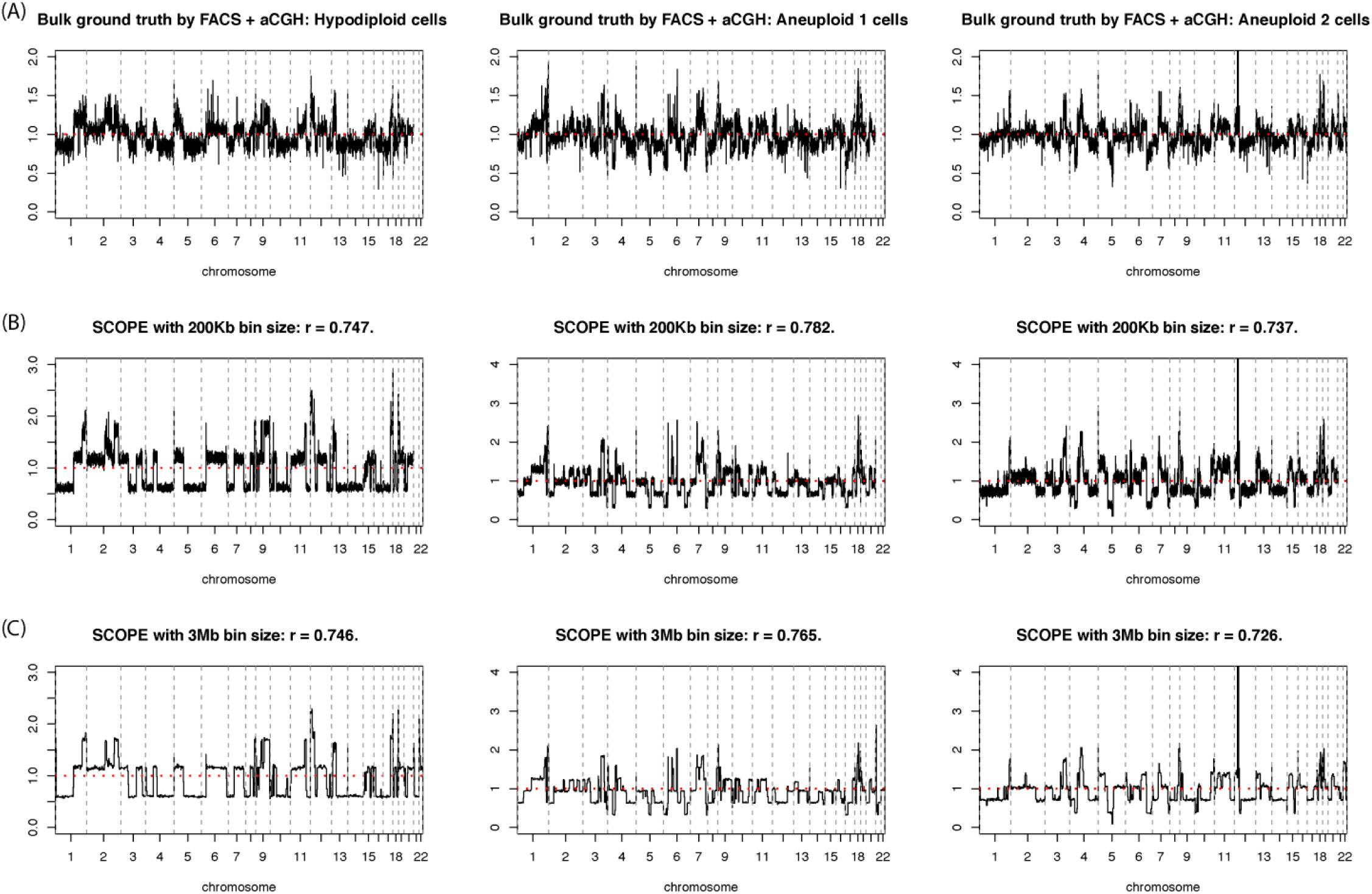
Copy number estimation by SCOPE is robust to bin size selection. SCOPE, by its default, adopts a fixed bin size of 500Kb. Empirical evidence using ground truth by FACS and aCGH suggest that performance of SCOPE is invariant to bin size when determining large chromosomal level copy number changes. (A) Ground truth by aCGH on FACS purified bulk samples. (B) SCOPE’s performance using 200Kb bin size. (C) SCOPE’s performance using 3Mb bin size. Smaller bin size leads to higher resolution for breakpoints and can identify shorted copy number aberrations; large bin size averages out sparsity and leads to less noisy normalized results.

**Figure S19.**
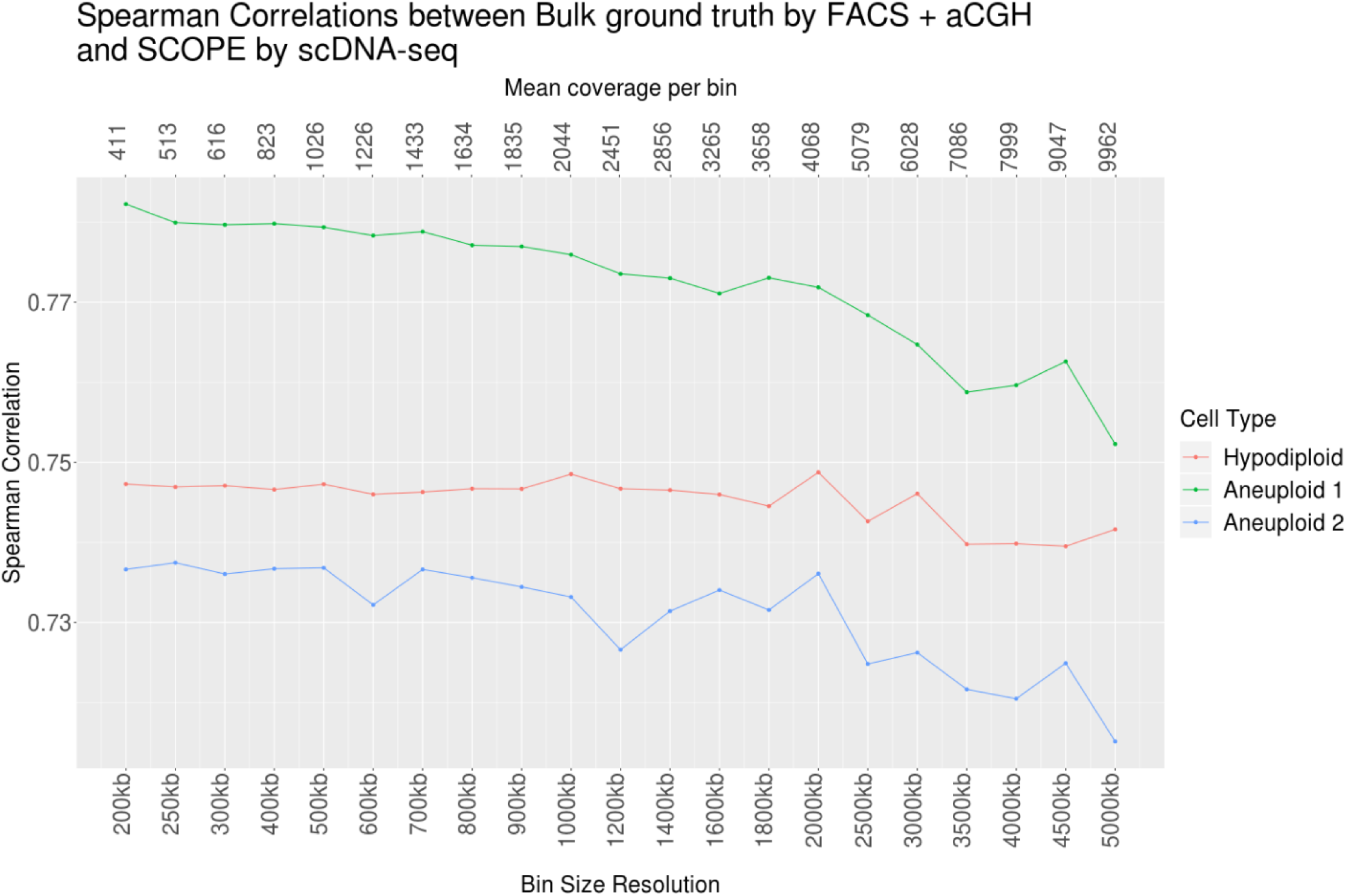
Performance assessment with varying bin size. Larger bin sizes average out sparsity and noise. Copy number estimation by SCOPE is robust to bin size from 200Kb to 1Mb, especially for large copy number events. Further increasing the bin size leads to estimation accuracy loss and thus lower Spearman correlation.

**Figure S20.**
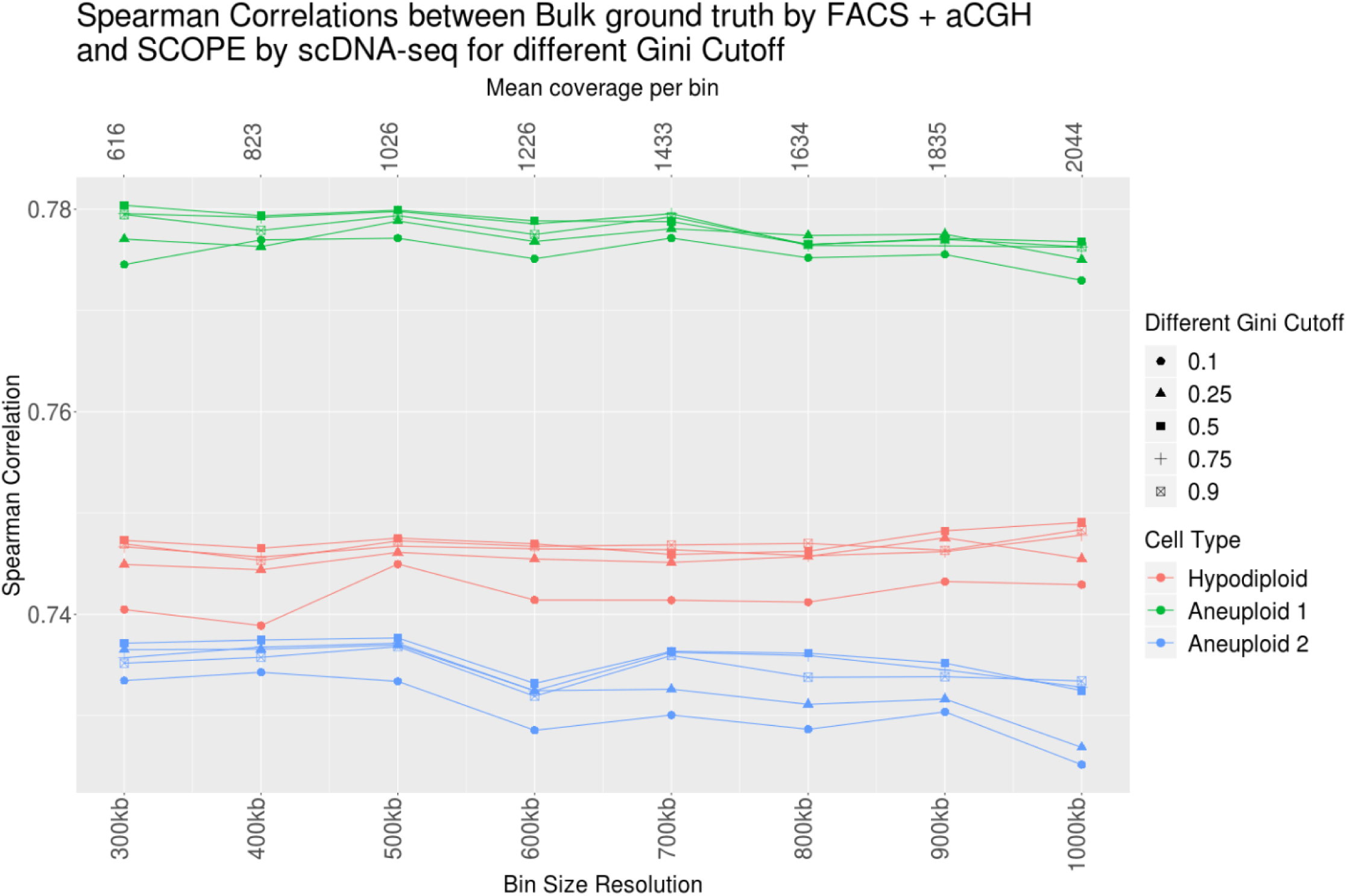
Performance assessment with varying threshold for Gini coefficients. We compare the performance of SCOPE against aCGH gold standards with varying percentiles *q* ∈ {10%, 25%, 50%, 75%, 90%} of Gini indices from true diploid cells as the Gini cutoff, which results in 5, 11, 22, 33, 39 cells on average used as negative controls, respectively. Empirical evidence suggests that SCOPE does not need to include all normal cells as negative controls – 10 to 20 cells suffice to achieve accurate estimation. Different Gini coefficient thresholds do not lead to different performance, even with varying bin size.

**Figure S21.**
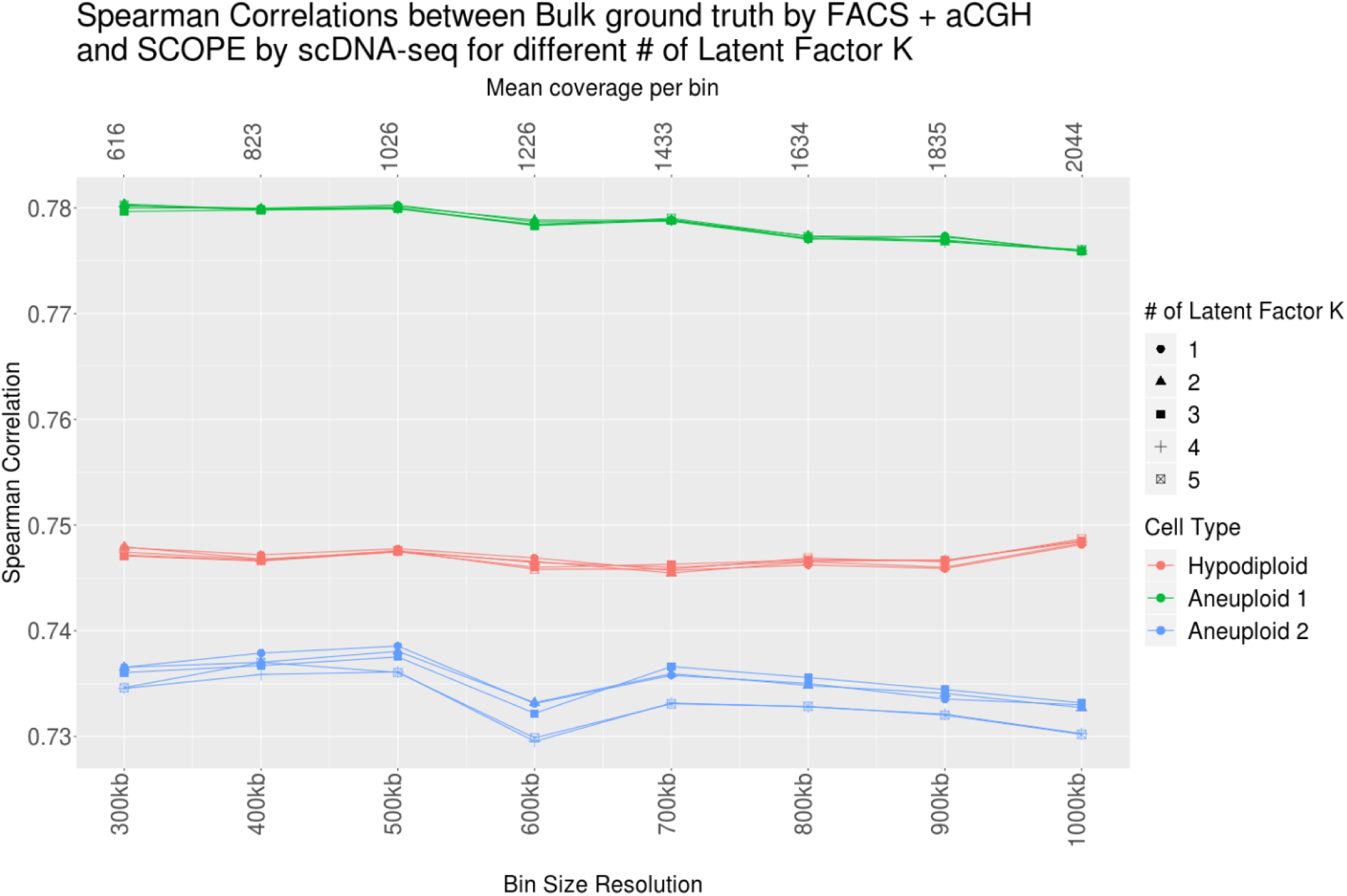
Performance assessment with varying number of Poisson latent factors. SCOPE adopts the EM algorithm to account for different genomic contexts when estimating GC content biases and uses only the normal cells to estimate the bin-specific noise terms. Our results show that SCOPE is robust to the different choice of the number of Poisson latent factors.

## References

1. Sudmant, P.H., Rausch, T., Gardner, E.J., Handsaker, R.E., Abyzov, A., Huddleston, J., Zhang, Y., Ye, K., Jun, G., Fritz, M.H., et al. (2015). An integrated map of structural variation in 2,504 human genomes. Nature 526, 75–81.

2. McCarroll, S.A., and Altshuler, D.M. (2007). Copy-number variation and association studies of human disease. Nat Genet 39, S37–42.

3. Gonzalez, E., Kulkarni, H., Bolivar, H., Mangano, A., Sanchez, R., Catano, G., Nibbs, R.J., Freedman, B.I., Quinones, M.P., Bamshad, M.J., et al. (2005). The influence of CCL3L1 gene-containing segmental duplications on HIV-1/AIDS susceptibility. Science 307, 1434–1440.

4. Pinto, D., Pagnamenta, A.T., Klei, L., Anney, R., Merico, D., Regan, R., Conroy, J., Magalhaes, T.R., Correia, C., Abrahams, B.S., et al. (2010). Functional impact of global rare copy number variation in autism spectrum disorders. Nature 466, 368–372.

5. Marshall, C.R., Howrigan, D.P., Merico, D., Thiruvahindrapuram, B., Wu, W., Greer, D.S., Antaki, D., Shetty, A., Holmans, P.A., Pinto, D., et al. (2017). Contribution of copy number variants to schizophrenia from a genome-wide study of 41,321 subjects. Nat Genet 49, 27–35.

6. Yang, Y., Chung, E.K., Wu, Y.L., Savelli, S.L., Nagaraja, H.N., Zhou, B., Hebert, M., Jones, K.N., Shu, Y., Kitzmiller, K., et al. (2007). Gene copy-number variation and associated polymorphisms of complement component C4 in human systemic lupus erythematosus (SLE): low copy number is a risk factor for and high copy number is a protective factor against SLE susceptibility in European Americans. Am J Hum Genet 80, 1037–1054.

7. Fanciulli, M., Norsworthy, P.J., Petretto, E., Dong, R., Harper, L., Kamesh, L., Heward, J.M., Gough, S.C., de Smith, A., Blakemore, A.I., et al. (2007). FCGR3B copy number variation is associated with susceptibility to systemic, but not organ-specific, autoimmunity. Nat Genet 39, 721–723.

8. Cancer Genome Atlas Research, N., Weinstein, J.N., Collisson, E.A., Mills, G.B., Shaw, K.R., Ozenberger, B.A., Ellrott, K., Shmulevich, I., Sander, C., and Stuart, J.M. (2013). The Cancer Genome Atlas Pan-Cancer analysis project. Nat Genet 45, 1113–1120.

9. International Cancer Genome, C., Hudson, T.J., Anderson, W., Artez, A., Barker, A.D., Bell, C., Bernabe, R.R., Bhan, M.K., Calvo, F., Eerola, I., et al. (2010). International network of cancer genome projects. Nature 464, 993–998.

10. Jiang, Y., Qiu, Y., Minn, A.J., and Zhang, N.R. (2016). Assessing intratumor heterogeneity and tracking longitudinal and spatial clonal evolutionary history by next-generation sequencing. Proc Natl Acad Sci U S A 113, E5528–5537.

11. Urrutia, E., Chen, H., Zhou, Z., Zhang, N.R., and Jiang, Y. (2018). Integrative pipeline for profiling DNA copy number and inferring tumor phylogeny. Bioinformatics 34, 2126–2128.

12. Wellcome Trust Case Control, C., Craddock, N., Hurles, M.E., Cardin, N., Pearson, R.D., Plagnol, V., Robson, S., Vukcevic, D., Barnes, C., Conrad, D.F., et al. (2010). Genome-wide association study of CNVs in 16,000 cases of eight common diseases and 3,000 shared controls. Nature 464, 713–720.

13. Benjamini, Y., and Speed, T.P. (2012). Summarizing and correcting the GC content bias in high-throughput sequencing. Nucleic Acids Res 40, e72.

14. Sims, D., Sudbery, I., Ilott, N.E., Heger, A., and Ponting, C.P. (2014). Sequencing depth and coverage: key considerations in genomic analyses. Nat Rev Genet 15, 121–132.

15. Iakovishina, D., Janoueix-Lerosey, I., Barillot, E., Regnier, M., and Boeva, V. (2016). SV-Bay: structural variant detection in cancer genomes using a Bayesian approach with correction for GC-content and read mappability. Bioinformatics 32, 984–992.

16. Jiang, Y., Oldridge, D.A., Diskin, S.J., and Zhang, N.R. (2015). CODEX: a normalization and copy number variation detection method for whole exome sequencing. Nucleic Acids Res 43, e39.

17. Fromer, M., Moran, J.L., Chambert, K., Banks, E., Bergen, S.E., Ruderfer, D.M., Handsaker, R.E., McCarroll, S.A., O’Donovan, M.C., Owen, M.J., et al. (2012). Discovery and statistical genotyping of copy-number variation from whole-exome sequencing depth. Am J Hum Genet 91, 597–607.

18. Jiang, Y., Wang, R., Urrutia, E., Anastopoulos, I.N., Nathanson, K.L., and Zhang, N.R. (2018). CODEX2: full-spectrum copy number variation detection by high-throughput DNA sequencing. Genome Biol 19, 202.

19. Carter, S.L., Cibulskis, K., Helman, E., McKenna, A., Shen, H., Zack, T., Laird, P.W., Onofrio, R.C., Winckler, W., Weir, B.A., et al. (2012). Absolute quantification of somatic DNA alterations in human cancer. Nat Biotechnol 30, 413–421.

20. Favero, F., Joshi, T., Marquard, A.M., Birkbak, N.J., Krzystanek, M., Li, Q., Szallasi, Z., and Eklund, A.C. (2015). Sequenza: allele-specific copy number and mutation profiles from tumor sequencing data. Ann Oncol 26, 64–70.

21. Chen, H., Jiang, Y., Maxwell, K.N., Nathanson, K.L., and Zhang, N. (2017). Allele-Specific Copy Number Estimation by Whole Exome Sequencing. Ann Appl Stat 11, 1169–1192.

22. Navin, N., Kendall, J., Troge, J., Andrews, P., Rodgers, L., McIndoo, J., Cook, K., Stepansky, A., Levy, D., Esposito, D., et al. (2011). Tumour evolution inferred by single-cell sequencing. Nature 472, 90–94.

23. Wang, Y., Waters, J., Leung, M.L., Unruh, A., Roh, W., Shi, X., Chen, K., Scheet, P., Vattathil, S., Liang, H., et al. (2014). Clonal evolution in breast cancer revealed by single nucleus genome sequencing. Nature 512, 155–160.

24. Kim, C., Gao, R., Sei, E., Brandt, R., Hartman, J., Hatschek, T., Crosetto, N., Foukakis, T., and Navin, N.E. (2018). Chemoresistance Evolution in Triple-Negative Breast Cancer Delineated by Single-Cell Sequencing. Cell 173, 879–893 e813.

25. Liu, J., Adhav, R., and Xu, X. (2017). Current Progresses of Single Cell DNA Sequencing in Breast Cancer Research. Int J Biol Sci 13, 949–960.

26. Dean, F.B., Hosono, S., Fang, L., Wu, X., Faruqi, A.F., Bray-Ward, P., Sun, Z., Zong, Q., Du, Y., Du, J., et al. (2002). Comprehensive human genome amplification using multiple displacement amplification. Proc Natl Acad Sci U S A 99, 5261–5266.

27. Baslan, T., Kendall, J., Rodgers, L., Cox, H., Riggs, M., Stepansky, A., Troge, J., Ravi, K., Esposito, D., Lakshmi, B., et al. (2012). Genome-wide copy number analysis of single cells. Nat Protoc 7, 1024–1041.

28. Zong, C., Lu, S., Chapman, A.R., and Xie, X.S. (2012). Genome-wide detection of single-nucleotide and copy-number variations of a single human cell. Science 338, 1622–1626.

29. Navin, N.E. (2014). Cancer genomics: one cell at a time. Genome Biol 15, 452.

30. Shlien, A., and Malkin, D. (2009). Copy number variations and cancer. Genome Med 1, 62.

31. Wang, X., Chen, H., and Zhang, N.R. (2018). DNA copy number profiling using single-cell sequencing. Brief Bioinform 19, 731–736.

32. Garvin, T., Aboukhalil, R., Kendall, J., Baslan, T., Atwal, G.S., Hicks, J., Wigler, M., and Schatz, M.C. (2015). Interactive analysis and assessment of single-cell copy-number variations. Nat Methods 12, 1058–1060.

33. Li, H., Handsaker, B., Wysoker, A., Fennell, T., Ruan, J., Homer, N., Marth, G., Abecasis, G., Durbin, R., and Genome Project Data Processing, S. (2009). The Sequence Alignment/Map format and SAMtools. Bioinformatics 25, 2078–2079.

34. Laks, E., Zahn, H., Lai, D., McPherson, A., Steif, A., Brimhall, J., Biele, J., Wang, B., Masud, T., and Grewal, D. (2018). Resource: Scalable whole genome sequencing of 40,000 single cells identifies stochastic aneuploidies, genome replication states and clonal repertoires. bioRxiv, 411058.

35. Ha, G., Roth, A., Lai, D., Bashashati, A., Ding, J., Goya, R., Giuliany, R., Rosner, J., Oloumi, A., Shumansky, K., et al. (2012). Integrative analysis of genome-wide loss of heterozygosity and monoallelic expression at nucleotide resolution reveals disrupted pathways in triple-negative breast cancer. Genome Res 22, 1995–2007.

36. Fan, X., Edrisi, M., Navin, N., and Nakhleh, L. (2019). Benchmarking Tools for Copy Number Aberration Detection from Single-cell DNA Sequencing Data. bioRxiv, 696179.

37. Huang, L., Ma, F., Chapman, A., Lu, S., and Xie, X.S. (2015). Single-Cell Whole-Genome Amplification and Sequencing: Methodology and Applications. Annu Rev Genomics Hum Genet 16, 79–102.

38. Risso, D., Ngai, J., Speed, T.P., and Dudoit, S. (2014). Normalization of RNA-seq data using factor analysis of control genes or samples. Nat Biotechnol 32, 896–902.

39. Lee, S., Chugh, P.E., Shen, H., Eberle, R., and Dittmer, D.P. (2013). Poisson factor models with applications to non-normalized microRNA profiling. Bioinformatics 29, 1105–1111.

40. Olshen, A.B., Venkatraman, E.S., Lucito, R., and Wigler, M. (2004). Circular binary segmentation for the analysis of array-based DNA copy number data. Biostatistics 5, 557–572.

41. Shen, J.J., and Zhang, N.R. (2012). Change-Point Model on Nonhomogeneous Poisson Processes with Application in Copy Number Profiling by Next-Generation DNA Sequencing. Annals of Applied Statistics 6, 476–496.

42. Nilsen, G., Liestol, K., Van Loo, P., Moen Vollan, H.K., Eide, M.B., Rueda, O.M., Chin, S.F., Russell, R., Baumbusch, L.O., Caldas, C., et al. (2012). Copynumber: Efficient algorithms for single- and multi-track copy number segmentation. BMC Genomics 13, 591.

43. Anders, S., and Huber, W. (2010). Differential expression analysis for sequence count data. Genome Biol 11, R106.

44. Zhang, N.R., Siegmund, D.O., Ji, H., and Li, J.Z. (2010). Detecting simultaneous changepoints in multiple sequences. Biometrika 97, 631–645.

45. Zhang, N.R., and Siegmund, D.O. (2012). Model Selection for High-Dimensional, Multi-Sequence Change-Point Problems. Stat Sinica 22, 1507–1538.

46. Navin, N., Krasnitz, A., Rodgers, L., Cook, K., Meth, J., Kendall, J., Riggs, M., Eberling, Y., Troge, J., Grubor, V., et al. (2010). Inferring tumor progression from genomic heterogeneity. Genome Res 20, 68–80.

47. Gao, R., Davis, A., McDonald, T.O., Sei, E., Shi, X., Wang, Y., Tsai, P.C., Casasent, A., Waters, J., Zhang, H., et al. (2016). Punctuated copy number evolution and clonal stasis in triple-negative breast cancer. Nat Genet 48, 1119–1130.

48. Patel, A.P., Tirosh, I., Trombetta, J.J., Shalek, A.K., Gillespie, S.M., Wakimoto, H., Cahill, D.P., Nahed, B.V., Curry, W.T., Martuza, R.L., et al. (2014). Single-cell RNA-seq highlights intratumoral heterogeneity in primary glioblastoma. Science 344, 1396–1401.

49. Tirosh, I., Venteicher, A.S., Hebert, C., Escalante, L.E., Patel, A.P., Yizhak, K., Fisher, J.M., Rodman, C., Mount, C., Filbin, M.G., et al. (2016). Single-cell RNA-seq supports a developmental hierarchy in human oligodendroglioma. Nature 539, 309–313.

50. Patro, R., Duggal, G., Love, M.I., Irizarry, R.A., and Kingsford, C. (2017). Salmon provides fast and bias-aware quantification of transcript expression. Nat Methods 14, 417–419.

51. Maaten, L.v.d., and Hinton, G. (2008). Visualizing data using t-SNE. Journal of machine learning research 9, 2579–2605.

52. Fan, J., Lee, H.O., Lee, S., Ryu, D.E., Lee, S., Xue, C., Kim, S.J., Kim, K., Barkas, N., Park, P.J., et al. (2018). Linking transcriptional and genetic tumor heterogeneity through allele analysis of single-cell RNA-seq data. Genome Res 28, 1217–1227.

53. Muller, S., Cho, A., Liu, S.J., Lim, D.A., and Diaz, A. (2018). CONICS integrates scRNA-seq with DNA sequencing to map gene expression to tumor sub-clones. Bioinformatics 34, 3217–3219.

## References

1. Li, H., and Durbin, R. (2010). Fast and accurate long-read alignment with Burrows-Wheeler transform. Bioinformatics 26, 589–595.

2. Li, H., Handsaker, B., Wysoker, A., Fennell, T., Ruan, J., Homer, N., Marth, G., Abecasis, G., Durbin, R., and Genome Project Data Processing, S. (2009). The Sequence Alignment/Map format and SAMtools. Bioinformatics 25, 2078–2079.

3. Navin, N., Kendall, J., Troge, J., Andrews, P., Rodgers, L., McIndoo, J., Cook, K., Stepansky, A., Levy, D., Esposito, D., et al. (2011). Tumour evolution inferred by single-cell sequencing. Nature 472, 90–94.

4. Gao, R., Davis, A., McDonald, T.O., Sei, E., Shi, X., Wang, Y., Tsai, P.C., Casasent, A., Waters, J., Zhang, H., et al. (2016). Punctuated copy number evolution and clonal stasis in triple-negative breast cancer. Nat Genet 48, 1119–1130.

5. Garvin, T., Aboukhalil, R., Kendall, J., Baslan, T., Atwal, G.S., Hicks, J., Wigler, M., and Schatz, M.C. (2015). Interactive analysis and assessment of single-cell copy-number variations. Nat Methods 12, 1058–1060.

6. Derrien, T., Estelle, J., Marco Sola, S., Knowles, D.G., Raineri, E., Guigo, R., and Ribeca, P. (2012). Fast computation and applications of genome mappability. PLoS One 7, e30377.

7. Kim, C., Gao, R., Sei, E., Brandt, R., Hartman, J., Hatschek, T., Crosetto, N., Foukakis, T., and Navin, N.E. (2018). Chemoresistance Evolution in Triple-Negative Breast Cancer Delineated by Single-Cell Sequencing. Cell 173, 879–893 e813.

8. Liu, J., Adhav, R., and Xu, X. (2017). Current Progresses of Single Cell DNA Sequencing in Breast Cancer Research. Int J Biol Sci 13, 949–960.

9. Baslan, T., Kendall, J., Rodgers, L., Cox, H., Riggs, M., Stepansky, A., Troge, J., Ravi, K., Esposito, D., Lakshmi, B., et al. (2012). Genome-wide copy number analysis of single cells. Nat Protoc 7, 1024–1041.

10. Dean, F.B., Hosono, S., Fang, L., Wu, X., Faruqi, A.F., Bray-Ward, P., Sun, Z., Zong, Q., Du, Y., Du, J., et al. (2002). Comprehensive human genome amplification using multiple displacement amplification. Proc Natl Acad Sci U S A 99, 5261–5266.

11. Zong, C., Lu, S., Chapman, A.R., and Xie, X.S. (2012). Genome-wide detection of single-nucleotide and copy-number variations of a single human cell. Science 338, 1622–1626.

12. Navin, N., Krasnitz, A., Rodgers, L., Cook, K., Meth, J., Kendall, J., Riggs, M., Eberling, Y., Troge, J., Grubor, V., et al. (2010). Inferring tumor progression from genomic heterogeneity. Genome Res 20, 68–80.

13. Wang, X., Chen, H., and Zhang, N.R. (2018). DNA copy number profiling using single-cell sequencing. Brief Bioinform 19, 731–736.

14. Patel, A.P., Tirosh, I., Trombetta, J.J., Shalek, A.K., Gillespie, S.M., Wakimoto, H., Cahill, D.P., Nahed, B.V., Curry, W.T., Martuza, R.L., et al. (2014). Single-cell RNA-seq highlights intratumoral heterogeneity in primary glioblastoma. Science 344, 1396–1401.

15. Patro, R., Duggal, G., Love, M.I., Irizarry, R.A., and Kingsford, C. (2017). Salmon provides fast and bias-aware quantification of transcript expression. Nat Methods 14, 417–419.

